# Mapping the rRNA methylome reveals contributions of methyltransferases to ribosome function and antibiotic sensitivity

**DOI:** 10.64898/2026.05.25.727669

**Authors:** Jennie L. Hibma, Kaley M. Simcox, Lia M. Munson, Lauren E. Barnes, Kristin S. Koutmou, Lyle A. Simmons

## Abstract

Ribosomal RNA (rRNA) folds into a complex structure used as the macromolecular core for protein synthesis. Chemical modification of rRNA contributes to ribosome structure, function, and susceptibility to antibiotics. Despite their importance, the enzymes responsible for specific rRNA modifications remain unknown in most species. In this work, we integrate genetics and biochemistry with sequencing and mass spectrometry to uncover enzymes responsible for rRNA methylation events in *Bacillus subtilis*. We characterize 17 enzymes responsible for 20 methylation modifications on the 16S and 23S rRNAs, 11 of which are encoded by previously uncharacterized genes. For each rRNA methyltransferase, we define the modification identity, location, and we determine the impact of loss of cognate rRNA methylation on ribosome biogenesis and antibiotic sensitivity. Our findings demonstrate that loss of nearly half of the 17 genes studied results in alterations to ribosome assembly or antibiotic sensitivity underscoring the importance of chemical modifications to ribosome function.

## Introduction

Ribosomal RNA (rRNA) forms the catalytic core of the translation machinery required for protein production in every cell^1, 2^. The chemical diversity of rRNA nucleotides far exceeds that of the four standard nucleosides, increasing the chemistries and structures available for ribosome assembly and function^3, 4^. Modifications that strategically cluster in crucial functional areas of the ribosome, specifically the peptidyl transferase center (PTC) and the decoding site, ensure efficient and accurate protein synthesis^5, 6^. While several studies show that rRNA modifications are not merely passive, the biological functions of most rRNA modification sites remain to be defined across virtually all biological systems^7, 8^. This problem is further complicated by the fact that each organism possesses a unique collection of rRNA modifying enzymes responsible for its rRNA modification landscape^3^. As such, the investigation of each individual modification site and their cognate enzyme must be pursued one by one, resulting in slow progress in assessing the importance of each modification to ribosome function in most organisms^3, 9^.

Modifications are incorporated into the large and small ribosomal subunits by a host of enzymes acting both co- and post-transcriptionally^10, 11^. Although >170 unique-types of RNA modifications have been identified^3, 9, 12, 13^, rRNA modifications generally fall into three categories: methylation of the ribose sugars at the C2-position, isomerization of uridines to pseudouridines (Ψ), and nitrogenous base modifications **(Figure 1A)**^10^. The loss of rRNA modifications can disrupt multi-step pre-rRNA processing, leading to abnormal translation and negatively impacting cellular fitness^14,15, 16, 17, 18, 19, 20, 21, 22^. Nonetheless, our knowledge about the function of rRNA modifications still comes from a relatively limited number of studies that have examined only individual sites, and as such the consequences of the vast majority of rRNA modifications remain unknown^23, 24, 25, 26^.

**Figure 1.**
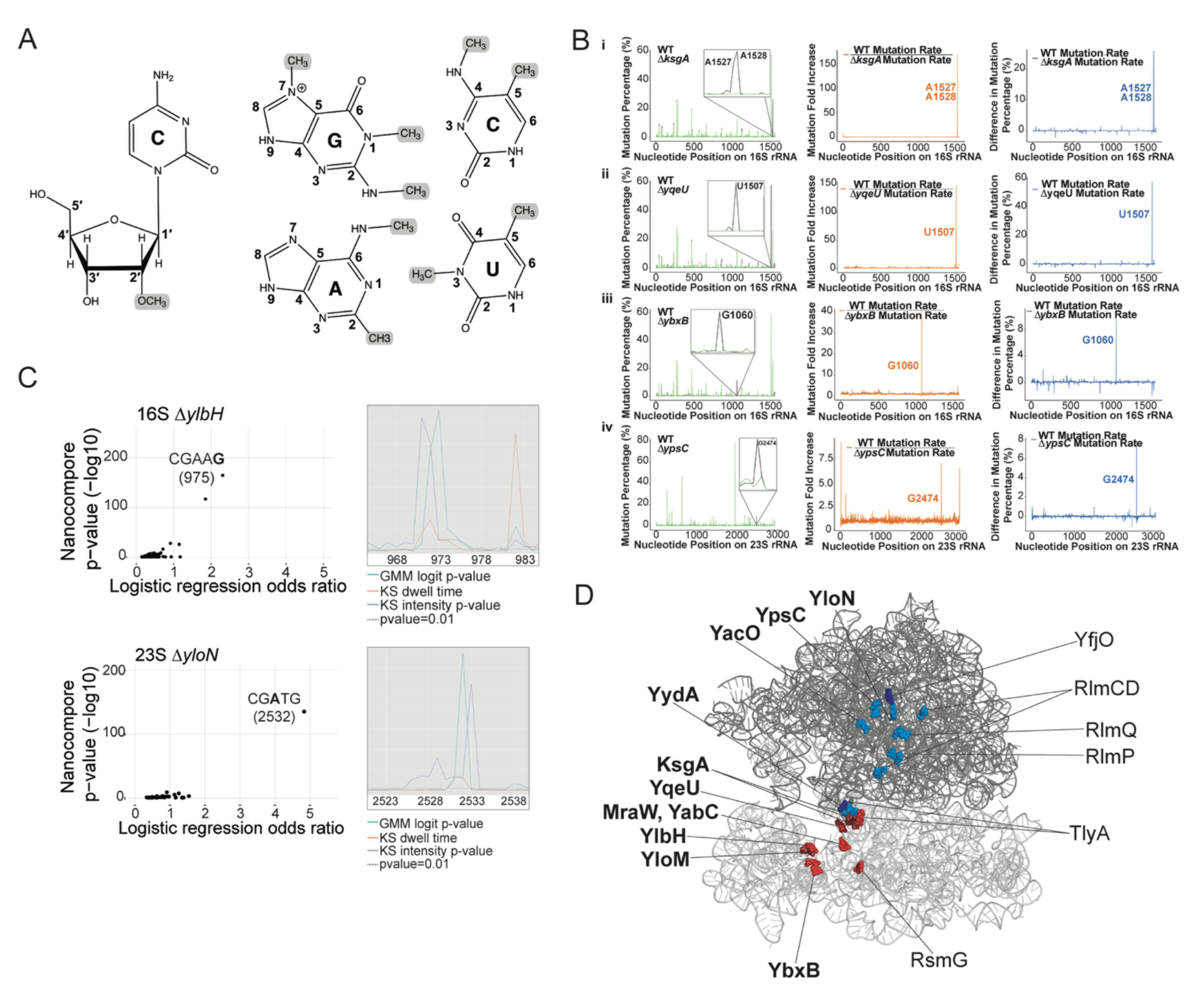
Locations of rRNA methylations in the *B. subtilis* ribosome. **A)** Shown are the positions of relevant RNA methylations. From left to right: 2′-*O*-methylcytidine (Cm); *N*^7^-methylguanine (m^7^G); 1-methylguanine (m^1^G); *N*^2^-methylguanine (m^2^G); *N*^4^-methylcytosine (m^4^C); 5-methylcytosine (m^5^C); *N*^2^-methyladenine (m^2^A); *N*^6^-methyladenine (m^6^A); 3-methyluracil (m^3^U); 5-methyluracil (m^5^U). **B)** The MaP-seq data analysis for the indicated deletion strains. From left to right we show mutation percentage, the fold mutation increase and the difference in mutation percentage used to identify the nucleotide position. **C)** Example of the log odds ratio logistic regression using the nanopore sequencing data (left panels) and the comparison of the ion currents from mutant and WT strains (right panel). This allows for the visualization of modification locations by comparing the sequences of WT to deletion using a logistic regression log odds ratio, kmer dwell time, and kmer electrical signal intensity^53^. Shown are Nanocompore results from Δ*ylbH* 16S rRNA and Δ*yloN* 23S rRNA, revealing modifications at G975 and A2535 respectively. Other nanopore data are shown in Supplemental **Figure S2, S3. D)** Shown is a model summary of the *B. subtilis* ribosome with the position of each methylated nucleotide indicated and the enzyme responsible for the methylation. Bold enzymes have not been previously described.

While there is a conserved core of rRNA modifications across biology^4^ the quantity and identity of modified sites differ markedly between organisms^27, 28^. rRNA modification profiles have been comprehensively mapped in *E. coli, S. cerevisiae* and *H. sapiens* ^9^, and until recently limited data was available about the locations of rRNA modifications outside of these three organisms^8, 29^. However, deep sequencing technologies are beginning to expand our knowledge of rRNA modification distributions across a broader phylogenetic range of organisms. Within the last two years complete maps of rRNA modifications isolated from *B. subtilis* and hyperthermophiles *S. enterica, P. furiosus*, and *T. kodakarensis* have been published^8, 29^. Although modification mapping is extremely valuable, it is only a starting point for understanding the biological function of rRNA modifications. For most organisms, it remains unknown which enzymes are responsible for incorporation of the specific sites of modification. In particular, there is a dearth of knowledge about rRNA modifying enzymes in gram-positive bacteria. Establishing the impact of rRNA modification on ribosome function and antibiotic sensitivity will be essential for this clinically relevant group of gram-positive bacteria.

These significant gaps in our understanding of rRNA modification location, incorporation and function are largely attributed to long-standing technical barriers. In protein post-modification workflows, robust mass spectrometry analyses are used to characterize sites of chemical modification^30, 31, 32^. Analogous workflows for detecting modified oligonucleotides are much more challenging and are typically only performed in a handful of labs with access to specialized technologies^33, 34, 35, 36, 37, 38, 39, 40, 41^. Instead, mass spectrometry is widely employed to quantify nucleoside modifications. Here, RNAs are hydrolyzed to single nucleosides thereby removing the sequence context of modified nucleotides^42^. As a result, for many organisms, we know the overall set of modifications present, but we do not know their precise positions. Recently sequencing-based approaches have started to fill this gap (see reference in ^29^), though these methods are often limited to detecting just one modification at a time. Nanopore direct-RNA sequencing is emerging as a promising platform for simultaneously detecting multiple sites of modification^43, 44^, but it has not yet reached the point where it can reliably generate de novo maps of diverse rRNA modifications without additional chemical information.

Here, we overcome these technical challenges to identify the enzymes responsible for incorporating methylations into *B. subtilis* rRNA by implementing orthogonal modification mapping approaches. We further show that several rRNA methylation modifications are essential for ribosome biogenesis and proper ribosome function. By integrating genetics, sequencing, mass spectrometry, and biochemical methods, we reveal the impact of 20 rRNA modifications in this important gram-positive model organism which diverged from *E. coli* over two billion years ago^45, 46, 47^. Our work not only provides new insight into RNA modification identification and functionality, but we also demonstrate how a combined workflow can advance our understanding of RNA biology.

## Results

### Identifying the enzyme-specific locations of rRNA methylations

To identify putative rRNA methyltransferase (MTase) genes, we performed homology searches of the *B. subtilis* PY79 sequenced genome^48^ for sequence similarity to known bacterial MTase genes primarily from *E. coli* since all the enzymes in this species were comprehensively studied^7^ **(Table S1)**. From this search we establish a list of 26 total rRNA MTase genes of which six had been previously studied and 20 remained as candidate rRNA methyltransferases. Individual gene deletions were constructed for all 26 candidates to enable the investigation of their potential rRNA methylation activities **(Table S2)**. 16S and 23S rRNA were isolated from each putative rRNA MTase deletion strain, as well as wild-type (WT) *B. subtilis*. To assess if (and how) each putative rRNA MTase deletion strain altered the modification profile of the rRNAs, the isolated rRNAs were subjected to Mutational-Profiling and Sequencing (MaP-Seq) or direct RNA sequencing by Oxford Nanopore Technologies (DRS)^49, 50, 51^. Both experimental approaches have advantages and disadvantages for detecting particular modifications. These two approaches allow us to identify rRNA sites that exhibit changes in their modification status between the putative MTase deletion cells and WT control.

MaP-seq was performed for deletion strains that had a putative modification in the Watson-Crick face, namely 1-methylguanosine (m^1^G), *N*^2^-methylguanosine (m^2^G), *N*^6^-methyladenosine (m^6^A), 2-methyladenosine (m^2^A), *N*^4^-methylcytidine (m^4^C), or 3-methyluridine (m^3^U) based on homology to known *E. coli* enzymes **(Table S1, Figure 1B)**. In this workflow, rRNAs are reverse transcribed with the highly sensitive reverse transcriptase SuperScript II and the positions modified induce increased levels of dNTP misincorporation. This sensitivity to modification has made MaP-seq an excellent tool for probing RNA secondary structure following the addition of structure-sensitive nucleotide-modifying reagents, such as DMS as part of the method SHAPE ^49, 52^. We used MaP-seq to compare the modification profiles of rRNAs isolated from WT and putative methyltransferase strains. These analyses revealed the loss of methyl-induced mutations in four putative methyltransferase strains, allowing us to assign biological roles to four previously uncharacterized genes. **(Figure 1B, left)**. Mutation rate ratios and differences were calculated at each nucleotide and highlight the contributions of the enzymes encoded by *ksgA, yqeU*, and *ybxB* on the 16S rRNA and *ypsC* on the 23S rRNA **(Figure 1B)**. Since there are a host of mutations profiled across the rRNA to varying degrees **(Figure 1B, left)**, likely due to rDNA operon variants or other RNA modifications, both ratio and difference calculations were used to detect single nucleotide positions that significantly differed between strains. Thus, MaP-seq effectively identified native rRNA methylation sites in the Watson-Crick face defining the function of four previously uncharacterized genes we now define as rRNA methyltransferases **(Table 1, Figure 1B)**.

**Table 1.**
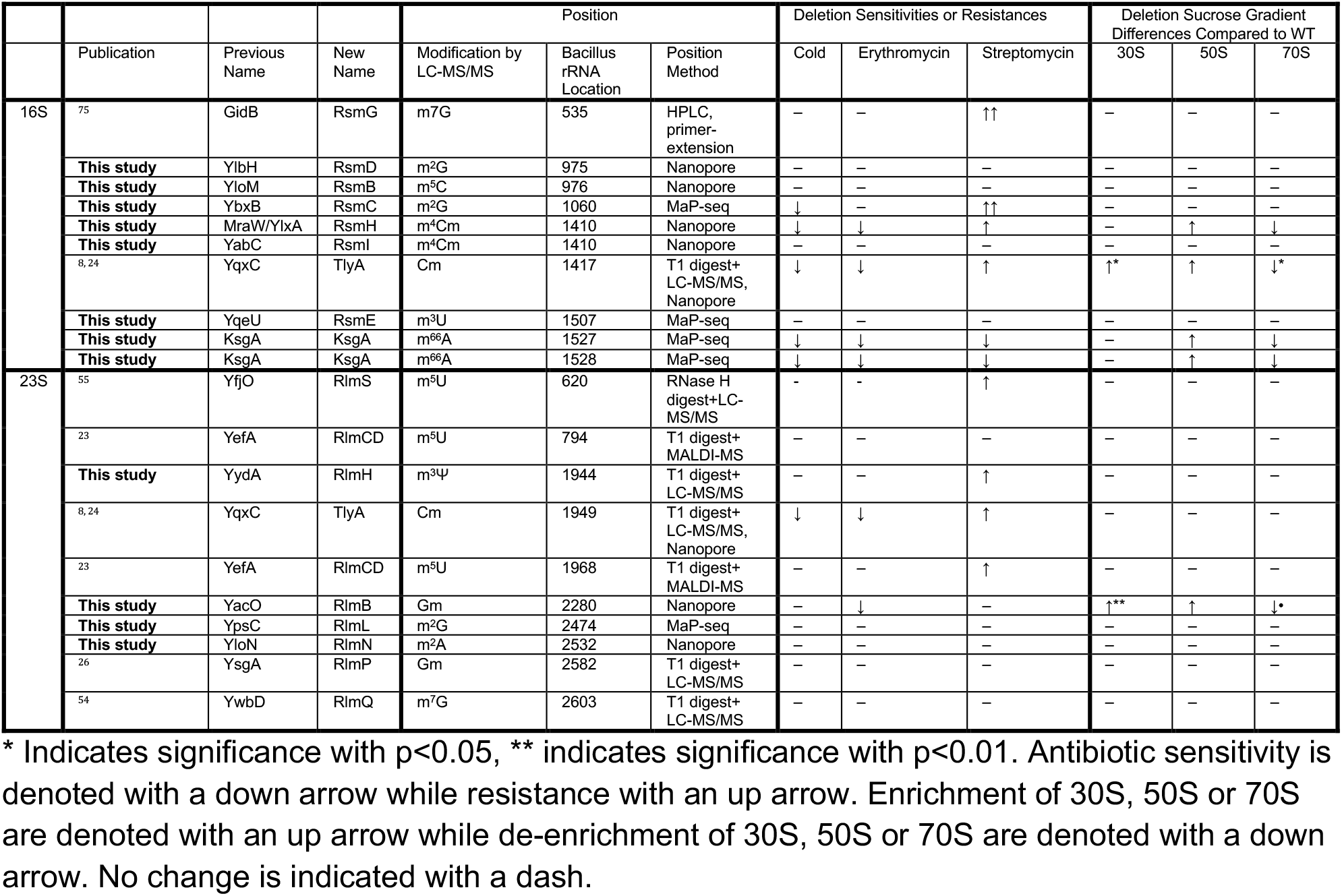
Summary of *B. subtilis* rRNA modification, MTase gene, deletion phenotype and methods used.

Given that MaP-seq is most effective for modifications predicted to be in the Watson-Crick face, we used Oxford Nanopore direct RNA sequencing (DRS) to map the remaining unknown methylation sites. This included DRS in six putative methyltransferase deletion strains where MaP-seq was not successful **(Figure S1)**. In nanopore DRS, modified and unmodified bases produce distinct electrical signals as RNA molecules pass through the nanopore. To identify sites where modifications are lost in deletion strains, raw ionic current data from WT and deletion strains were compared using Nanocompore analysis^53^. Comprehensive results for all nanopore sequencing showing the Gaussian mixture model (GMM) logit p-value, dwell times, and intensities along the 16S or 23S molecules are provided in **Figure S2 and S3**. This approach enabled us to identify four 16S enzymes (YlbH, YloM, MraW, and YabC), and two 23S enzymes YloN and YacO that modify rRNA **(Figure S2 and S3)**. We were unable to confidently assign two putative enzymes, YfjO and YydA, but discuss their writer roles below.

Together, use of MaP-seq and Nanopore (DRS) discovered 10 new *B. subtilis* rRNA MTases and identified the sites they modify. It is important to note that while these sequencing data confidently assign the positions modified by the rRNA modifying enzymes we found, they do not provide insight into the chemical identity of the modification they incorporate. With the data presented here and that follows, we provide a summary of the modification locations mapped on the rRNA structure for all 20 modifications made by their 17 respective enzymes. 19 modifications by 16 enzymes were identified or verified in this work including 11 previously undefined genes (bold) along with six known methyltransferases^8, 23, 24, 26, 54, 55^ **(Table 1, Figure 1A)**. This structural mapping reveals that, like other organisms, the sites of modification are largely clustered around the ribosome active site.

### Identification of chemical modifications by LC-MS/MS

To complement our modification sequencing-based mapping assays, we used nucleoside liquid chromatography-tandem mass spectrometry (LC-MS/MS) to identify the modifications introduced by candidate MTases into 16S and 23S rRNAs purified from *B. subtilis*. LC-MS/MS provides orthogonal information regarding the chemical identity of the nucleosides, something that cannot be obtained from the modification-mapping approaches alone. The levels of 58 possible canonical and modified nucleosides were assessed in 16S and 23S rRNA samples purified from WT cells and from the 26 methyltransferase candidate knockout cell lines **(Table S2, Supplemental Figures S4-S16)**^56^

In 16S rRNA samples isolated from WT cells, we detected nine modified nucleosides in addition to the four canonical bases: dihydrouridine (D), pseudouridine (Ψ), 5-methylcytidine (m^5^C), *N*^7^-methylguanosine (m^7^G), 2′-*O*-methylcytidine (Cm), *N*^4^,2′-O-dimethylcytidine (m^4^Cm), 3-methyluridine (m^3^U), *N*^2^-methylguanosine (m^2^G), and N6,N6-dimethyladenosine (m^6,6^A). After calculating the pM abundances of modified nucleosides, the number of modifications per strain was calculated using the modification/main base ratio ^24, 56^ and the number of each canonical base in the unmodified RNA transcript. The enzymes responsible for all seven of the 16S methylated nucleosides were established by comparing the difference in levels of modified nucleosides from the WT RNA and the putative MTase knockout strains we generated **(Figure 2A)**. The enzyme and modified nucleoside assignments are as follows: YloM (m^5^C), RsmG (m^7^G), TlyA (Cm), MraW (m^4^Cm), YqeU (m^3^U), YbxB (m^2^G), YlbH (m^2^G), and KsgA (m^6,6^A) **(Figure 2, Figure S16, Figure S17)**. Our findings corroborate previous reports demonstrating that RsmG and TlyA incorporate m^7^G and Cm, respectively, into 16S *B. subtilis* rRNA (**Figures S16-S17)**^8, 24, 25^. Comparison of extracted ion chromatograms (EICs) demonstrate that m^7^G and m^3^U are incorporated at substoichiometric levels (< 0.5 mod per strain) by RsmG and YqeU **(Figure S17)**. While most types of modification are incorporated by a single enzyme, some (e.g. m^2^G, m^4^Cm) are installed by multiple RNA modifiers **(Figure 2A)**. For example, the gene products of *ybxB* and *ylbH* incorporate ~1 m^2^G each per 16S rRNA, while m^4^Cm is added by *mraW* and *yabC* **(Figure 2, Figure S16, S17)**. We observed no cross-talk between any enzymes, as loss of any single putative MTase did not perturb the nucleoside abundances of other 16S rRNA modifications - as determined by the combined analysis of MaP-seq, nanopore DRS and nucleoside LC-MS/MS abundances **(Figure 2, Figure S1-S3, S16-S17)**.

**Figure 2.**
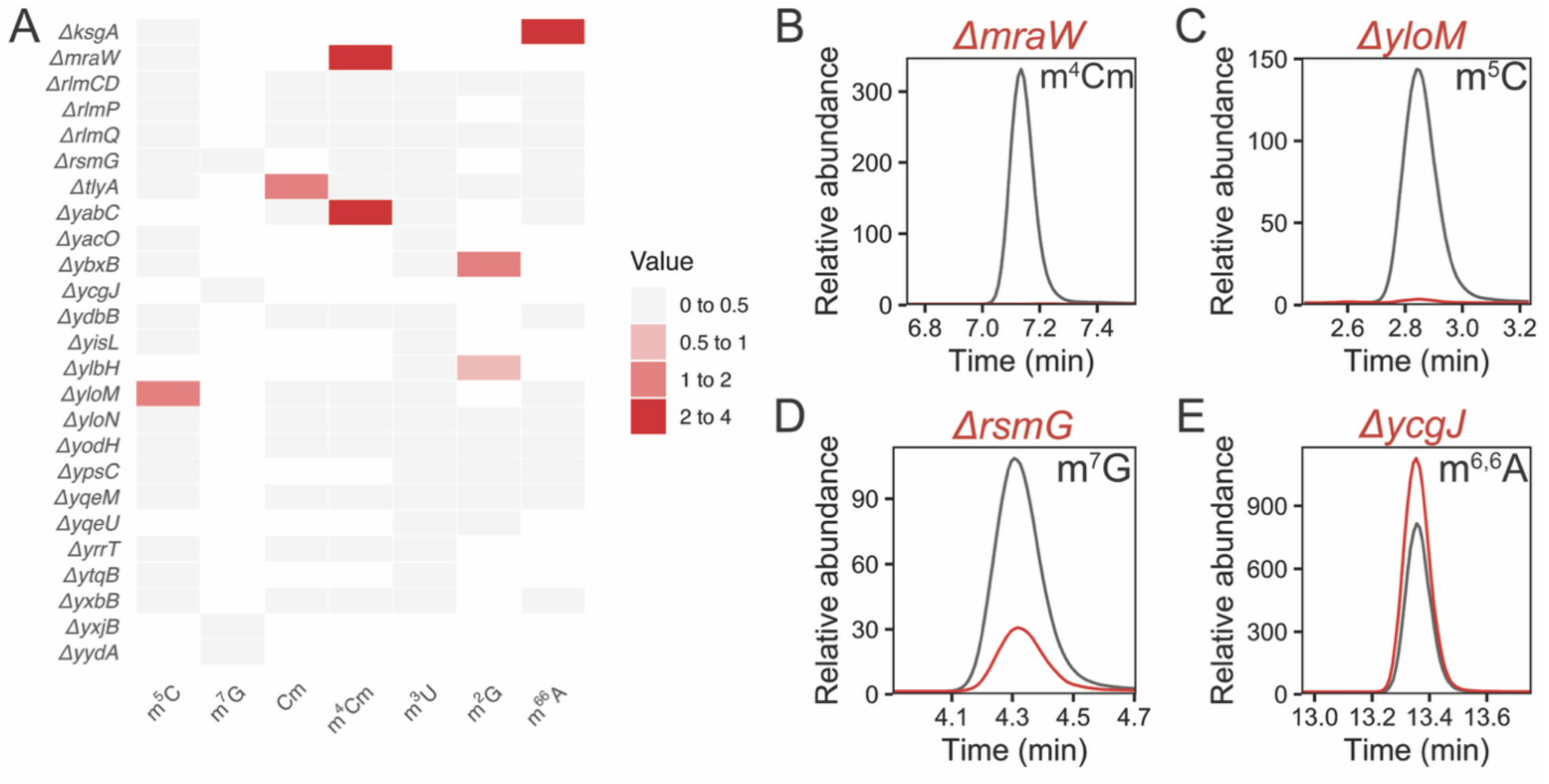
LC-MS/MS analysis of *B. subtilis* 16S rRNA. **A)** Difference heatmap of the average number of methylated nucleosides from WT and mutant strains. Data were filtered to remove strains where the number of methylated nucleosides is increased compared to wildtype. These are represented by white boxes in the heatmap. Knockout strains that change modification levels sub-stoichiometrically (< 0.5 mod per strain), but still reduced modification levels in extracted ion chromatograms include: Δ*rsmG* (m^7^G) and Δ*yqeU* (m^3^U). **B-D)** Extracted ion chromatograms (EICs) of putative methyltransferase knockout strains that were found to incorporate a 16S methylated nucleoside. WT rRNA nucleosides are shown in grey and mutant rRNA nucleosides are shown in red. **E)** An example EIC of a putative methyltransferase knockout strain that did not incorporate a 16S methylated nucleoside. EICs of the 16S methylated nucleosides for assigned enzymes and all strains profiled are in **Figures S16** and **Figure S17**, respectively.

The modification profile of 23S *B. subtilis* rRNA also includes nine modified nucleosides: D, Ψ, m^7^G, Cm, 3-methylpseudouridine (m^3^Ψ), 5-methyluridine (m^5^U), *N*^2^-methyladenosine (m^2^A), 2′-*O*-methylguanosine (Gm), and m^2^G. As with the 16S, the enzymes responsible for incorporating these modifications were identified by visualizing the difference in number of modifications between WT and mutant strains **(Figure 3A)**. Our results recapitulate prior findings showing that YefA (*rlmCD*), TlyA, YsgA,YwbD (RlmQ) incorporate m^5^U, Cm, Gm, and m^7^G respectively ^8, 23, 24, 26, 54^. Furthermore, we were able to assign four putative methyltransferases to the modification they incorporate in the 23S. These include: YacO (Gm), YpsC (m^2^G), YydA (m^3^Ψ), and YloN (m^2^A) **(Figure 3, Figure S16 and S18)**. Like the 16S, some modifications (e.g. Gm and m^2^A) are incorporated by two enzymes **(Figure 3A)**. For example, YacO and YsgA both add Gm at different positions. While YsgA was previously known to modify one guanosine residue to Gm^26^, we have identified an additional enzyme, YacO, which installs ~1 Gm on the 23S rRNA (**Figure S18, panel C)**. LC-MS/MS abundances of m^2^A decrease in Δ*ybxB* compared to WT, suggesting that *ybxB* (23S) incorporates m^2^A. Nonetheless, our nanopore DRS data of rRNA from these deletions does not detect loss of modification within the *B. subtilis* 23S rRNA for these strains. Therefore, we are unable to confidently assign a catalytic function to YbxB in 23S rRNA methylation.

**Figure 3.**
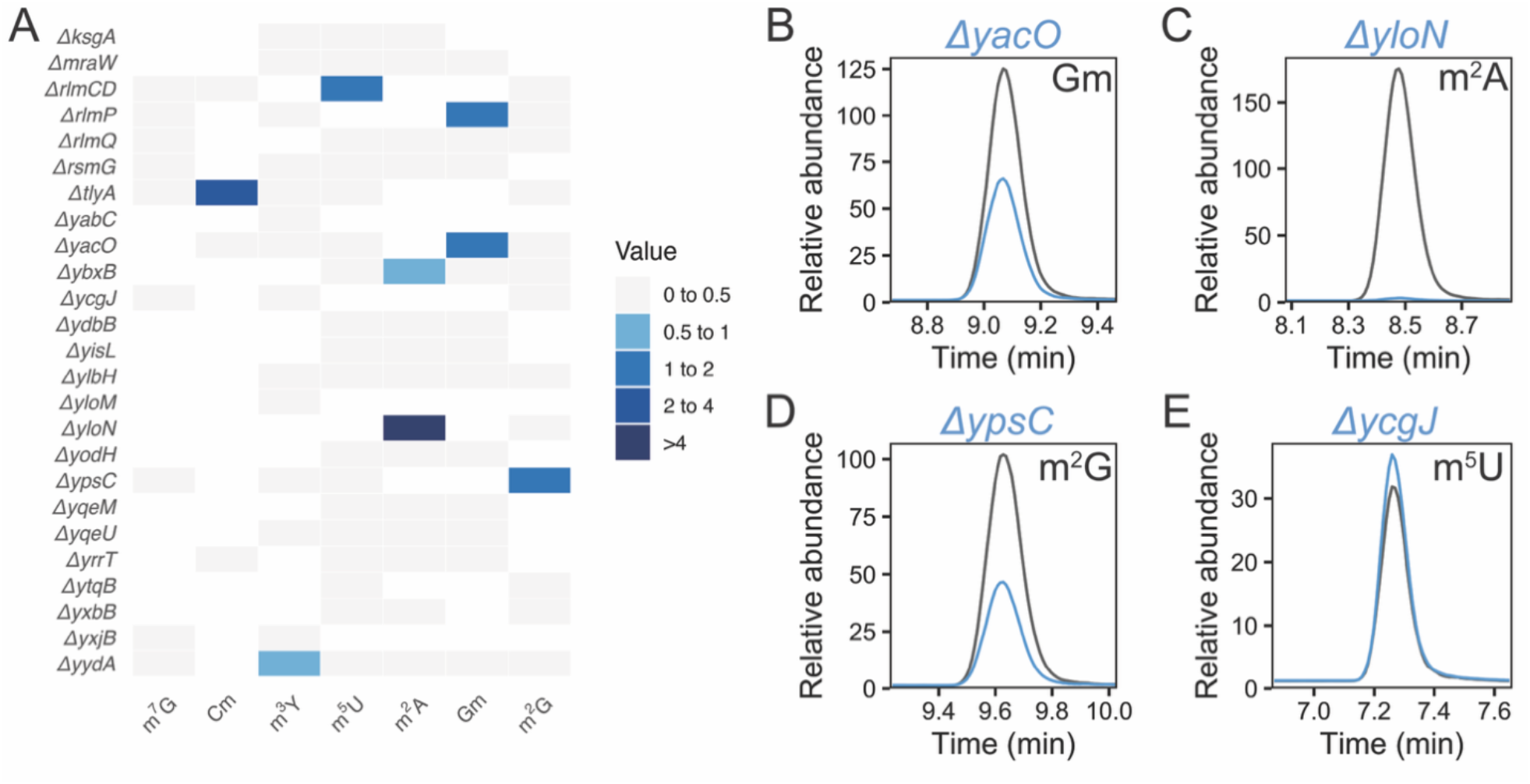
LC-MS/MS analysis of *B. subtilis* 23S rRNA. **A)** Difference heatmap of the average number of methylated nucleosides from WT and mutant strains. Data were filtered to remove strains where the number of methylated nucleosides is increased compared to WT. These are represented by white boxes in the heatmap. **B-D)** Extracted ion chromatograms (EICs) of putative methyltransferase knockout strains that were found to incorporate a 23S methylated nucleoside. WT rRNA nucleosides are shown in grey and mutant rRNA nucleosides are shown in blue. The Δ*rlmQ* (m^7^G) changes modification levels sub-stoichiometrically (< 0.5 mod per strain) but still reduces modification levels in extracted ion chromatograms. **E)** An example EIC of a putative methyltransferase knockout strain that did not incorporate a 23S methylated nucleoside. EICs of the 23S methylated nucleosides for assigned enzymes and all strains profiled are in **Figures S16** and **Figure S18**, respectively.

In addition to identifying the modifications that four candidate *B. subtilis* MTases incorporate, we also determine that the remaining putative methyltransferases are not modifying the 23S rRNA through the combination of nucleoside LC-MS/MS and nanopore sequencing. The only modification we were unable to assign to a putative methyltransferase was m^5^U. Here, we have a clear signal that YefA (RlmCD) incorporates ~1 m^5^U per 23S rRNA; however, there is a remaining signal which corresponds to an additional ~1 m^5^U. Recent studies demonstrate that YfjO is the enzyme that incorporates this additional m^5^U^55^. In total, through our work we can assign 16 enzymes installing 19 modifications. With a recent report defining YfjO as RlmS^55^ this brings the total rRNA methylation modifications to 17 enzymes installing 20 modifications in *B. subtilis* providing the first complete picture of RNA MTases in a gram-positive bacterium.

### Antibiotic sensitivity modulated in rRNA methyltransferase deletion strains

Given that we now have the full complement of rRNA methylating enzymes and their corresponding modifications **(Figure 1, Table 1)**, we asked if any rRNA MTase deletion strains confer phenotypic differences compared with WT. We performed spot titer growth assays in response to a variety of stressors. Because ribosome assembly defects in *B. subtilis* and other bacteria have been shown to be exacerbated by cold temperatures^24, 57^, single deletion strains were grown at 25°C. We found reduced growth of Δ*tlyA*, Δ*ybxB, ΔmraW*, and Δ*ksgA* strains **(Figure 4A+B, S19)** compared to control growth at 37°C. Additionally, deletion strains were grown on plates containing two ribosome targeting antibiotics: erythromycin, a macrolide targeting the peptide exit tunnel of the 23S rRNA, and streptomycin, an aminoglycoside targeting the 16S rRNA near the ribosomal decoding site^58, 59^. Strains with Δ*tlyA*, Δ*mraW, ΔksgA, ΔyacO::kan*, and Δ*rlmP* exhibit reduced growth on erythromycin as compared to WT **(Figure 4A+B, S19)**. Strains with single deletions of *ΔtlyA, ΔrsmG and ΔybxB* were resistant to streptomycin showing increased growth relative to WT, while strains with *ΔyqeU, ΔksgA, ΔyacO:kan* and *ΔyloN* showed decreased growth **(Figure 4A+B, S19)**. In addition, we were able to complement the phenotypic effects of the deletion strains by ectopic expression of the appropriate gene in their respective deletion background **(Figure S20)**. These results demonstrate that loss of several rRNA MTase genes causes striking antibiotic sensitivity phenotypes underscoring the importance of rRNA methylation for proper ribosome function in *B. subtilis*.

**Figure 4.**
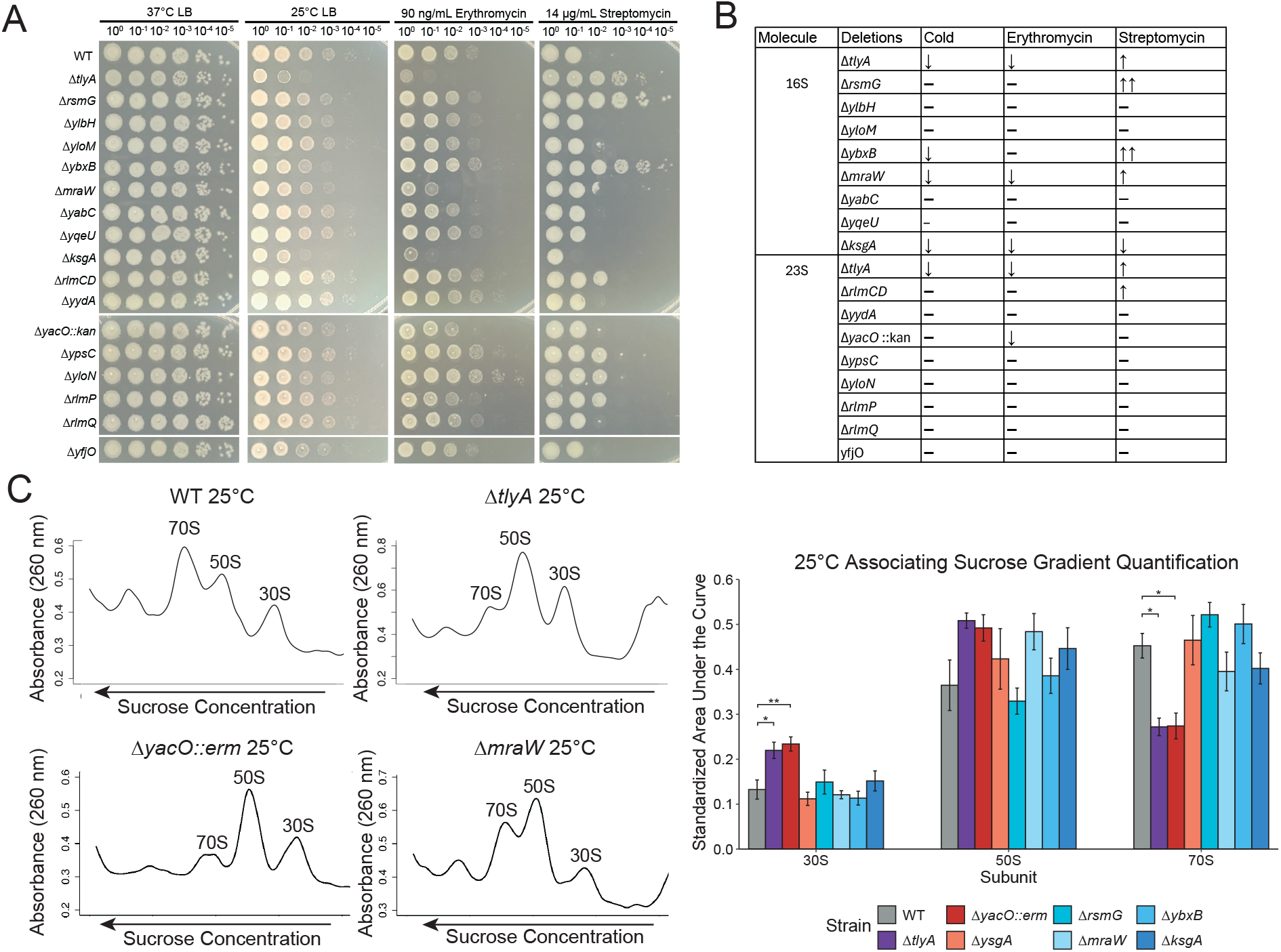
Loss of rRNA methylation impacts antibiotic efficacy and 70S ribosome assembly. **A)** Shown are spot plates with serial dilutions of the indicated strains with the stressors listed. All strain spot titers shown are from the same 150mm large diameter LB agar dish with the indicated condition. Strains Δ*yacO* through Δ*yfjO* were spotted on a different section of the corresponding petri dish. **B)** A table summarizing the phenotypic results. **C)** Example sucrose gradients with rRNA MTase deletions that showed an effect on ribosome assembly with the area under the curve quantified in the right panel. Spot plates and sucrose gradients for the other strains are shown in Supplementary **Figures S19-S23**. The asterisk indicates values are significant with p<0.05, double asterisk indicates values are significant with p<0.01.

### Ribosome assembly intermediates accumulate in rRNA methyltransferase deletion strains

To determine if any individual modification impacts ribosome assembly, we used sucrose gradient ultracentrifugation at 37°C and 25°C for each rRNA MTase deletion mutants that showed a distinct growth defect compared to wild type in the spot titer assays under any condition tested **(Figure S22-S24)**. Sucrose gradients were performed using associating conditions where the full 70S ribosome is formed or dissociating conditions where the ribosomal subunits remain separate as the large 50S subunit and the small 30S subunit. Our results show significant increases of the 30S subunit in Δ*tlyA* and Δ*yacO::erm* strains compared to WT. These strains also showed increases in 50S subunits and decreases in mature 70S ribosomes suggesting rRNA modifications from *tlyA* and *yacO* are important for full ribosome assembly. In the absence of modifications from *tlyA* and *yacO* the small and large subunits do not assemble efficiently **(Figure 4C)**. Moderate increases in the 50S subunit and decreases in the 70S subunit were also observed in Δ*mraW* and Δ*ksgA* gradients **(Figure 4C)**. Other deletions including Δ*ysgA*, Δ*rsmG*, and Δ*yxbB* showed minimal differences when compared to WT **(Figure S21-S24)**. Therefore, our results show that loss of several rRNA MTase genes results in changes to antibiotic challenge and assembly of mature ribosomes. Since rRNA MTase genes from *B. subtilis* are highly conserved in many gram-positive pathogens **(Figure S25)**, our results underscore the importance of understanding how several rRNA MTase gene deletions could alter antibiotic susceptibility or resistance in clinically relevant settings.

## Discussion

Across biology, the catalytic RNA components of the molecular machine responsible for protein synthesis, the ribosome, contain enzymatically modified nucleosides^1, 3, 60^. Although many common rRNA modifications have long been chemically characterized, it has remained challenging to move beyond cataloging their existence to determining where they occur, which enzymes install them, and what molecular functions they perform. In this work, we combine genetic, analytical chemistry, sequencing, and phenotypic characterization to provide the first comprehensive identification and functional characterization of the methyltransferases (MTases) that install rRNA methylations in a gram-positive bacterium. In total, we directly characterize 16 enzymes that install 19 modifications, and we assign biological functions to 11 previously uncharacterized genes in *B. subtilis*. With RlmS^55^, this brings the total rRNA methylations in *B. subtilis* to 17 enzymes installing 20 modifications. Importantly, we find that deletion of several MTase genes resulted in alterations in ribosome maturation and/or antibiotic challenge. Since several of the genes we identify here are conserved in many gram-positive pathogens **(Figure S25)** and differ in their function from *E. coli*, we expect that our results provide new insight into potential gene targets for novel therapeutics or gene inactivating mutations that could result in significant drug resistance in clinical settings. Therefore, our modification identification, mapping and phenotypic characterization provides new avenues for contending with bacterial infections from gram-positive bacterial pathogens harboring conserved MTase genes.

The first goal of our study was to identify the genes that methylate *B. subtilis* rRNA. Of the 26 putative rRNA genes we identified and characterized, 17 are responsible for installing 20 rRNA methylations. 11 of these 17 enzymes are new to this study with five of the other six verified here, but also described previously^8, 23, 24, 26, 54, 55^. The remaining 8 putative enzymes we investigated are not responsible for rRNA methylation but still have the potential to be involved in tRNA or mRNA methylation. Although DNA can also be modified in cells, the *B. subtilis* enzymes incorporating m^6^A and m^5^C have already been identified^61, 62^, and we therefore do not anticipate that any of the unassigned MTase enzymes investigated here act upon DNA.

We identified 19 rRNA modification sites using a combination of LC-MS/MS, nanopore DRS, and MaP-seq. No single technique was optimal for detecting all modifications across all sites. Nucleoside LC-MS/MS provided critical insight into which enzymes install specific classes of modifications, but it cannot determine where those modifications occur. This limitation is especially important when trying to distinguish modifications installed by multiple enzymes. In contrast, both MaP-seq and nanopore DRS can provide positional information, but they generally do not reveal the chemical identity of the modification. We found that MaP-seq was most effective for detecting methylations located on the Watson-Crick face of the corresponding nucleobase, such as m^6,6^A, m^3^U and m^2^G (**Figure 1**). Nanopore DRS was more robust for detecting a range of modified sites, but it still did not produce universally clear results. This highlights the challenge of using nanopore DRS to map modifications *de novo*, as opposed to investigating established modification sites. For some samples, differences between WT and rRNA MTase deletion strains were evident only in electrical signal intensity, and not in additional measures such as logits or dwell time (e.g., *mraW*). This may reflect differences in the ability of the nanopore to detect and distinguish certain modified bases from their canonical counterparts. Alternatively, some modifications may be present at substoichiometric levels in the cell, producing only subtle differences between WT and deletion mutants. Modification mapping by nanopore DRS is also complicated by the fact that the modified position can generally only be narrowed to a five-nucleotide kmer unless the modification identity is already known. Moreover, the nanopore analysis used here was unable to detect all nucleobase methylations. For example, our LC-MS/MS results show that Δ*yydA* loses an m^3^Ψ in 23S rRNA, but the corresponding nanopore electrical signal peak does not match prior reports placing m^3^Ψ at U1944 in *B. subtilis* 23S rRNA, or at the analogous position in homologous *E. coli* 23S rRNA^8, 63, 64^. Similarly, prior work found a m^5^U methylation at 23S U620, and our work shows Δ*rlmCD* loses only one of two m^5^U methylations^23^. Previous work as well as our own blast analysis **(Table S1)** show YfjO shares high sequence identity with m^5^U modifying enzyme RlmCD, suggesting it may have a similar function^23^. In line with this, a recent report used LC-MS/MS to verify that this is in fact the case^55^. However, the noise of the nanopore traces for RNAs isolated from *yjfO* (*rlmS*) knockout were inconclusive though one peak did align to the now characterized position **(Figure S3C)**. Together, these findings highlight the importance of integrating LC-MS/MS with more than one sequencing-based approach, as well as selecting the most appropriate sequencing method, to confidently map modification sites.

An important challenge in rRNA modification biology is determining how (and if) individual modifications impact ribosome assembly and/or function. We find strong evidence for the importance of several methylations by analysis of the MTase deletion phenotypes. The spot plate growth assays show clear phenotypes with specific MTase deletions. Sensitivity to cold indicates a potential ribosome assembly defect^24, 57^. Particular deletion strains sensitive to cold, notably Δ*tlyA* and Δ*ksgA*, show increased 30S and 50S subunit accumulation and reduced mature 70S formation. This is likely due to the fact that these two enzymes confer two modifications instead of one. Loss of *ksgA* has previously been shown to accumulate the 16S precursor 17S rRNA^57^. The assembly defect is also likely influenced by their location as all four modifications are located near the interface of the 23S and 16S subunits. We suggest that loss of methylation from TlyA and KsgA act by destabilizing the ribosome during stress conditions. Importantly, Δ*yacO* also showed significant changes in subunit accumulation with decreases in mature 70S formation compared to WT. We did not identify an MTase deletion strain that showed resistance to erythromycin suggesting loss of rRNA modification does not prevent erythromycin from binding the ribosome. This is consistent with bacterial erythromycin resistance mechanisms. For example, in *S. aureus* gain, not loss, of rRNA modifications by inducing expression of the plasmid derived methyltransferase ErmC causes erythromycin resistance^65^. Strains showing the greatest sensitivities to erythromycin included Δ*tlyA*, Δ*mraW*, Δ*ksgA*, and Δ*yacO*, which also exhibited differences in ribosome assembly suggesting ribosome assembly defects may generally impact cell growth which is further exacerbated by erythromycin blocking polypeptide formation ^59^. The aminoglycoside streptomycin binds to the interface of the rRNAs at helix 69 of the 23S and helix 44 of the 16S interfering with the ribosome decoding center ^66, 67^. The three streptomycin resistant deletion strains Δ*tlyA*, Δ*rsmG, and* Δ*ybxB* have modifications located on helices that constitute the A- and P-sites of the ribosome decoding center at helix 44, 31, and 34, respectively. Therefore, loss of these modifications may change the local structure preventing streptomycin binding and allowing for resistance.

Gram-positive and gram-negative eubacteria evolved approximately two billion years ago^45, 46, 47^. Although many rRNA methyltransferases remain conserved, our work reveals important differences between gram-negative and gram-positive bacteria. In *E. coli* 23 rRNA MTases confer 24 modifications^7^, while in *B. subtilis* 17 enzymes confer 20 modifications **(Table 1, Figure 1)**. These two species share 14 conserved methylation modifications with *E. coli* containing 9 and *B. subtilis* containing 5 unique modifications, respectively^7^. Though not all modifications are conserved, many are clustered in functional areas of the ribosome including the peptidyl transfer center, the decoding center containing the A- and P-tRNA binding sites and inter-subunit bridges^7, 68^. In *B. subtilis* TlyA introduces one modification on both the 16S and the 23S rRNA, causing defects in ribosome assembly and antibiotic sensitivity^24^. *E. coli* lacks TlyA but has RlmN which is unique from *B. subtilis* and it modifies both the 23S and a tRNA^69^. Only five methylation modifications are universally conserved between *E. coli*, yeast, and humans. These include the modifications made by 16S modifying prokaryotic enzymes RsmH and KsgA, which makes two modifications, and RlmB and RlmE on the 23S^70, 71, 72, 73^. *B. subtilis* contains homologs to RsmH, KsgA, and RlmB but does not contain a homolog for RlmE. Instead, *B. subtilis* contains the enzyme RlmP conferring a 2582-Gm modification at the *E. coli* equivalent position 2553 directly adjacent to the *E. coli* RlmE 2552-Um modification^26^. Further, ribosome assembly defects defined by sucrose gradient subunit quantification in *B. subtilis* show significant decreases in Δ*rlmB* 70S particles and a significant increase in 30S subunits. This differs from *E. coli* where Δ*rlmB* shows minimal distinct phenotype when grown under a variety of stressors^7, 74^. Therefore, our map of rRNA methylations, their methylating enzymes and functional relevance are more likely to be conserved among gram-positive bacteria than their *E. coli* counterparts.

This study provides a comprehensive framework for defining the enzymes, sites, and biological consequences of rRNA modifications. Our results demonstrate that no single analytical method is sufficient to define the full rRNA modification landscape, and an integrated strategy is required for confident modification mapping. Furthermore, we highlight the persistent challenges in *de novo* modification mapping, particularly for substoichiometric or poorly resolved modifications. Finally, we emphasize that mapping and cataloging should not be the only goal of modification studies as we demonstrate that loss of specific rRNA MTases have biological consequences. Most notably, deletion of MTases such as TlyA, KsgA, YacO, RsmG, and YbxB have defects in ribosome subunit accumulation or altered sensitivity to erythromycin and streptomycin, suggesting that methylations near key functional centers of the ribosome can influence both ribosome biogenesis and antibiotic interactions. Because many of the MTases characterized here are conserved across gram-positive pathogens **(Figure S25)**, these findings provide a foundation for understanding how rRNA modification shapes bacterial physiology and identify conserved modification enzymes as potential targets for antimicrobial development or determinants of clinically relevant drug resistance.

## Methods

### Strains and media

All strains were derived from *B. subtilis* strain PY79^48^. All strains, used in this work are listed in **Tables S2**. Strains were grown as described previously^24^ with detailed methods available in the Supplementary Materials. Spot plate assays were performed as described (^24^ and Supplementary Information).

### Total RNA isolation for LC-MS/MS and Nanopore DRS

Total RNA isolation for LCD-MS/MS and nanopore DRS was done as described previously^24, 40^. Briefly, 50 mL cultures were grown with appropriate antibiotics at 37°C until an OD600 0.6-0.8 was reached. For lysis, cell pellets were resuspended in 5.6 mLs NE (0.1 M NaCl, 0.05 M EDTA) and incubated at 37°C for 5 min followed by the addition of 300 μL fresh lysozyme (40 mg/mL) and incubation at 37°C for 15 min. 700 μL 10% N-lauroylsarcosine was added and the solution was incubated on ice for 5 min. Two extractions were done by adding 1 volume phenol/chloroform/iso-amyl alcohol (Millipore Sigma P19444), followed by centrifuging at max speed (3200 x g) for 30 min at 4°C. Nucleic acids were precipitated overnight at −20°C using 2.5 volumes of 100% ethanol and 1/10^th^ volume 3M NaOAc (pH 5.2). The nucleic acid pellet was washed twice with 70% ethanol and then dried for 15 min. The pellet was resuspended in 200 μL water, 20 μL DNase I incubation buffer, and 7 μL DNase I (Roche 04716728001), mixed by pipetting, and incubated for 30 min at 37°C. Following, 300 μL ddH2O was added to increase the reaction volume and the reaction was quenched with one phenol:chloroform:iso-amyl alcohol and one chloroform extraction. Total RNA was precipitated with 2.5 volumes of 100% ethanol and ½ volume 7.5M ammonium acetate and left overnight at −20°C or at −80°C for 2 hr. RNA was pelleted by centrifugation at 12,000 x g for 15 min at 4°C, washed twice with 70% ethanol, and left to dry for 10 min. After, RNA pellets were resuspended in 200 μL water and precipitated again using 2.5 volumes 100% ethanol and 1/10^th^ volume 3M NaOAC (pH 5.2), pelleted and washed as before, and dried for 15 min. Resulting pure total RNA was resuspended in 80 μL of water. A more detailed procedure is available in the Supplementary materials.

### Nanopore and MaP-seq

Direct RNA sequencing was performed according to Oxford Nanopore’s protocol (SQK-RNA004) using 200 ng total RNA isolated as described above and custom adapters designed for the 3′ end of *B. subtilis* 23S or 16S (**Table S2**). RNA was sequenced using an RNA flowcell (FLO-MIN004RA) and Oxford Nanopore’s MinION Mk1B technology. Two samples of each 23S and 16S were sequenced per strain. Reads were called using Doarado (1.0.2) with high accuracy base calling and modification location was identified by comparing WT and deletion strain reads using Nanocompore as previously described ^51, 53^. Mutational profiling and sequencing (MaP-seq) was done as described with the data analyzed using ShapeMapper^49, 52^ (see Supplementary Methods).

### Nucleoside Hydrolysis and LC-MS/MS analysis

Total nucleoside hydrolysis and LC-MS/MS analysis was done as described previously^24, 56^ with a detailed method available in the Supplementary Materials. We have expanded this method to include 58 nucleosides **(Table S3-14)**.

### Cloning

Cloning and plasmid construction is described in the supplemental materials. Oligonucleotides and plasmids are listed in **Tables S15-S16**, respectively.

## Author contributions

Conceptualization: JLH, KMS, LMM, LEB, KSK, LAS; Data curation: JLH, KMS, LMM, LEB, KSK, LAS; Formal analysis: JLH, KMS, LMM, LEB, KSK, LAS; Funding acquisition: JLH, KSK, LAS; Investigation: KSK, LAS; Methodology: JLH, KMS, LMM, LEB; Project administration: KSK, LAS; Resources: JLH, KMS, LMM, LEB; Supervision: KSK, LAS; Validation: JLH, KMS, LMM, LEB, KSK, LAS; Visualization: JLH, KMS, LMM, LEB, KSK, LAS; Writing-original draft: JLH, KMS, LMM, LEB, KSK, LAS; Writing-review &editing: JLH, KMS, LMM, LEB, KSK, LAS.

## Data availability

All raw nanopore and MaP-seq reads are deposited to SRA with the accession number SRA accession number PRJNA1450124. The LC-MS/MS raw data are available in Supplementary **Tables S3-S18**.

## Acknowledgements

We wish to thank Lainy Kulak for help with the initial spot plate growth assays, and Maddy Zamecnik and Miten Jain for help with Nanocompore analysis. This study was supported by funds from the National Institutes of Health grant R35GM131772 to LAS and R01HG013876 to KSK. In addition, JLH was supported in part by the Genetics Training Program Grant T32–GM007544 and the Edwards Award and the Cole summer Fellowships from MCDB.

