## Supplementary Tables S1-3, S15-S16; Figures S1-25. for "Mapping the rRNA methylome reveals contributions of methyltransferases to ribosome function and antibiotic sensitivity"

**Supporting Materials and Methods**

*Strains and media*

All strains were derived from *B. subtilis* strain PY79 [1]. All strains, primers, and plasmids used in this work are listed in Tables S1-S3. Typically, strains were grown in liquid lysogeny broth (LB) (10g/L NaCl, 10 g/L tryptone, and 5g/L yeast extract) or on LB agar plates using the same LB supplemented with agar (20 g/L). When necessary, spectinomycin or erythromycin at concentrations of 100 µg/mL or 0.5 µg/mL, respectively, were added to plates or cultures for selection. Strains were regularly grown at 37°C instead of the regular *B. subtilis* growth temperature of 30°C to account for cold sensitivity phenotype displayed in some strains [2]. Freezer stocks of each strain were made by mixing 10% dimethyl sulfoxide (DMSO) mixed with cultures at OD600 > 1.0 and stored at -80°C.

*Spot titer sensitivity assays*

Strains were grown overnight on LB agar plates at 37°C supplemented with selection antibiotics when appropriate. On the day of the experiment, LB or LB antibiotic plates were made with antibiotics and/or IPTG added at indicated concentrations in each figure. For cultures, one or two colonies was used to inoculate 2 mLs LB for each strain and grown at 37°C until OD600 was 0.5-0.8. Serial dilutions were done in 96 well plates using cultures normalized to an OD600 = 0.5 and diluted $10^{-5}$ in sterile saline. 5 µL were spotted onto appropriate plates and incubated for 10 hours at 37°C. All spot plates were done in biological triplicate with a technical replicate performed for each. Brightness and contrast of whole images was adjusted using ImageJ.

*Genomic DNA (gDNA) purification*

One colony of the desired strain was used to inoculate 5 mLs of LB and grown at 37°C to an OD600 = 2.0. The culture was centrifuged at 8,000 x g for 5 min at room temperature. Resulting supernatant was discarded, and the pellet was resuspended in 200 µL lysis buffer made with fresh lysozyme (50 mM Tris pH 8.0, 10 mM EDTA, 1% Triton X – 100, 0.5 mg/mL RNase A, 20 mg/mL lysozyme). The resuspended pellet was incubated at 37°C for 30 min before 20 µL of 10% SDS and 20 µL 10 mg/mL proteinase K were added and incubated at 55°C for 30 min. 500 µL of PB buffer (5 M GuHCl, 30% isopropanol) was added and the solution was mixed well via pipetting. The entire volume was loaded onto a spin column (Fisher NC0066803). The column was washed with 500 µL of PB buffer followed by washing with 750 µL PE buffer (10 mM Tris pH 7.5, 80% ethanol). All centrifugation steps were done at 10,000 x g for 1 min with an additional drying spin done after PE wash. DNA was eluted with 100 µL of ddH2O at 13,000 x g for 1 min. Concentration and quality was determined using NanoDrop, and gDNA was stored at -20°C.

*Determination of conservation of MTases across gram-positive species.*
Characterized methyltransferase conservation analyses were carried out through the NCBI Protein Basic Local Alignment Search Tool (BLAST) where the input query amino acid sequences were sourced from SubtiWiki (V5). The search set was against the non-redundant protein sequences (nr) database and specified to Staphylococcus aureus (taxid: 1280), Enterococcus faecalis (taxid: 1351), Streptococcus pyogenes (taxid: 1314), Mycobacterium tuberculosis (taxid: 1773), Streptococcus pneumoniae (taxid: 1313), Bacillus anthracis (taxid: 1392), and Clostridioides difficile (taxid: 1496). The seven species sampled are representative of common pathogenic gram-positive bacteria. Each unique pair of enzyme sequence and species ID were run through the blastp algorithm with the default parameters with the exception of max target sequences set to 20 and expect threshold set to 10. In each search, percent sequence identity, similarity, and query coverage were recorded. The data were assembled in R and depicted with heatmaps that report the *B. subtilis* enzyme name, its previous uncharacterized gene name, protein length, queued species, percent query coverage (qcov), and percent identity or similarity.

*Mutational profiling and sequencing (MaP-seq)*
23S and 16S rRNAs were isolated and extracted as mentioned above. Reverse transcription reactions were assembled by adding 9.3 µL of rRNA standardized to 200 ng and 1.7 µL of 60 µM NEB random primer mix and incubated at 65°C for 5 minutes and immediately cooled on ice. 8 µL of 2.5X MaP buffer (equal parts 30 mM $MnCl_2$ and 5X Pre-MaP buffer; 250 mM Tris (pH 8.0), 375 mM KCl, 50 mM DTT, and 2.5 mM each dNTP) was added and incubated at room temperature for 2 minutes, as described [3]. 1 µL of Invitrogen SuperScript II Reverse Transcriptase (RT) was added and left to incubate for 10 minutes at room temperature. Samples were immediately placed at 42°C and allowed the RT reaction to proceed for 3 hours. The reaction was heated for 15 minutes at 70°C to inactivate the RT and then cDNAs were purified using Cytiva MicroSpin G-25 columns according to the manufacturer's recommendations. Samples were measured with the Qubit ssDNA assay kit and stored at -20°C. Samples were sent to the Advanced Genomics Core at the University of Michigan in a skirted PCR plate for library prep and Illumina NovaSeq X 10B 300-cycle kit sequencing. Raw sequencing reads are reported in SRA accession number PRJNA1450124. ShapeMapper v2.2.0 [4]. https://github.com/Weeks-UNC/shapemapper2) was utilized with the following parameters to calculate mutation profiles --name [input file] --target mtase_data/[23S.fasta or 16S.fasta] --random-primer-len 6 --overwrite --min-depth 30000 --star-aligner --modified --folder [output folder]. The 23S.fasta and 16S.fasta reference sequences used for mapping were downloaded from the NIH GenBank and are from the *B. subtilis* *rrnC* and *rrnD* operons, respectively. The

ShapeMapper mutation profile output files were cumulated to 23S comparison and 16S comparison .xlsx files to be used for downstream analysis. Plotted mutation rates are the direct outputs from ShapeMapper mutational profiling converted to percentages. Mutation rate fold changes and differences were calculated using the mutation rate of the sample of interest compared to the median mutation rate of all samples sequenced at each nucleotide on the 23S and 16S.

*Deletion and complementation strain construction and transformation*
Gene substitution strains were obtained from the Bacillus Genetic Stock Center (BGSC). Catalog numbers are listed in Table S1. Genomic DNA was isolated as described above from these strains and used to transform PY79. For all transformations, strains were made competent by first growing in 2 mL LM media (LB supplemented with 3mM $MgSO_4$) at 37°C on a rolling rack for approximately 3 hours or until OD600 = ~1. Following, 500 µL pre-warmed MD minimal media (1x PC buffer [10X PC buffer: 107 g/L $K_2HPO_4$, 60 g/L $KH_2PO_4$, 10 g/L trisodium citrate (pentahydrate)], 2% glucose, 50 µg/mL tryptophan, 50 µg/mL phenylalanine, 11 µg/mL ferric ammonium citrate, 2.5 µg/mL potassium aspartate, 3 mM $MgSO_4$) was inoculated with 20 µL culture and grown in a rolling rack at 37°C for 4 hours. For transformation, up to 100 ng gDNA or plasmid was added to competent strains and grown for 1 hour at 37°C in a rolling rack. Cultures were plated on appropriate antibiotic containing plates for selection. Clean deletion strains with only a *loxP* scar were generated using competent gene substitution strains transformed with Cre recombinase driven from plasmid pDR244 as previously described [5]. All strains were made into clean deletions as described except for *yacO.* This strain exhibited genomic instability when transformed with pDR244 despite multiple trials. Strain Δ*yydA* was cleanly deleted using pMiniMAD by placing ~1000 bases upstream and downstream of *yydA* into the pMiniMAD backbone using Gibson assembly as described above and recovery and screening was done performed previously described [5]. Resulting strains were confirmed via PCR amplification of the *yydA* locus.

*Total RNA isolation for LC-MS/MS and Nanopore*
50 mL cultures were grown with appropriate antibiotics at 37°C at 200 rpm until an OD600 0.6-0.8 was reached. Forty mLs of each culture was centrifuged at 7,500 rpm for 15 min at room temperature. For lysis, cell pellets were resuspended in 5.6 mLs NE (0.1 M NaCl, 0.05 M EDTA) and incubated at 37°C for 5 min followed by the addition of 300 µL fresh lysozyme (40 mg/mL) and incubation at 37°C for 15 min. 700 µL 10% N-lauroylsarcosine was added and the solution was incubated on ice for 5 min. Following lysis, two extractions were done by adding 1 volume phenol/chloroform/iso-amyl alcohol (Millipore Sigma P19444), shaking and vortexing vigorously, centrifuging at max speed (3200 x g) for 30 min at 4°C, and transferring the aqueous phase into a new tube. One

chloroform extraction was then done to ensure removal of all traces of phenol. Nucleic acids were precipitated overnight at -20°C using 2.5 volumes of 100% ethanol and 1/10$^{th}$ volume 3M NaOAc (pH 5.2). Precipitated nucleic acids were pelleted by centrifugation at max speed (3200 x g) for 15 min at 4°C. The pellet was washed twice with 70% ethanol and then dried for 15 min. To digest the DNA present, the pellet was resuspended in 200 µL water, 20 µL DNase I incubation buffer, and 7 µL DNase I (Roche 04716728001), mixed by pipetting, and incubated for 30 min at 37°C. Following, 300 µL ddH2O was added to increase the reaction volume and the reaction was quenched with one phenol:chloroform:iso-amyl alcohol and one chloroform extraction. Total RNA was precipitated with 2.5 volumes of 100% ethanol and ½ volume 7.5M ammonium acetate and left overnight at -20°C or at -80°C for 2 hrs. RNA was pelleted by centrifugation at 12,000 x g for 15 min at 4°C, washed twice with 70% ethanol, and left to dry for 10 min. After, RNA pellets were resuspended in 200 µL water and precipitated again using 2.5 volumes 100% ethanol and 1/10$^{th}$ volume 3M NaOAC (pH 5.2), pelleted and washed as before, and dried for 15 minutes. Resulting pure total RNA was resuspended in 80 µL of water. Concentration was determined using qubit and Nanodrop and purity was determined using Nanodrop.

*Nucleoside Hydrolysis and LC-MS/MS analysis*
16S and 23S rRNA purified from WT *B. subtilis* and 26 deletion mutants was subject to a two-step enzymatic digestion resulting in nucleosides, as previously described [6]. Briefly, RNA (200 ng) was digested overnight to nucleotides using 300 U/µg nuclease P1 (100,000 U/mL, NEB) at 37°C in 100 mM ammonium acetate pH 5.3 and 100 µm zinc sulfate. The nucleotide mixture was then dephosphorylated using 50 U/µg bacterial alkaline phosphatase (150 U/µL, Invitrogen) at 37°C for 4 hours in 100 mM ammonium bicarbonate pH 8.1 and 100 µM zinc sulfate. Prior to digestion, nuclease P1 and bacterial alkaline phosphatase were buffer exchanged into their respective buffers using Micro Bio-Spin 6 desalting columns (BioRad). The resulting nucleoside mixture was lyophilized and reconstituted in 40 µM $^{15}N_4$-Inosine in water.

Nucleosides were analyzed using an Agilent 1290 II LC interfaced to either an Agilent 6410 triple quadrupole mass spectrometer with an electrospray ionization (ESI) source or an Agilent 6460 triple quadrupole mass spectrometer with an Agilent Jet Stream source. Nucleosides were separated using reversed-phase chromatography with a Water’s Acquity HSS T3 column (1.0 x 100 mm, 1.8 µm, 100 Å) with an attached guard column (2.1 x 5 mm, 1.8 µm, 100 Å). LC conditions used were as previously described (Table S3-18) [6]. After separation, nucleosides were detected in positive mode using multiple reaction monitoring mode (Table S3-18). The ESI source on the 6410 QQQ had a gas temperature of 350°C, a gas flow rate of 10 L/min, and the nebulizer gas pressure was 25 psi. The AJS source on the 6460 QQQ had a gas temperature of 350°C, a gas flow of 5 L/min, a nebulizer pressure of 45 psi, and a sheath gas flow rate of 11 L/min.

To quantify the abundance of modified nucleosides in rRNA samples, calibration curves with 58 nucleosides in 40 μM $^{15}N_4$-Inosine internal standard were used. Previously, this method included 50 nucleosides [6]. We have expanded this method to include 58 nucleosides (Table S3-14). Samples were analyzed using three biological replicates times, thus resulting in three calibration curves used to quantify the abundances of modified nucleosides **(SI Figure S4-S15 and SI Tables S3-5)**. For samples tested on various days, the signals were normalized using a sample that was analyzed during all three experiments. For the 16S and 23S rRNA, the signals were normalized using the WT RNA and Δ*ylbH*, respectively. RNA from Δ*ylbH* was used because it was included along with each 23S calibration curve and YlbH does not modify 23S rRNA.

*Sucrose gradient ultracentrifugation*

Ten mL 20-40% sucrose gradients were made as described [2] using buffer G (20 mM Tris-HCl pH 7.5, 200 mM $NH_4Cl$, 6 mM ß-mercaptoethanol), 1 mM Mg(OAc)2 for dissociating conditions or 10 mM Mg(OAc)2 for associating conditions, and sucrose to desired concentration [w/v] added to 13.2 mL tubes (Beckman Coulter 344059). Cultures were inoculated by plate washing and inoculating 160 mLs LB at a starting OD600 = 0.01. Cultures were grown at 25°C or 37°C until OD600 0.5-0.6. The culture was split and 75 mLs was pelleted by spinning at max speed for 10 min at room temperature. Pellets were stored at -80°C until used. Pellets were resuspended in either associating or dissociating buffer G, 1 μL 50 mg/mL lysozyme, and ¼ pellet protease inhibitors (Pierce, EDTA-free). Lysis was done by freezing at -80°C for 2 min and then thawing to room temperature followed by sonication on ice (1 min–10 sec on/10 sec off, 25% amplitude). DNA was digested using DNase I (Roche 04716728001) according to manufacturer's instructions. Lysate was cleared by centrifugation at 16,000 x g for 30 min at 4°C. Approximately 15 A260 units (1 mL lysate) were added to the top of the gradient. Gradients were separated by ultracentrifugation in a Thermo TH-641 swinging bucket rotor at 25,000 rpm for 15 hrs at 4°C. Gradients were fractionated by hand into 96 well plates. Fractions were diluted 1:10 into UV-transparent plates (Grenier One) and A260 was determined with a Tecan Spark Spectrophotometer. Absorbances were plotted as curves in R by fitting with a smoothing spline. Area under the curve was quantified for each peak using the Simpson's Rule method and normalized by the area of all three peaks. Significance was determined using an ANOVA with Bonferroni post hoc analysis using four replicates of each strain.

**Table S1. Sequence identity of known or putative *B. subtilis* rRNA MTases with known bacterial enzymes**

| *E. coli* or (*M. tuberculosis*) MTase | Modification site | Putative *B. subtilis* MTase | Percent identity of *B. subtilis* gene to known protein |
|---|---|---|---|
| 23S rRNA | | | |
| TlyA (*M. tuberculosis*) | Cm1920 | YqxC[2, 7] | 41.4% (TlyA) |
| RlmB | Gm2251 | YacO/RlmB | 75.5% (RlmB) |
| RlmB | Gm2253 | YsgA [8] | 41.2% (23S MT) 35% (RlmB) |
| RlmA | $m^1$G745 | YxjB | 33.1% (RlmA) |
| RlmA | $m^1$G745 | YcgJ | 35.7% (SAM) |
| RlmG | $m^2$G1835 | YqeM | 38.0% (SAM) |
| RlmKL | $m^7$G2069, | YpsC | 32.7% (RlmKL) 57% (SAM) |
| RlmKL | $m^7$G2069, | YxbB | 36.5% (SAM) |
| RlmN | $m^2$A2503 | YloN | 59.3% (RlmN) |
| RlmJ | $m^6$A2030 | YbdB | 50% (Cupid domain-containing) |
| RlmI | $m^5$C1962 | YwbD[9] | 33.6% (RlmI) 57.8% (I/K Family) |
| RlmC | $m^5$U747 | YefA [10] | 73.5% (RlmD) |
| RlmD | $m^5$U1939 | YefA [10] | 73.5% (RlmD) |
| RlmD | $m^5$U1939 | YfjO (RlmS)[11] | 73.5% (RlmD) |
| 16S rRNA | | | |
| TlyA (*M. tuberculosis*) | Cm1409 | YqxC [2, 7] | 41.4% (TlyA) |
| RsmH | $m^4$Cm1402 | MraW/YlxA | 66.0%(RsmH) |
| RsmI | $m^4$Cm1402 | YabC | 47.5% (RsmI) |
| RsmH | $m^4$Cm1402 | YtqB | 51.0% (RsmH) |
| RsmB | $m^5$C967 | YloM | 45.6% (16S rRNA MT) |
| RsmD | $m^2$G966 | YbH | 48.9% (RsmD) |
| RsmC | $m^2$G1207 | YbxB | 32.3% (RsmC) |
| RsmJ | $m^2$G1516 | YodH | 44.6% (MT domain-containing) |
| RsmJ | $m^2$G1516 | YisL | 43.3% (DUF1516) |
| RsmG | $m^7$G527 | RsmG [12] | 70.2% (RsmG) |
| RsmA | $m^6$A1518 | KsgA | 59.2% (RsmA) |
| RsmA | $m^6$A1519 | YrrT | 57.1% (SAM) |
| RsmE | $m^3$U1498 | YqeU/RsmE | 50.0% (16S rRNA MT) |

BLAST was used to identify conserved amino acid sequence between methyltransferases in *E. coli Mtb* and *B. subtilis*. The columns on the left list known methyltransferases and their respective methylation site(s) [13, 14]. The two columns on the right half identify the *B. subtilis* gene name with the most sequence similarity to characterized MTases and their percent amino acid sequence identity.

**Supplementary Table S2: Strains used in this work**

| **Strains** | **Genotype** | **Citation** |
|---|---|---|
| JLH005 | WT PY79 | [15] |
| JLH010 | Δ*tlyA* | [2] |
| JLH018 | Δ*tlyA*, *amyE*::$P_{hyperspank}$-*tlyA* | [2] |
| JLH020 | *E. coli* MC1061 | Lab Stock |
| JLH078 | Δ*yefA* | This study |
| JLH079 | Δ*yodH* | This study |
| JLH080 | Δ*rsmE* | This study |
| JLH081 | Δ*ypsC* | This study |
| JLH082 | Δ*yfjO* | This study |
| JLH083 | Δ*ywbD* | This study |
| JLH084 | Δ*rsmG* | This study |
| JLH086 | Δ*yxjB* | This study |
| JLH087 | Δ*ksgA* | This study |
| JLH088 | Δ*mraW* | This study |
| JLH123 | Δ*ycgJ* | This study |
| JLH124 | Δ*ysgA* | This study |
| JLH125 | Δ*yloN* | This study |
| JLH126 | Δ*yabC* | This study |
| JLH127 | Δ*ylbH* | This study |
| JLH128 | Δ*rsmB* (aka Δ*yloM*) | This study |
| JLH129 | Δ*yrrT* | This study |
| JLH130 | Δ*yisL* | This study |
| JLH131 | Δ*ydbB* | This study |
| JLH132 | Δ*yxbB* | This study |
| JLH133 | Δ*ybxB* | This study |
| JLH134 | Δ*yqeM* | This study |
| JLH135 | Δ*ytqB* | This study |
| JLH233 | *rlmB*::*kan* (*yacO*::kan) | This study |
| JLH269 | Δ*yydA* | This study |

**Supplemental Tables S3-14 are in a separate excel file. Tables S3-14 contain the LC MS/MS data for all replicates.**

**Supplementary Table S15: Primers used in this work**

| Primer | Sequence | Purpose |
|---|---|---|
| PEB3F | GCTAGCCGCATGCAAGCTAATTCG | for amplifying pPB194 or pDR110 |
| PEB259 | ATGTATACCTCCTTAGTCGACTAAGCTTAATTGTTATCCGCT CACAATTACACACATTATGCCACACCTTGTAGATA | for amplifying pPB194 or pDR110 |
| oPEB866 | TTTATGCAGCAATGGCAAGAAC | outside *amyE* locus to check for GOI insertion |
| oPEB867 | GCCGACTCAAACATCAAATCTTAC | outside *amyE* locus to check for GOI insertion |
| JLH015 | GCTGATATGATGTTGGCGGG | upstream 271 bp of *yxjB* |
| JLH015a | CGTTTCCGCGAACTAGAAAG | downstream 147 bp of *yxjB* |
| JLH016 | TGAATGACGGCATGTTTGAT | upstream 527 bp of *yefA* |
| JLH016a | AGATTGCAGGAGGTCAATGC | downstream 602 bp of *yefA* |
| JLH017 | CAAAGCCAAGAAAAGCAAGC | upstream 265 bp of *ywbD* |
| JLH017a | TGGCATTCTGAGCATAGCAG | downstream 370 bp of *ywbD* |
| JLH018 | GAAAGTGCCACAGTGACGAA | upstream 313 bp of *ypsC* |
| JLH018a | ATCACCGAAACATGCACAAA | downstream 295 bp of *ypsC* |
| JLH019 | AATGCCAAGTCAGGGACAAC | upstream 166 bp of *yacO/rlmB* |
| JLH019a | TAGTCAGAGGTCGCAACGTG | downstream 321 bp of *yacO/rlmB* |
| JLH020 | TTTGCGTCTGACAGAAATCG | upstream 566 bp of *rsmG* |
| JLH020a | CGCTCGGTGATTTTCTTCTC | downstream 658 bp of *rsmG* |
| JLH021 | TGAAGTTGTGGAGCGAAGTG | upstream 564 bp of *ylxA/rsmH** |
| JLH021a | TTCTCCCCAGGATGACAAAG | downstream 487 bp of *ylxA/rsmH** |
| JLH022 | GCACTGGTTCGGGAATTTTA | upstream 411 bp of *yqeU/rsmE* |
| JLH022a | TTTCAATGACGCTCTTGTGC | downstream 444 bp of *yqeU/rsmE* |

| | | |
|---|---|---|
| JLH023 | CTGATTTTTGGCCCAGGATA | upstream 261 bp of *yodH* |
| JLH023a | TCTCCCGCCTCTCTCTGTAA | downstream 160 bp of *yodH* |
| JLH024 | TAAAAACAAGCGGGGAATTG | upstream 293 bp of *ksgA* |
| JLH024a | AGCAATTCGAGCGACTCATT | downstream 398 bp of *ksgA* |
| JLH025 | GGCACACTCAGCTTGTCAGA | upstream 398 bp of *yfjO* |
| JLH025a | TTCACCTGCTTTTCAACACG | downstream 202 of *yfjO* |
| JLH041 | GATTTGCTGGCTGAGAGACG | upstream 225 bp of *ycgJ* |
| JLH041a | CGCCGAAGATGAACGCTAAA | downstream 597 bp of *ycgJ* |
| JLH042 | TTGCGATGATAAACGGGCAG | upstream 315 bp of *yqeM* |
| JLH042a | GACATCACGAGGGTTGCATC | downstream 411 bp of *yqeM* |
| JLH043 | GAAGCAGGTAGAAAAGGCGG | upstream 473 bp of *ydbB* |
| JLH043a | CGCGAGTAAAGAGACAAGCC | downstream 399 bp of *ydbB* |
| JLH044 | CGATTCGGTTACAGCACTCA | upstream 529 bp of *yxbB* |
| JLH044a | TCGTCTGGTAAAGCAAGCAC | downstream 308 bp of *yxbB* |
| JLH045 | CAGCTGTCGATCTTGAGGGA | upstream 270 bp of *yloN* |
| JLH045a | GCACATACAATCGTCGTCCC | downstream 327 bp of *yloN* |
| JLH046 | GTGCTGCCAAATGACTTCGA | upstream 268 bp of *ysgA* |
| JLH046a | TTCCGAGATATTGCACCCGA | downstream 458 bp of *ysgA* |
| JLH047 | CATCTGCGGGGTCCTTCTAT | upstream 301 bp of *ylbH* |
| JLH047a | TACTTCCGGCGGTACAAACT | downstream 455 bp of *ylbH* |
| JLH048 | ATGATGGAAAGGGCGCAAAG | upstream 174 bp of *yabC* |
| JLH048a | ACTGCGTCGTACTCTTGGAA | downstream 272 bp of *yabC* |
| JLH049 | ATGATGTGGTCGGCATTTCG | upstream 604 bp of *ytqB* |

| | | |
|---|---|---|
| JLH049a | GATGATGGACCGGCTTGAAC | downstream 273 bp of *ytqB* |
| JLH050 | TGAAGCGGGCGACATTATTG | upstream 255 bp of *yisL* |
| JLH050a | ACTCTTTACGGTCAGTGCCA | downstream 411 bp of yisL |
| JLH051 | GAAATAACGGCAGCCCTCAG | upstream 536 bp of *yrrT* |
| JLH051a | CTACATCATGGTGGCGAACG | downstream 322 bp of *yrrT* |
| JLH052 | ATGCGGGCGATATGATCTCA | upstream 510 bp of *yloM* |
| JLH052a | AGCCGTTTTCAGCGTTGTTA | downstream 253 bp of *yloM* |
| JLH057 | TTCAAGCTCCAGTTCGCAAC | upstream 580 of *ybxB* |
| JLH057a | TTCACCCCTCAAATCATGCG | downstream 241 of *ybxB* |
| oJLH058 | /5PHOS/GGCTTCTTCTTGCTCTTAGGTAGTAGGTTC | nanopore direct RNA sequencing oligo A |
| oJLH058a | GAGGCGAGCGGTCAATTTTCCTAAGAGCAAGAAGAAGCCAGAAAGGAGG | nanopore direct RNA sequencing oligo B (16S specific) |
| oJLH058b | GAGGCGAGCGGTCAATTTTCCTAAGAGCAAGAAGAAGCCATGGTTAAGT | nanopore direct RNA sequencing oligo B (23S specific) |
| JLH157 | CCCAGGCTTTACACTTTATGCTTCCCGCAAAAGCCGTCAATGCCAC | *yydA*-up-for for pMinimad Gibson |
| JLH157a | GTCTTTCTGTAATGGTGGAGATGCTCAGGTCATCCCCACTTTAAAAACAAGTTATTC | yydA-up-rev for pMinimad Gibson |
| JLH160 | GAATAACTTGTTTTTAAAGTGGGGATGACCTGAGCATCTCCACCATTACAGAAAGAC | *yydA*-down-for pMinimad Gibson |
| JLH160a | CCGCACAGATGCGTAAGGAGAAAATACCCGGAAGATTCAATACAACCTAACGTATGG | *yydA*-down-rev pMinimad Gibson |
| JLH161 | CGAAACGGTGCTGGATATGT | 216 bp upstream of the *yydA* |
| JLH161a | GACTCTTTCCACACCATCAA | 172 bp downstream of the *yydA* |

### Supplementary Table S16: Plasmids used in this work

| | | | |
|---|---|---|---|
| pPB194 | pDR110 | spc$^R$ | [16] |
| pDR244 | pDR243 | Cre-recombinase | Bacillus Genetic Stock Center (ECE274) |
| pJH029 | pPB194 | *yefA*, spc$^R$ | This study |
| pJH030 | pPB194 | *rsmE,* spc$^R$ | This study |
| pJH031 | pPB194 | *ypsC,* spc$^R$ | This study |
| pJH032 | pPB194 | *rsmG,* spc$^R$ | This study |
| pJH033 | pPB194 | *yxjB,* spc$^R$ | This study |
| pJH034 | pPB194 | *mraW,* spc$^R$ | This study |
| pJH035 | pPB194 | *ycgJ,* spc$^R$ | This study |
| pJH036 | pPB194 | *ysgA,* spc$^R$ | This study |
| pJH037 | pPB194 | *ybxB,* spc$^R$ | This study |
| pJH038 | pPB194 | *yisL,* spc$^R$ | This study |
| pJH039 | pPB194 | *rlmB,* spc$^R$ | This study |
| pJH040 | pPB194 | *yloN,* spc$^R$ | This study |
| pJH041 | pPB194 | *yloM,* spc$^R$ | This study |
| pJH042 | pPB194 | *yabC,* spc$^R$ | This study |
| pJH043 | pPB194 | *yqeM,* spc$^R$ | This study |
| pJH044 | pPB194 | *ylbH,* spc$^R$ | This study |
| pJH045 | pPB194 | *ydbB,* spc$^R$ | This study |
| pJH046 | pPB194 | *yrrT,* spc$^R$ | This study |
| pJLH054 | pMiniMad2 | amp$^R$,erm$^R$ | [17] |
| pJLH055 | pMiniMad2 | *yydA,* amp$^R$,erm$^R$ | This study |

Plasmids pJH029-046 are complementing plasmids that allow for expression of each allele from the chromosomal ectopic locus amyE using an $P_{hyperspank}$ IPTG inducible promoter

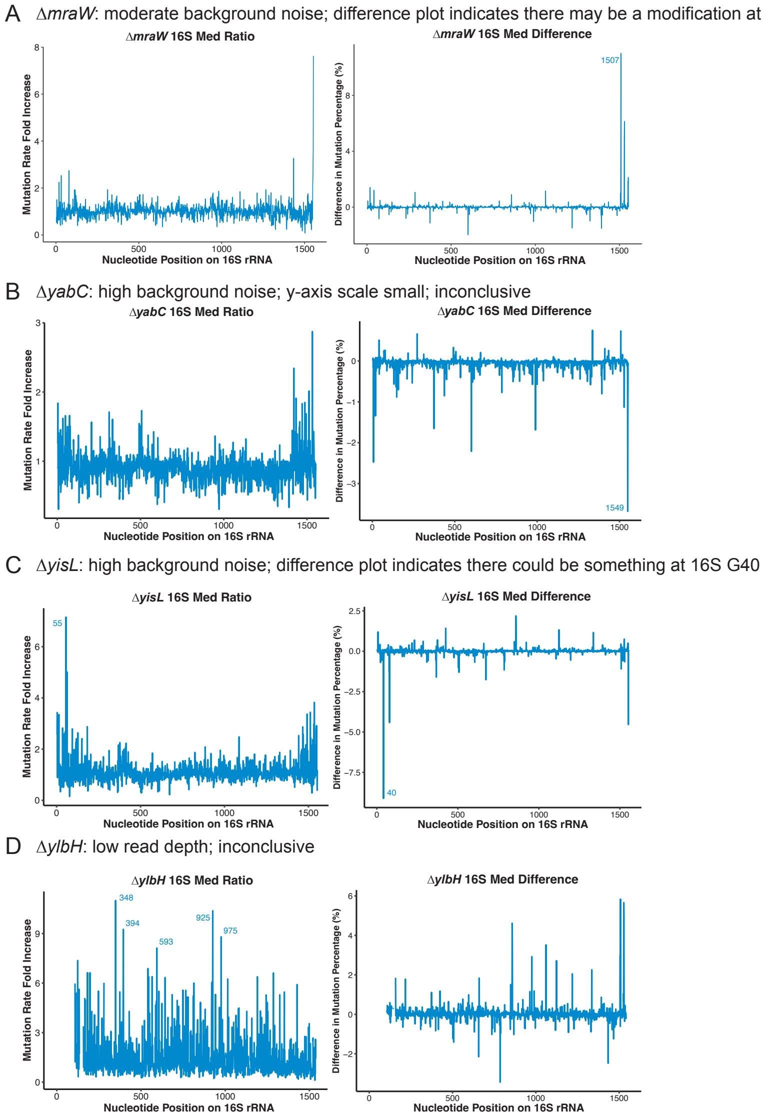


**Supplemental Figure S1. MaP-seq results that lead to positive identification of nucleotide position modified or inconclusive results for 16S rRNA.** MaP-seq plots displaying median mutation rate fold increase and difference in mutation percentage (%) for 16S putative rRNA methyltransferases (A) Δ*mraW*, (B) Δ*yabC*, (C) Δ*yisL,* (D) Δ*ylbH*, predicted to methylate within the Watson-Crick face. All contained high background noise or low read depth making results inconclusive. Possible modifications indicated in difference plots did not match existing literature[7] regarding present modifications.

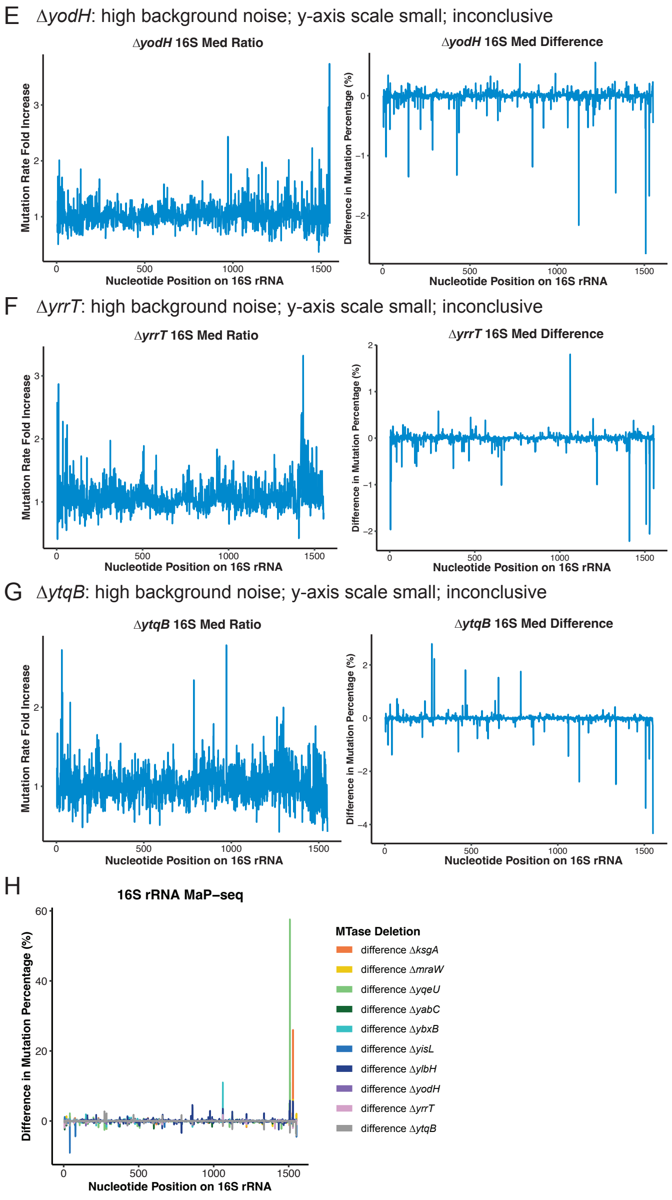


**Supplemental Figure S1 continued. MaP-seq results that lead to positive identification of nucleotide position modified or inconclusive results for 16S rRNA.** MaP-seq plots displaying median mutation rate fold increase and difference in mutation percentage (%) for 16S putative rRNA methyltransferases (E) Δ*yodH*, (F) Δ*yrrT*, and (G) Δ*ytqB* predicted to methylate within the Watson-Crick face. All contained high background noise or low read depth making results inconclusive. Possible modifications indicated in difference plots did not match existing literature[7] regarding present modifications. (H) Summary plot displays difference in mutation percentage of all candidates displaying three methyltransferases Δ*ksgA*, Δ*yqeU,* and Δ*ybxB* with clear differences allowing location determination.

A ∆*ycgJ*: high background noise; y-axis scale small; inconclusive

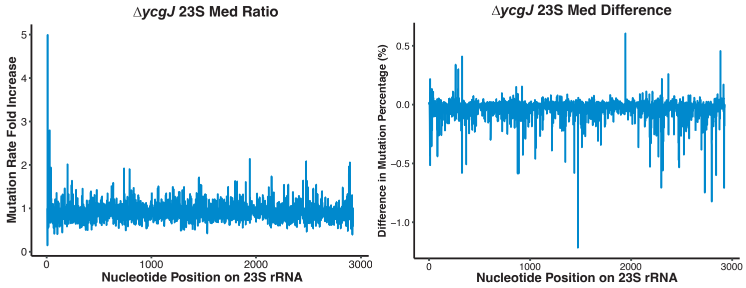


B ∆*ydbB*: high background noise; y-axis scale small; inconclusive

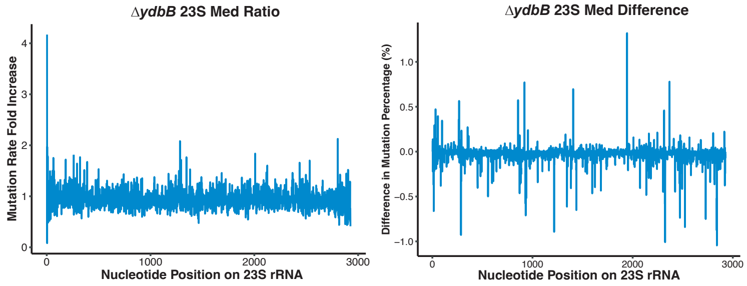


C ∆*yqeM*: moderate background noise; difference plot indicates there may be a modification at 23S G11

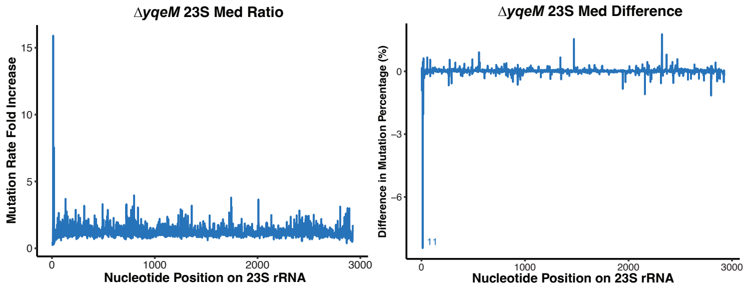


D ∆*yxbB:* high background noise; difference plot indicates there may be modifications at 23S C387, G1406

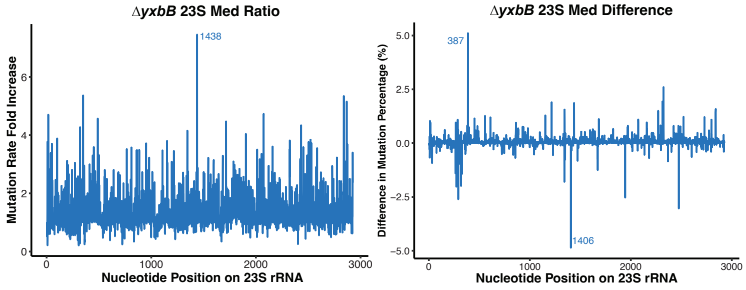


**Supplemental Figure S1 continued. MaP-seq results that lead to positive identification of nucleotide position modified or inconclusive results for 23S rRNA.** MaP-seq plots displaying median mutation rate fold increase and difference in mutation percentage (%) for 23S putative rRNA methyltransferases (A) ∆*ycgJ*, (B) ∆*ydbB*, (C) ∆*yqeM,* (D) ∆*yxbB*, predicted to methylate within the Watson-Crick face. All contained high background noise making results inconclusive. Possible modifications indicated in difference plots did not match existing literature[7] regarding present modifications.

E ΔyxjB: high background noise; y-axis scale small; inconclusive

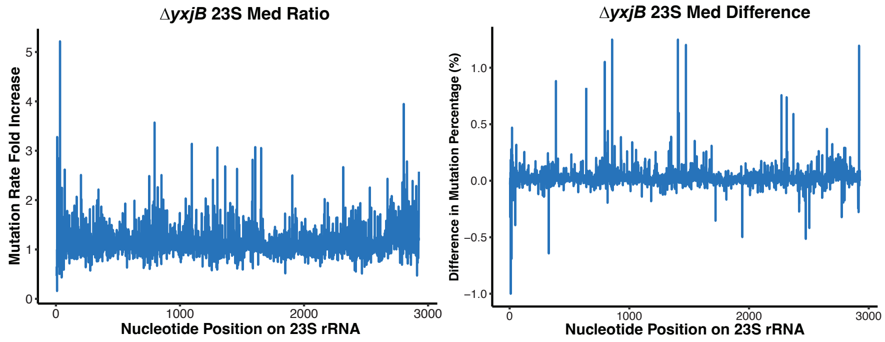


F

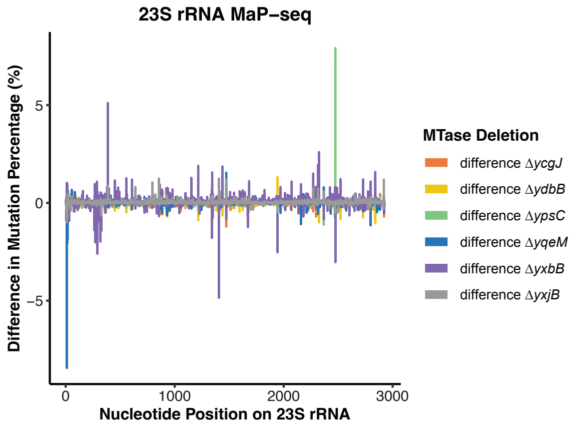


**Supplemental Figure S1 continued. MaP-seq results that lead to positive identification of nucleotide position modified or inconclusive results for 23S rRNA.** MaP-seq plots displaying median mutation rate fold increase and difference in mutation percentage (%) for 23S putative rRNA methyltransferases (E) Δ*yxjB* predicted to methylate within the Watson-Crick face. All contained high background noise making results inconclusive. Possible modifications indicated in difference plots did not match existing literature[7] regarding present modifications. (F) Summary plot displays difference in mutation percentage of all candidates displaying one methyltransferase Δ*ypsC* with a clear difference allowing location determination.

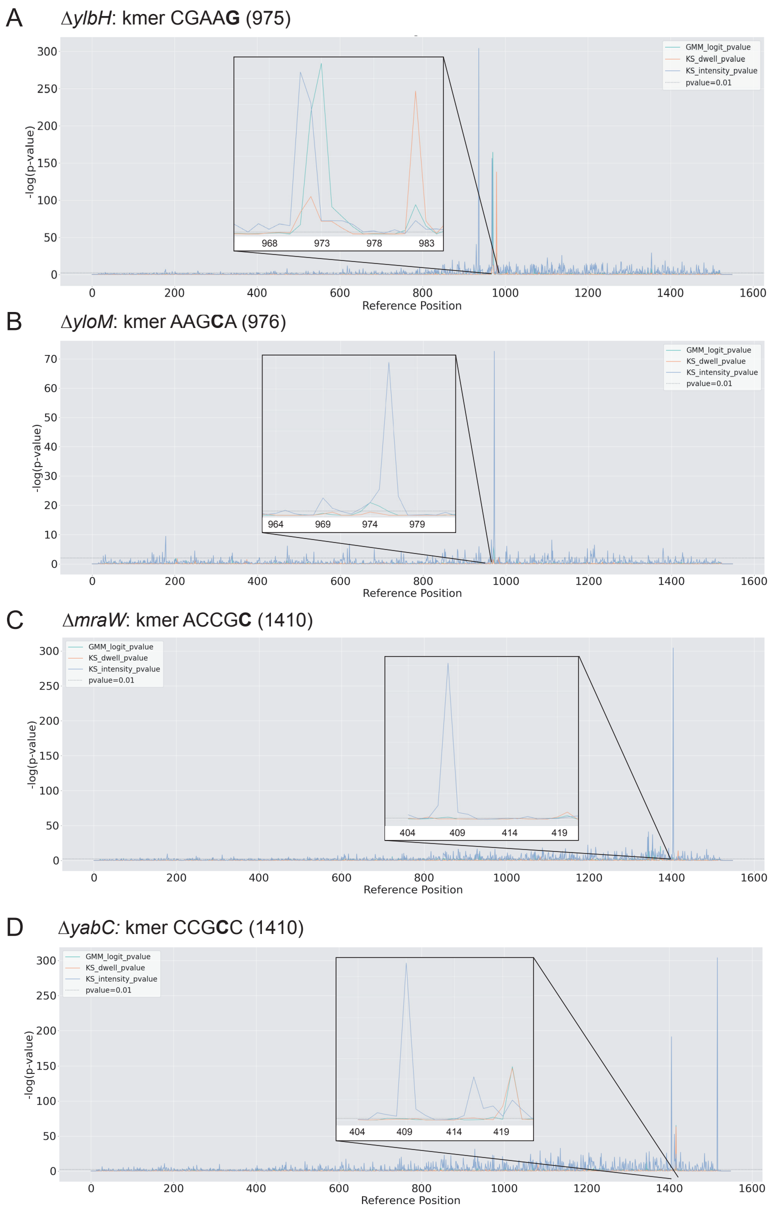


**Figure S2. 16S Nanopore sequencing results.** Shown are the Nanocompore analysis for modifications on the 16S rRNA in the indicated background. Shown are the p-values from logistic regression log odds ratio (GMM) and Kolmolgorov-Smirnov test for peak intensity value and dwell time. The strains shown are: **A)** Δ*ylbH*, **B)** Δ*yloM*, **C)** Δ*mraW*, **D)** Δ*yabC.*

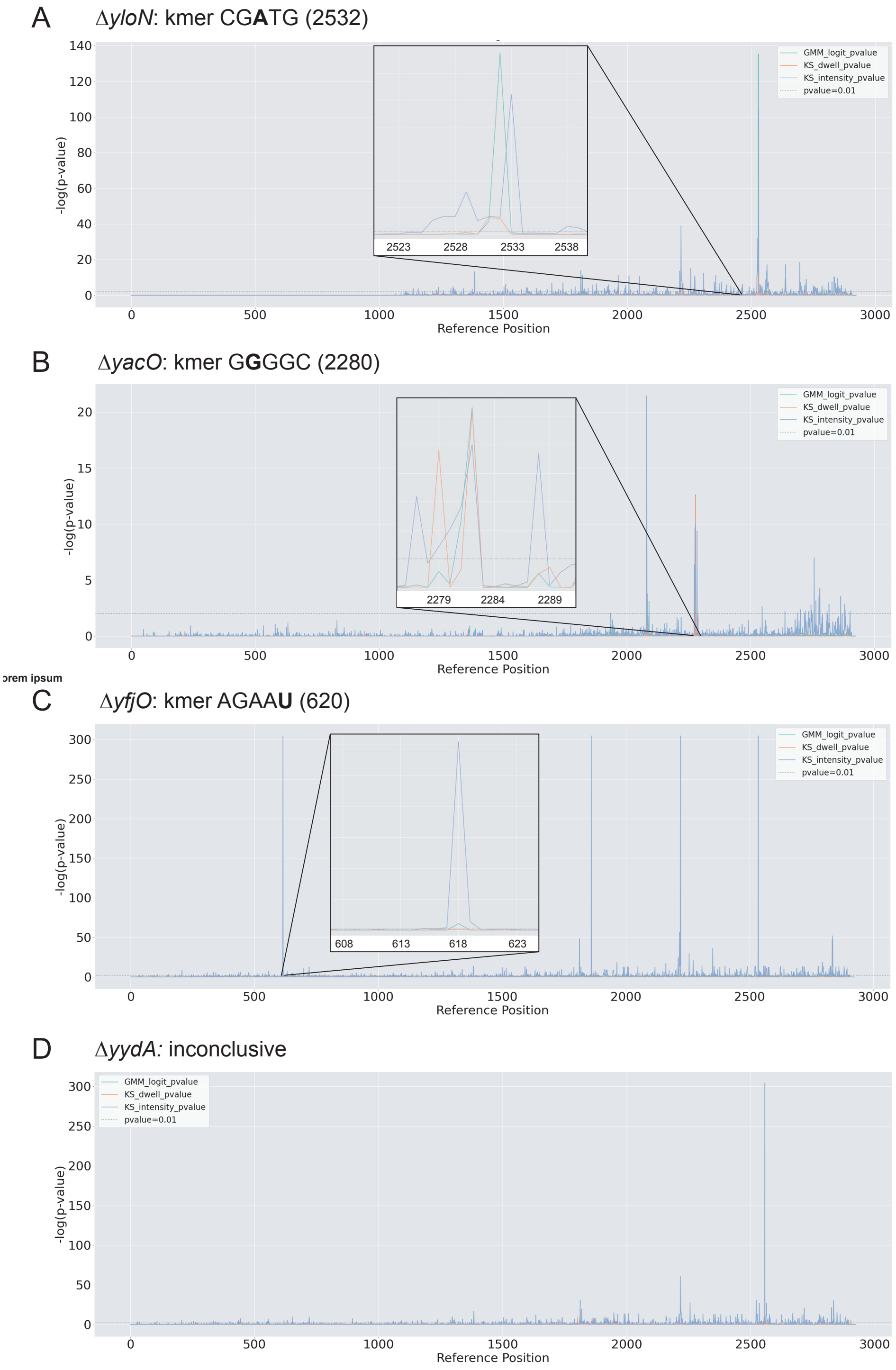


**Figure S3. 23S Nanopore sequencing results.** Shown are the Nanocompore analysis for modifications on the 23S rRNA in the indicated background. Shown are the p-values from logistic regression log odds ratio (GMM) and Kolmolgorov-Smirnov test for peak intensity value and dwell time. The strains shown are: **A)** Δ*yloN*, **B)** Δ*yacO*, **C)** Δ*yfijO*, **D)** Δ*yydA.*

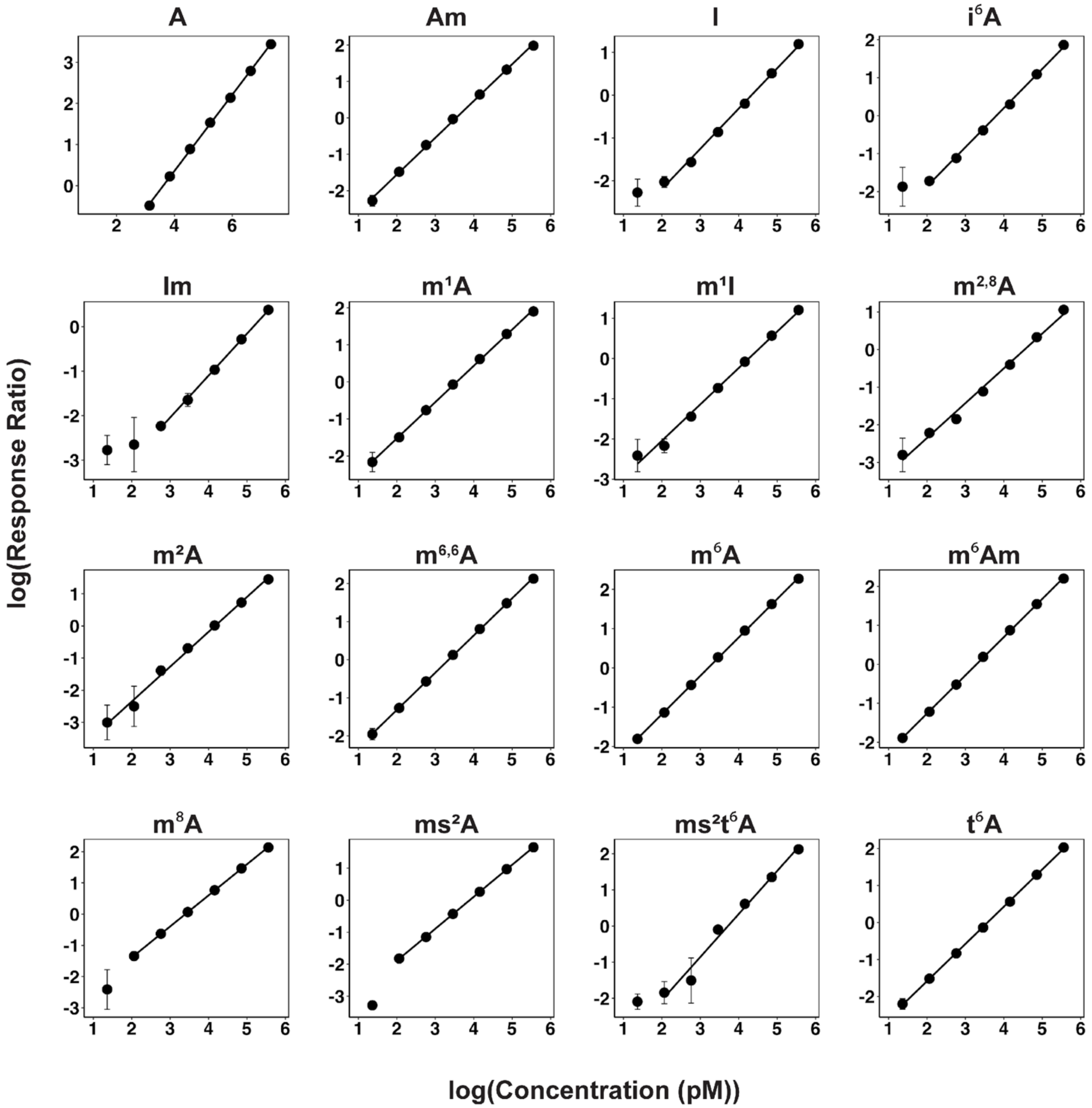


**Supplemental Figure S4.** Calibration curves used to quantify the concentration of adenosine modifications from set A. These data were run on the Agilent 6410 QQQ. Calibration curves are plotted with log(response ratio) vs log(concentration (pM)). Linear regression, limit of detection, and $R^2$ are reported in Table S3.

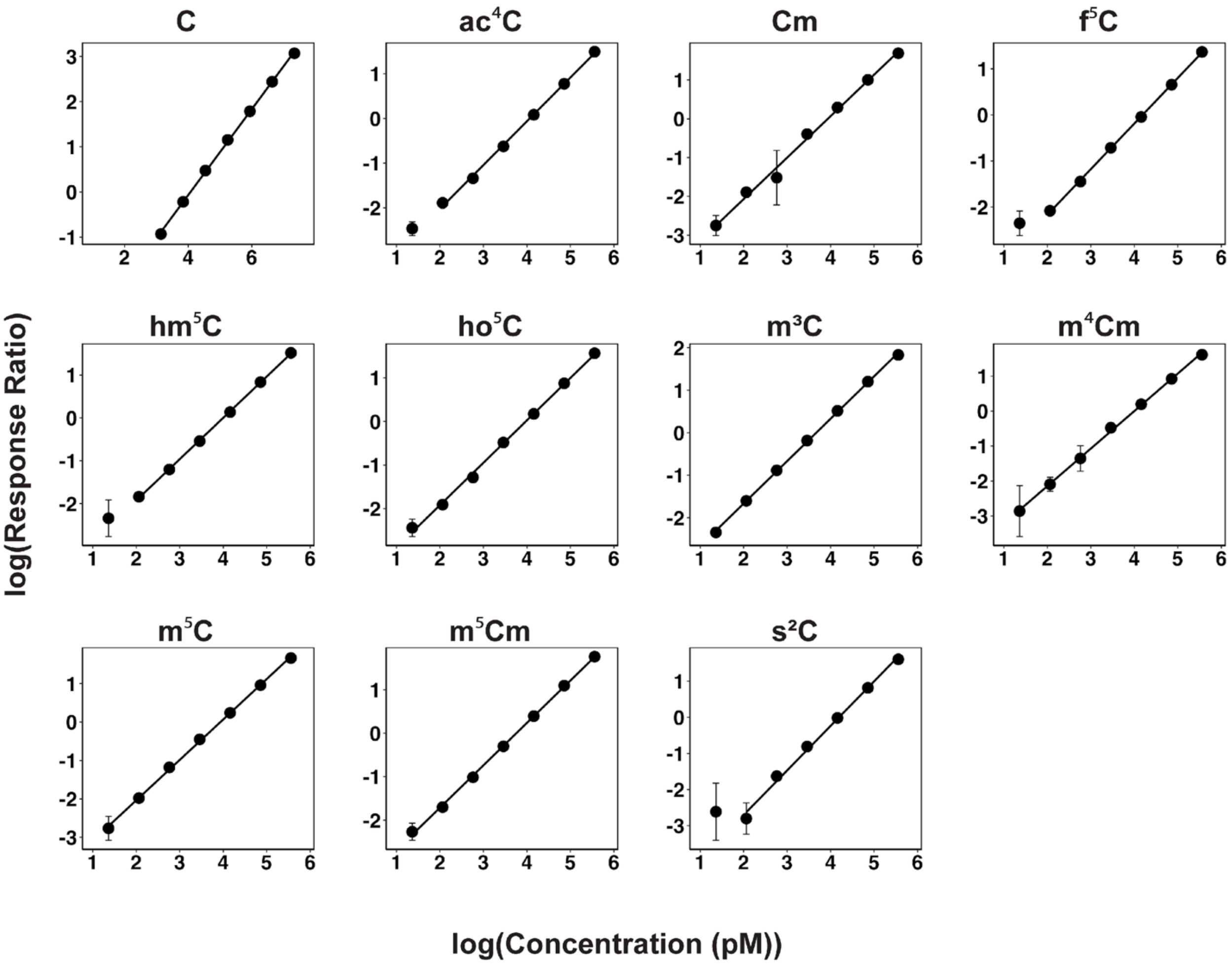


**Supplemental Figure S5.** Calibration curves used to quantify the concentration of cytidine modifications from set A. These data were run on the Agilent 6410 QQQ. Calibration curves are plotted with log(response ratio) vs log(concentration (pM)). Linear regression, limit of detection, and $R^2$ are reported in Table S3.

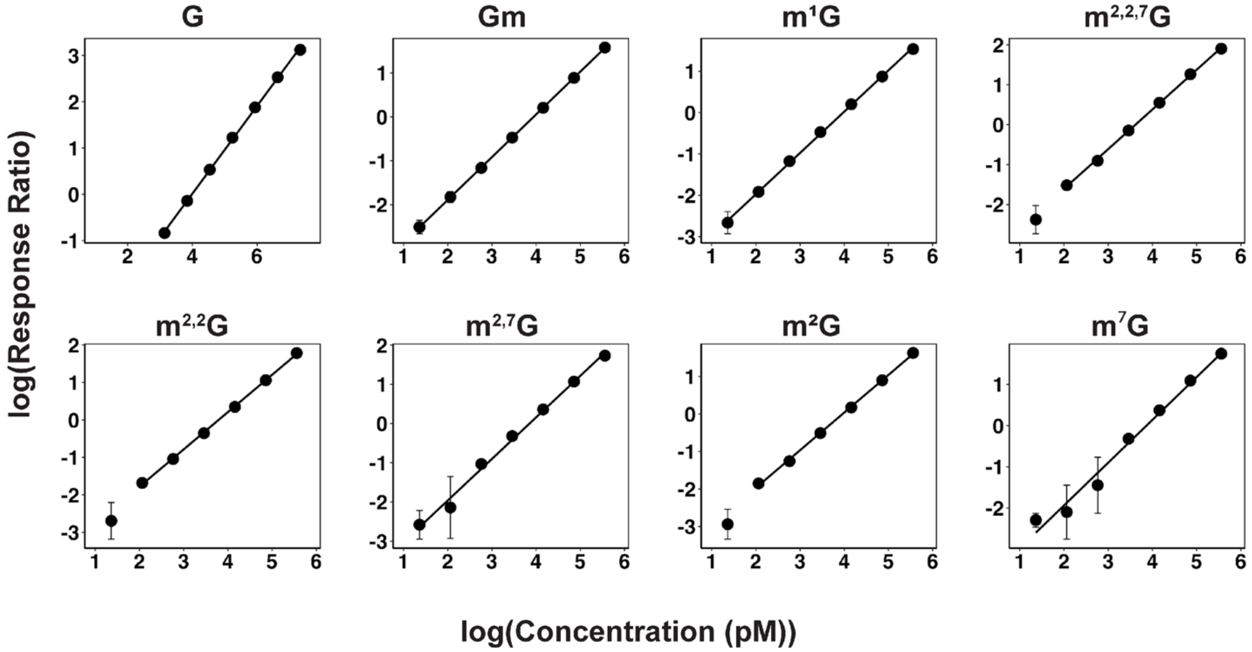


**Supplemental Figure S6.** Calibration curves used to quantify the concentration of guanosine modifications from set A. These data were run on the Agilent 6410 QQQ. Calibration curves are plotted with log(response ratio) vs log(concentration (pM)). Linear regression, limit of detection, and $R^2$ are reported in Table S3.

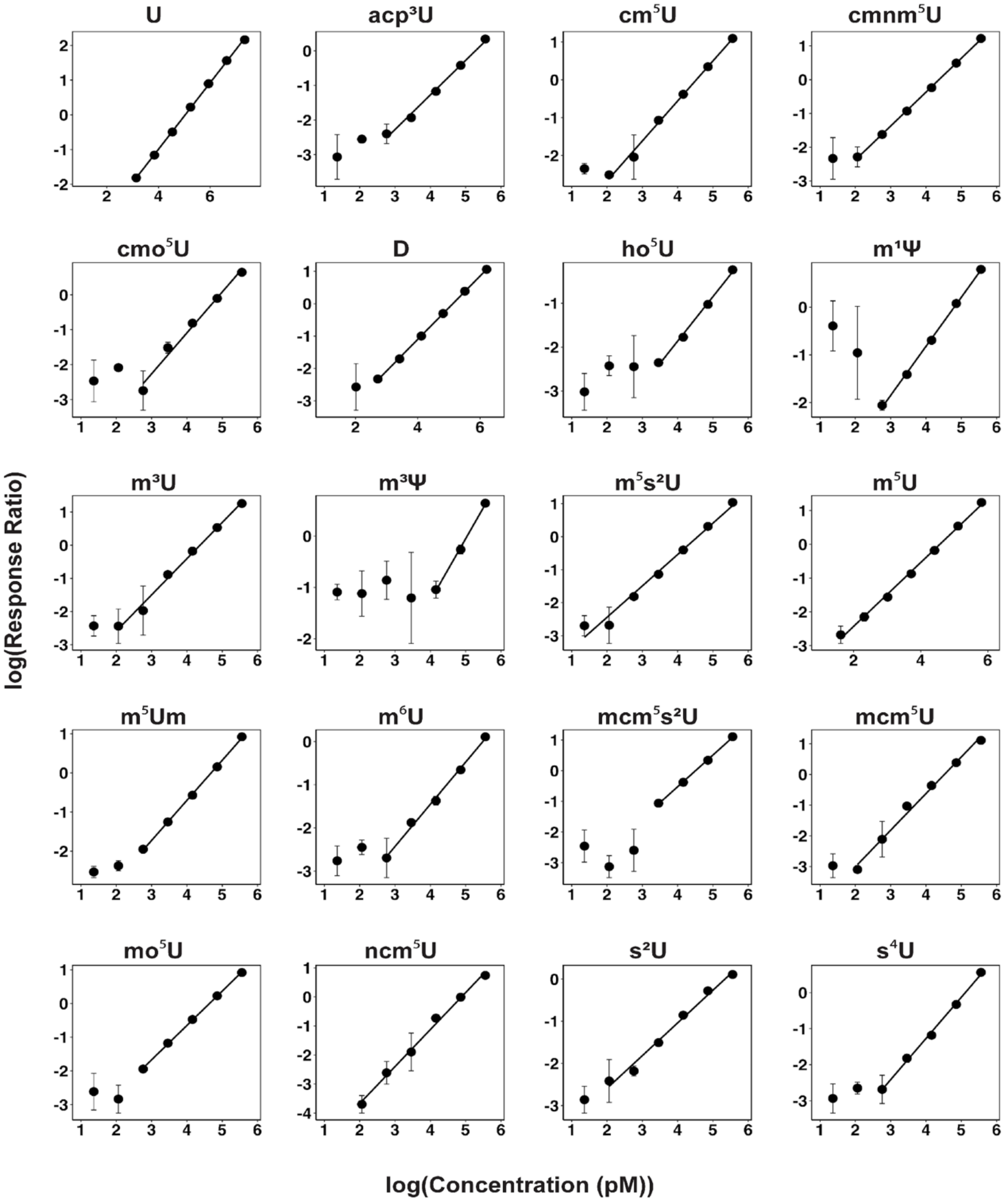


**Supplemental Figure S7.** Calibration curves used to quantify the concentration of uridine modifications from set A. These data were run on the Agilent 6410 QQQ. Calibration curves are plotted with log(response ratio) vs log(concentration (pM)). Linear regression, limit of detection, and $R^2$ are reported in Table S3.

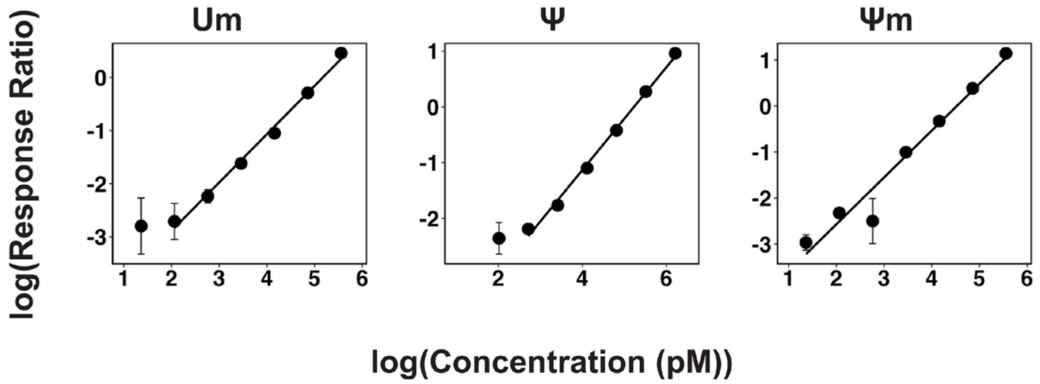


**Supplemental Figure S7(*cont.*).** Calibration curves used to quantify the concentration of uridine modifications from set A. These data were run on the Agilent 6410 QQQ. Calibration curves are plotted with log(response ratio) vs log(concentration (pM)). Linear regression, limit of detection, and $R^2$ are reported in Table S3.

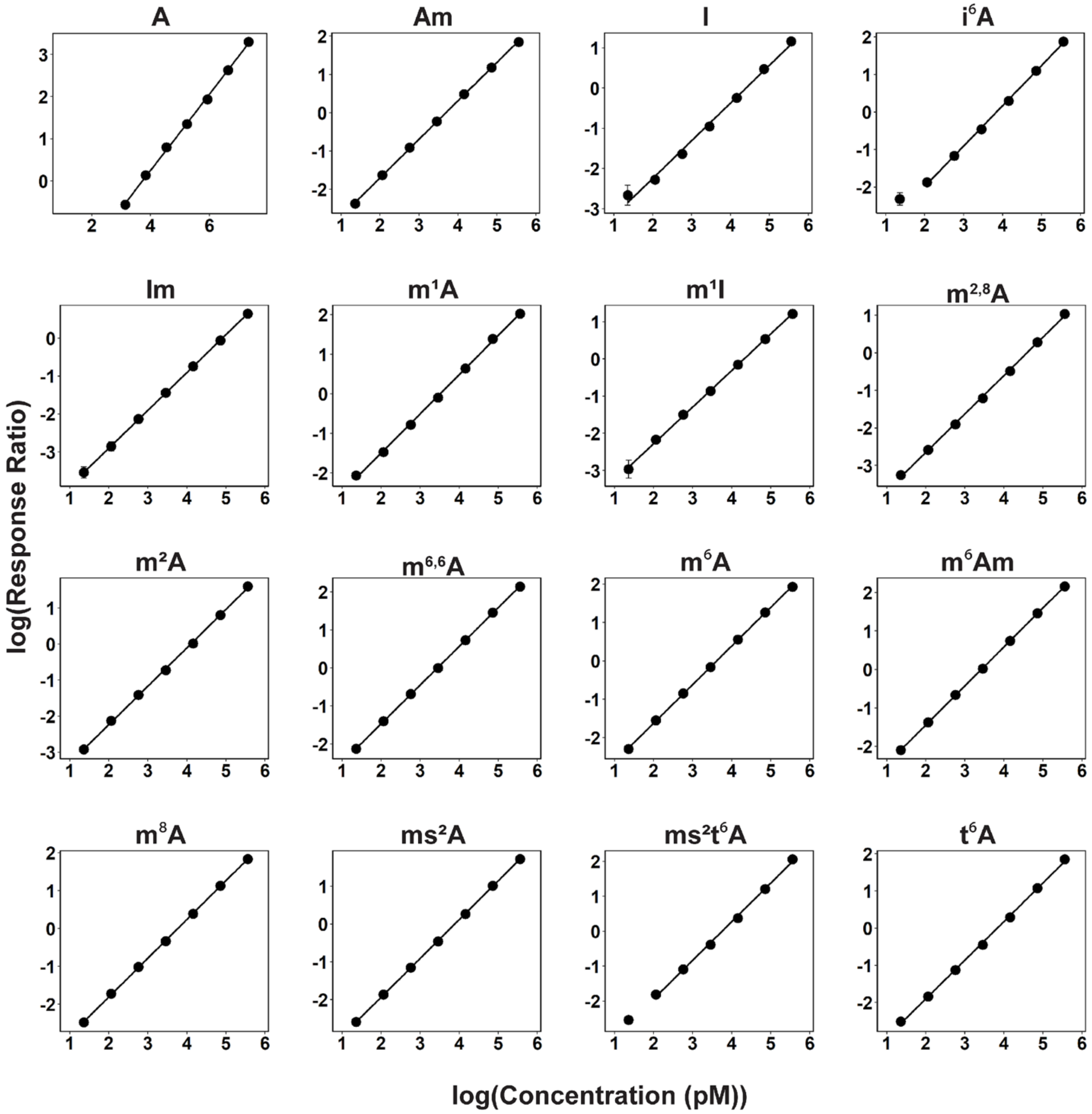


**Supplemental Figure S8.** Calibration curves used to quantify the concentration of adenosine modifications from set B. These data were run on the Agilent 6460 QQQ. Calibration curves are plotted with log(response ratio) vs log(concentration (pM)). Linear regression, limit of detection, and $R^2$ are reported in Table S4.

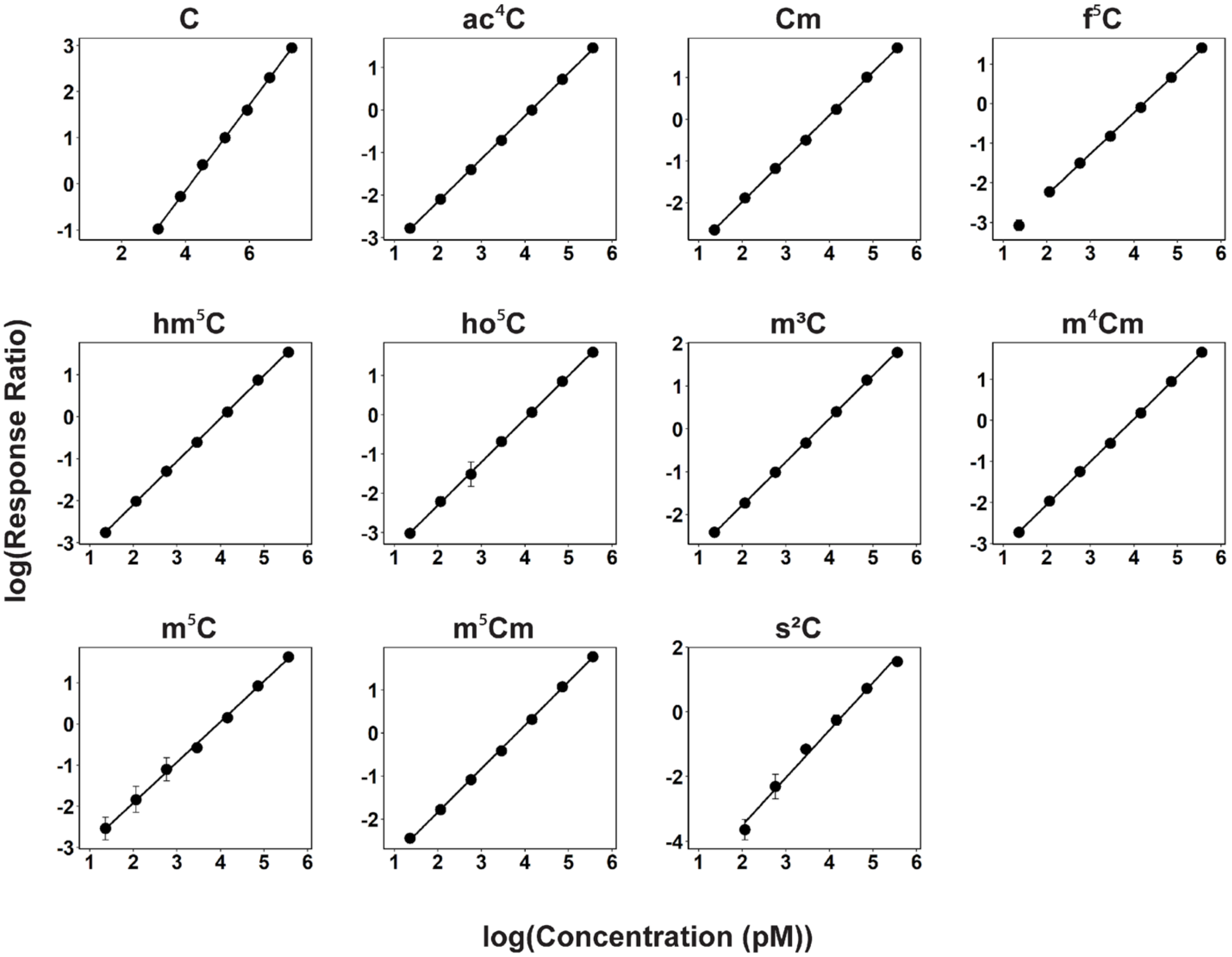


**Supplemental Figure S9.** Calibration curves used to quantify the concentration of cytidine modifications from set B. These data were run on the Agilent 6460 QQQ. Calibration curves are plotted with log(response ratio) vs log(concentration (pM)). Linear regression, limit of detection, and $R^2$ are reported in Table S4.

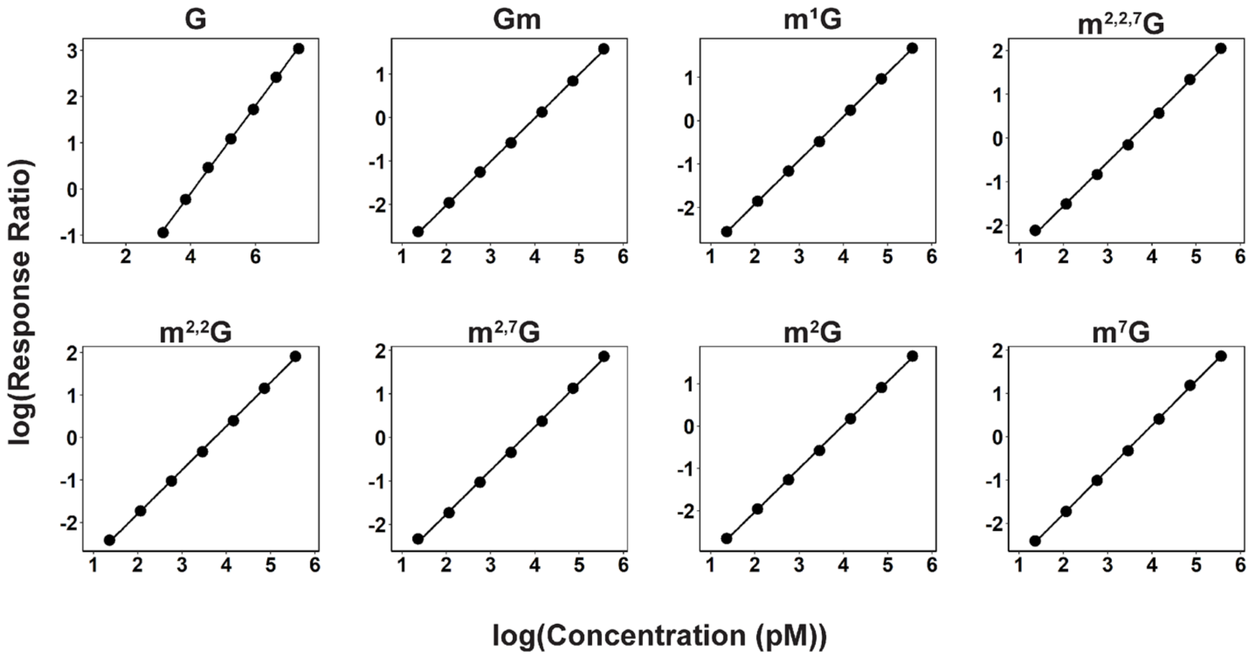


**Supplemental Figure S10.** Calibration curves used to quantify the concentration of guanosine modifications from set B. These data were run on the Agilent 6460 QQQ. Calibration curves are plotted with log(response ratio) vs log(concentration (pM)). Linear regression, limit of detection, and $R^2$ are reported in Table S4.

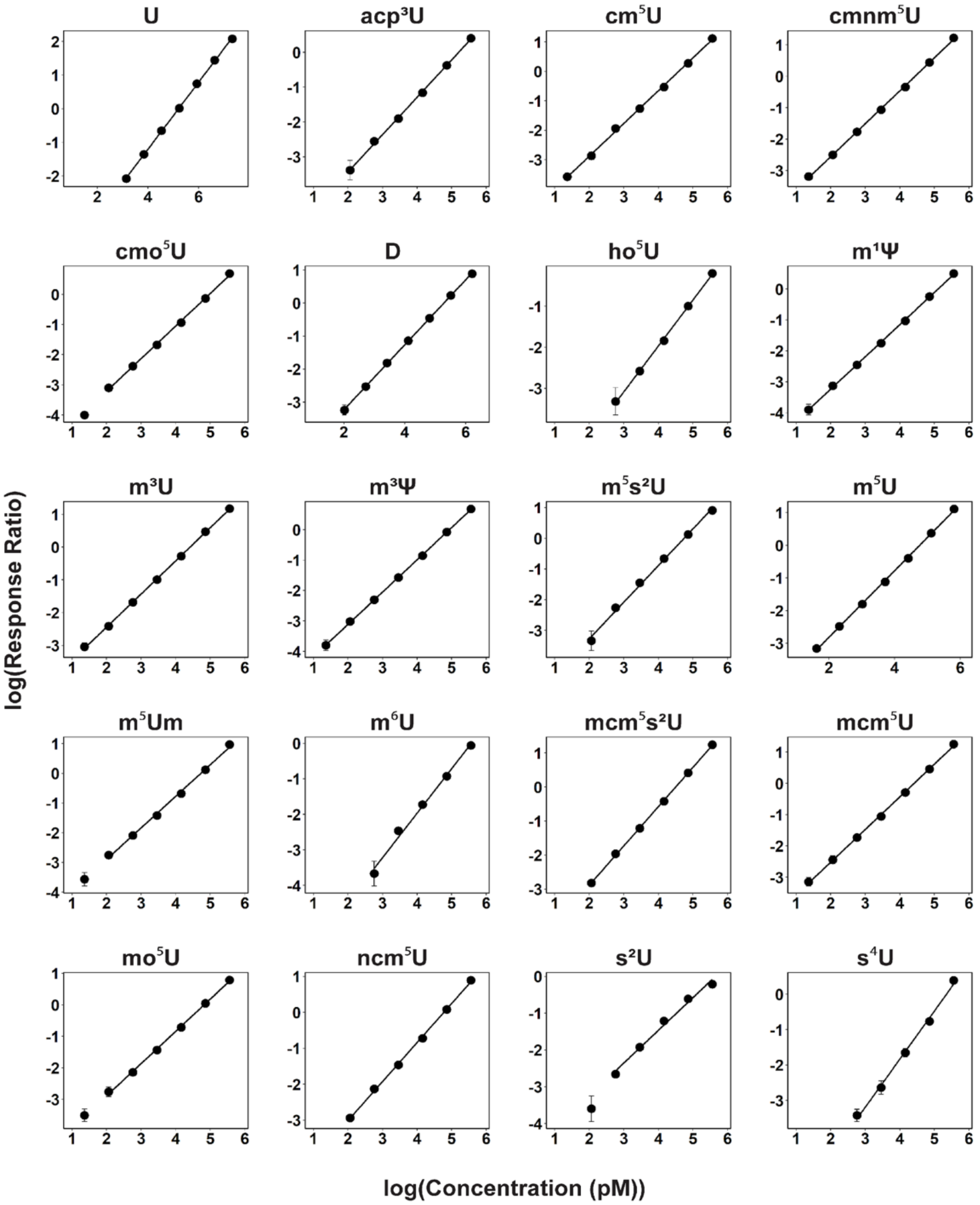


**Supplemental Figure S11.** Calibration curves used to quantify the concentration of uridine modifications from set B. These data were run on the Agilent 6460 QQQ. Calibration curves are plotted with log(response ratio) vs log(concentration (pM)). Linear regression, limit of detection, and $R^2$ are reported in Table S4.

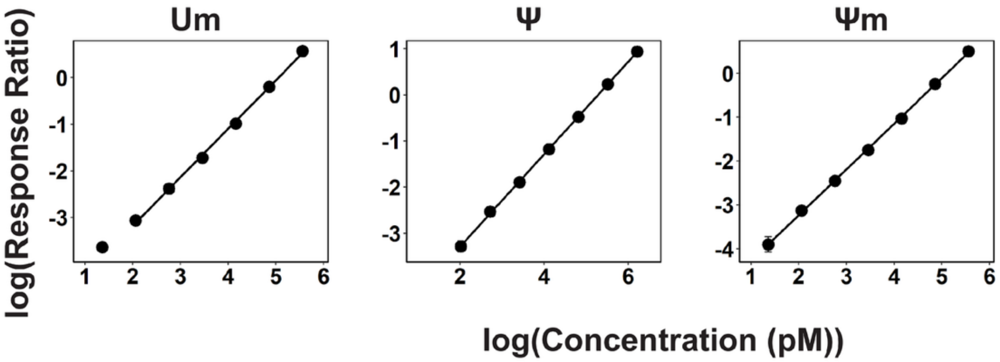


**Supplemental Figure S11 (*cont.*).** Calibration curves used to quantify the concentration of uridine modifications from set B. These data were run on the Agilent 6460 QQQ. Calibration curves are plotted with log(response ratio) vs log(concentration (pM)). Linear regression, limit of detection, and $R^2$ are reported in Table S4.

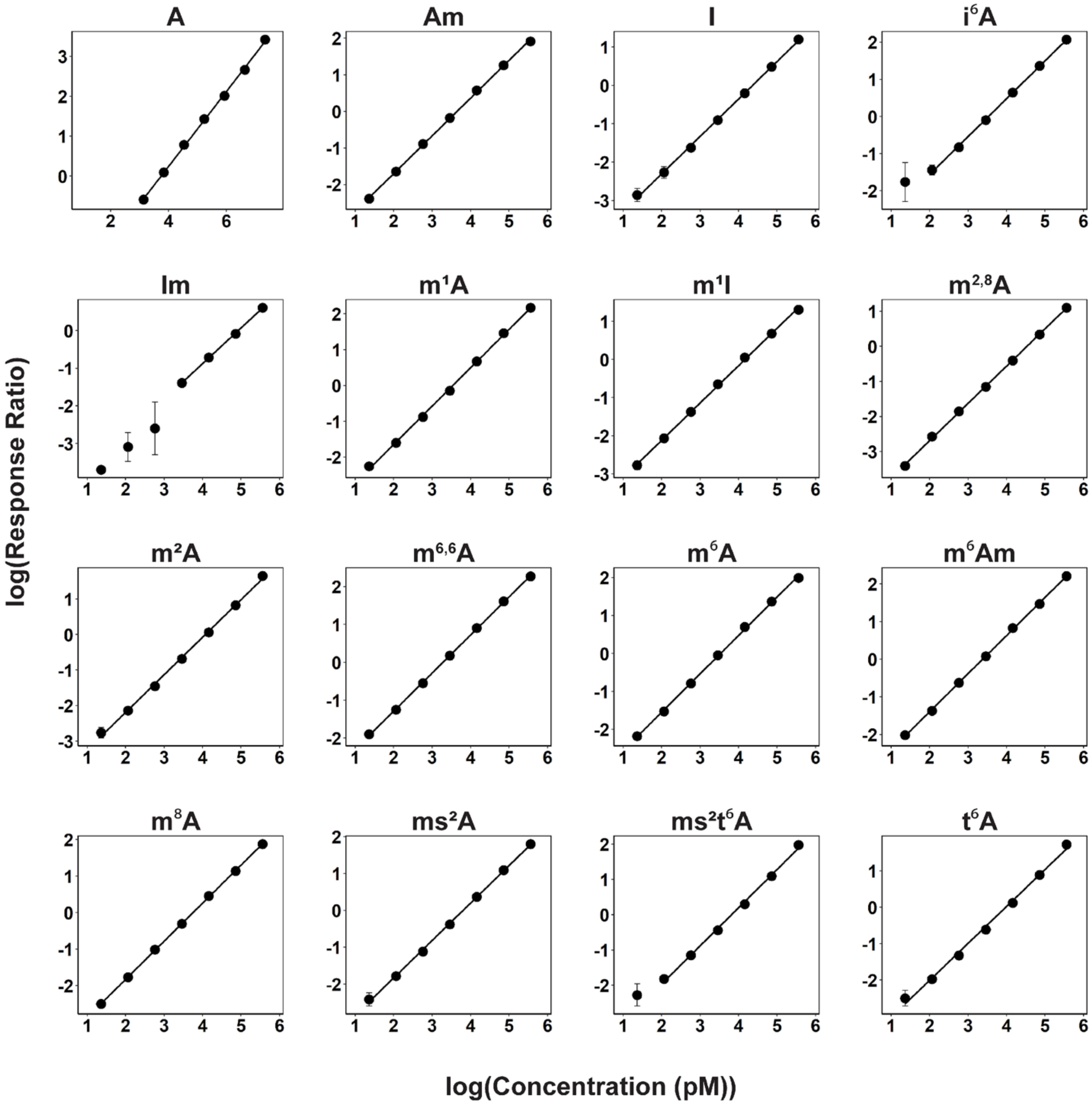


**Supplemental Figure S12.** Calibration curves used to quantify the concentration of adenosine modifications from set C. These data were run on the Agilent 6460 QQQ. Calibration curves are plotted with log(response ratio) vs log(concentration (pM)). Linear regression, limit of detection, and $R^2$ are reported in Table S5.

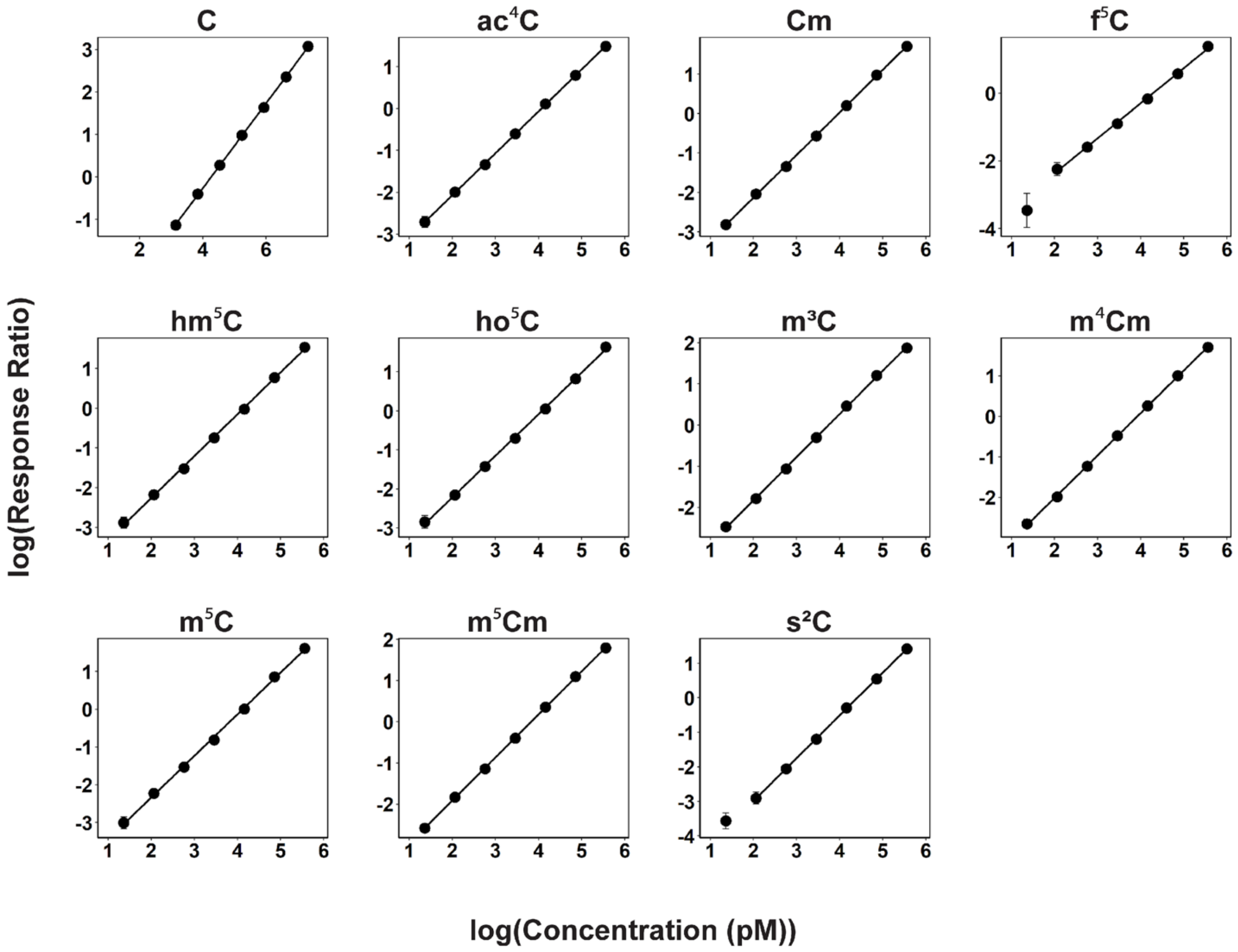


**Supplemental Figure S13.** Calibration curves used to quantify the concentration of cytidine modifications from set C. These data were run on the Agilent 6460 QQQ. Calibration curves are plotted with log(response ratio) vs log(concentration (pM)). Linear regression, limit of detection, and $R^2$ are reported in Table S5.

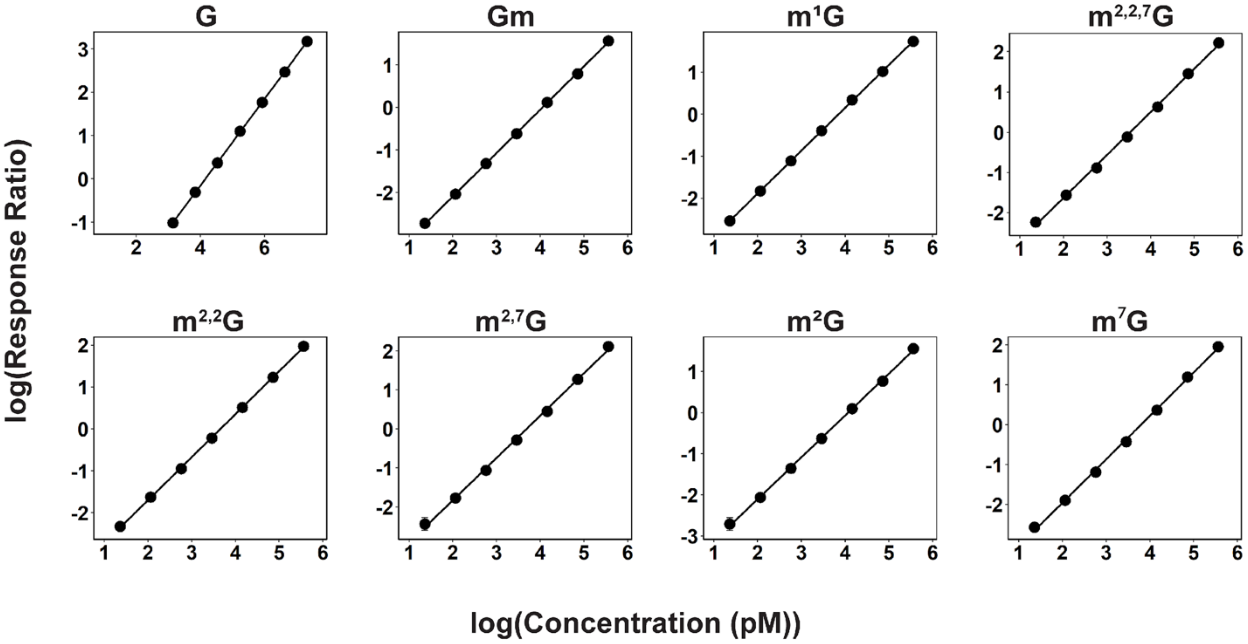


**Supplemental Figure S14.** Calibration curves used to quantify the concentration of guanosine modifications from set C. These data were run on the Agilent 6460 QQQ. Calibration curves are plotted with log(response ratio) vs log(concentration (pM)). Linear regression, limit of detection, and $R^2$ are reported in Table S5.

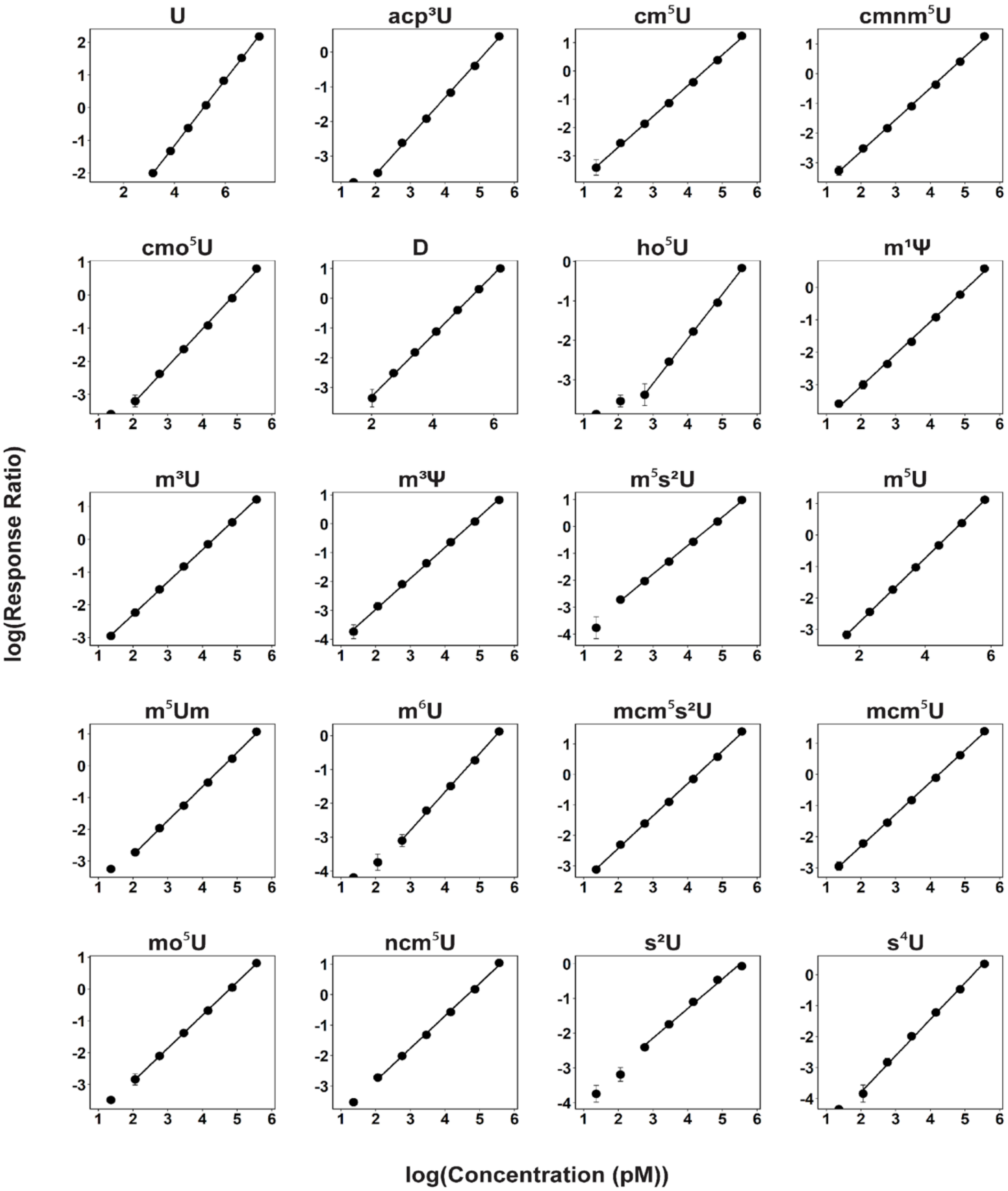


**Supplemental Figure S15.** Calibration curves used to quantify the concentration of uridine modifications from set C. These data were run on the Agilent 6460 QQQ. Calibration curves are plotted with log(response ratio) vs log(concentration (pM)). Linear regression, limit of detection, and $R^2$ are reported in Table S5.

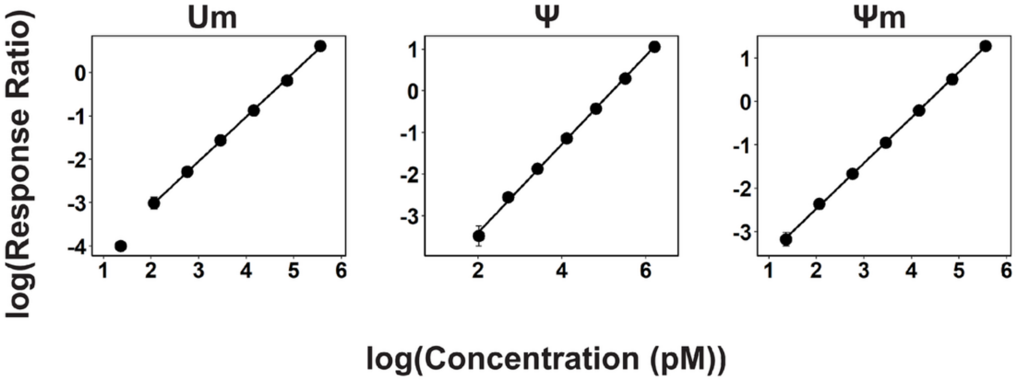


**Supplemental Figure S15 (*cont.*).** Calibration curves used to quantify the concentration of uridine modifications from set C. These data were run on the Agilent 6460 QQQ. Calibration curves are plotted with log(response ratio) vs log(concentration (pM)). Linear regression, limit of detection, and $R^2$ are reported in Table S5.

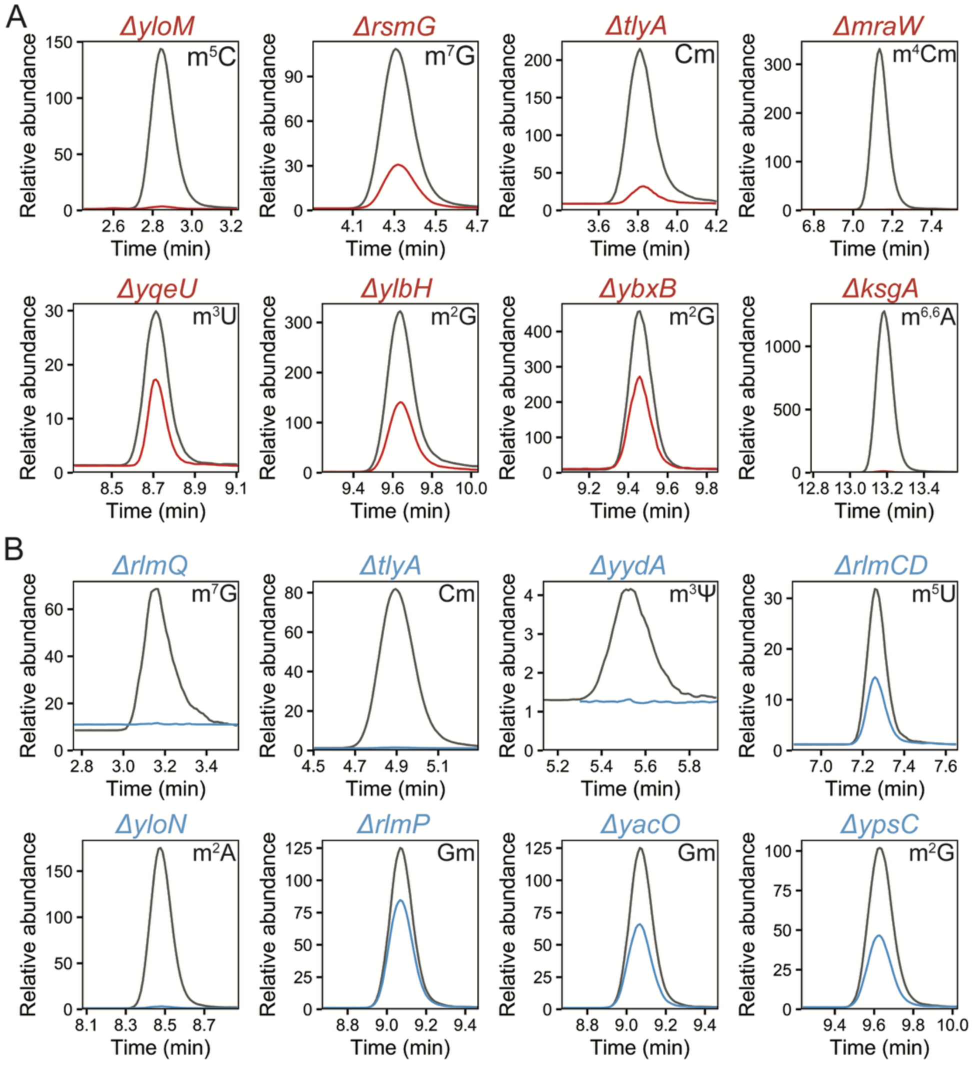


**Supplemental Figure S16.** Extracted ion chromatograms (EICs) of enzymes assigned to the modified nucleoside they install. Data were normalized to the maximum signal of the internal standard, and retention times were aligned to the wildtype sample. **A)** assigned enzymes and corresponding modifications in *B. subtilis* 16S rRNA. **B)** assigned enzymes and corresponding modifications in *B. subtilis* 23S rRNA.

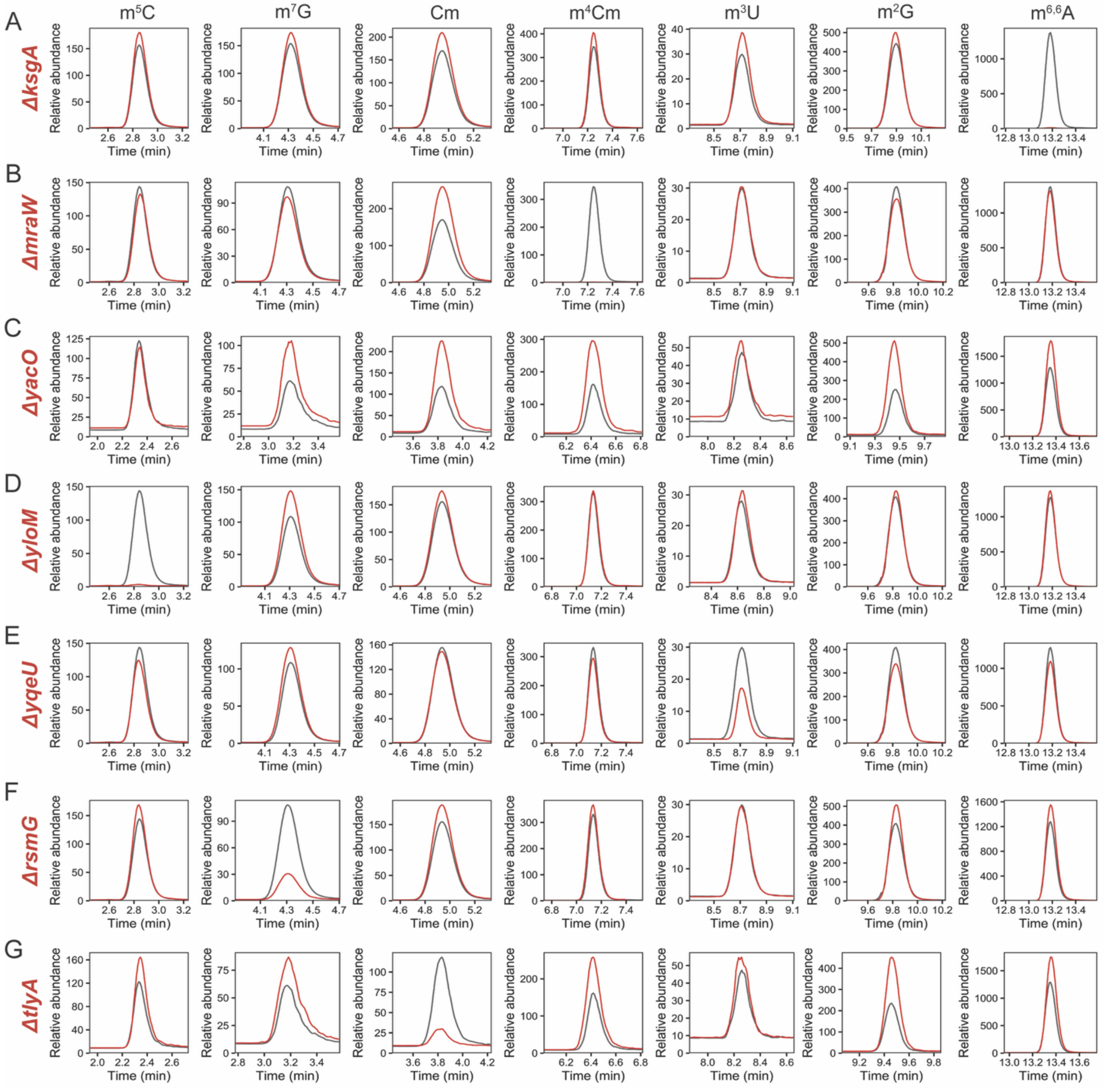


**Supplemental Figure S17.** EICs of methylated nucleosides in *B. subtilis* 16S rRNA across for profiled strains. Data were normalized to the maximum signal of the internal standard, and retention times were aligned to the wildtype sample. Wildtype rRNA nucleosides are shown in gray and mutant strains are shown in red. **A)** EICs from Δ*ksgA.* **B)** EICs from Δ*mraW.* **C)** EICs from Δ*yacO.* **D)** EICs from Δ*yloM.* **E)** EICs from Δ*yqeU.* **F)** EICs from Δ*rsmG.* **G)** EICs from Δ*tlyA.*

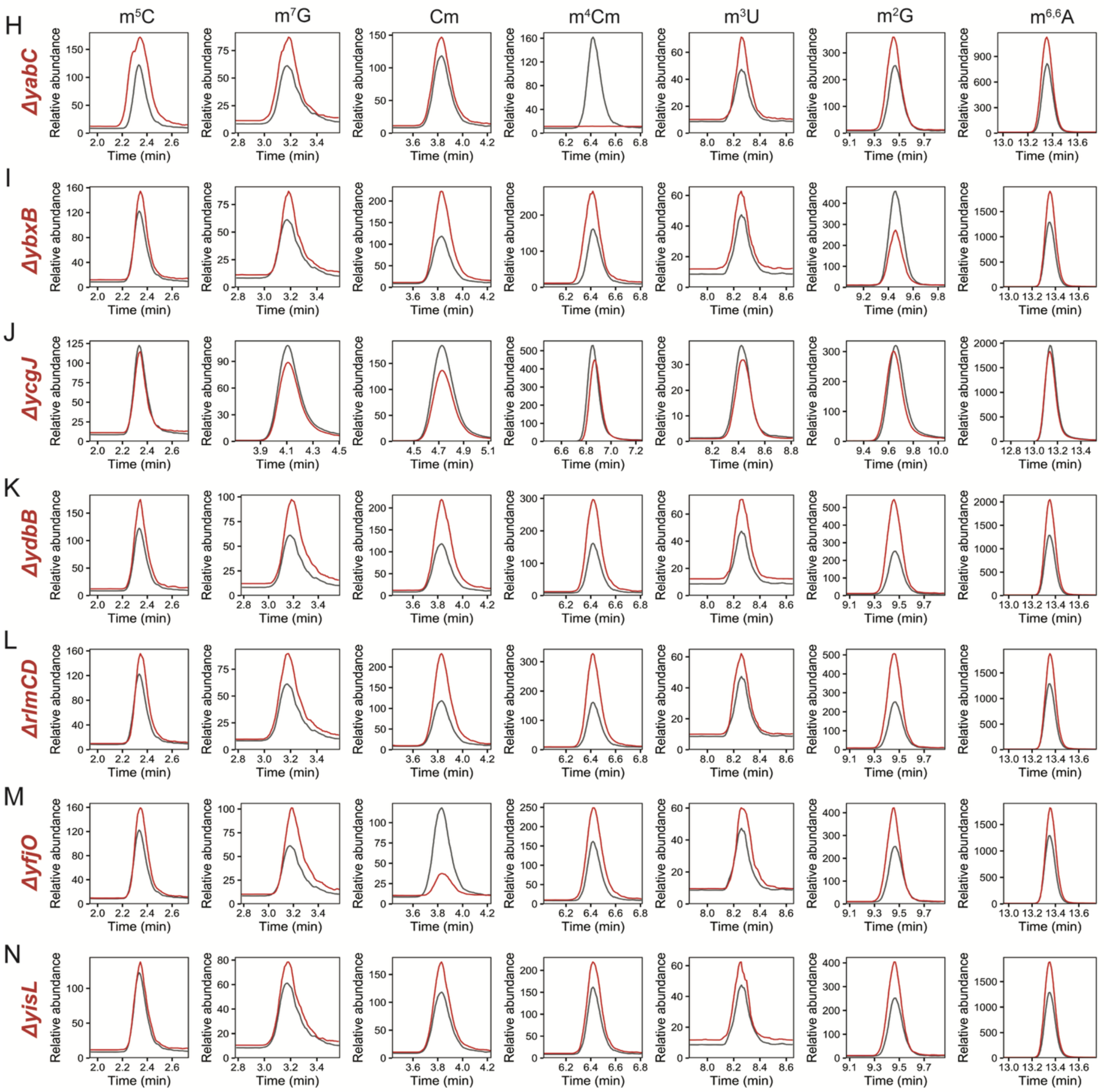

**Supplemental Figure S17 (*cont.*).** EICs of methylated nucleosides in *B. subtilis* 16S rRNA for profiled strains. Data were normalized to the maximum signal of the internal standard, and retention times were aligned to the wildtype sample. Wildtype rRNA nucleosides are shown in gray and mutant strains are shown in red. **H)** EICs from Δ*yabC.* **I)** EICs from Δ*ybxB.* **J)** EICs from Δ*ycgJ.* **K)** EICs from Δ*ydbB.* **L)** EICs from Δ*rlmCD.* **M)** EICs from Δ*yfjO.* **N)** EICs from Δ*yisL.*

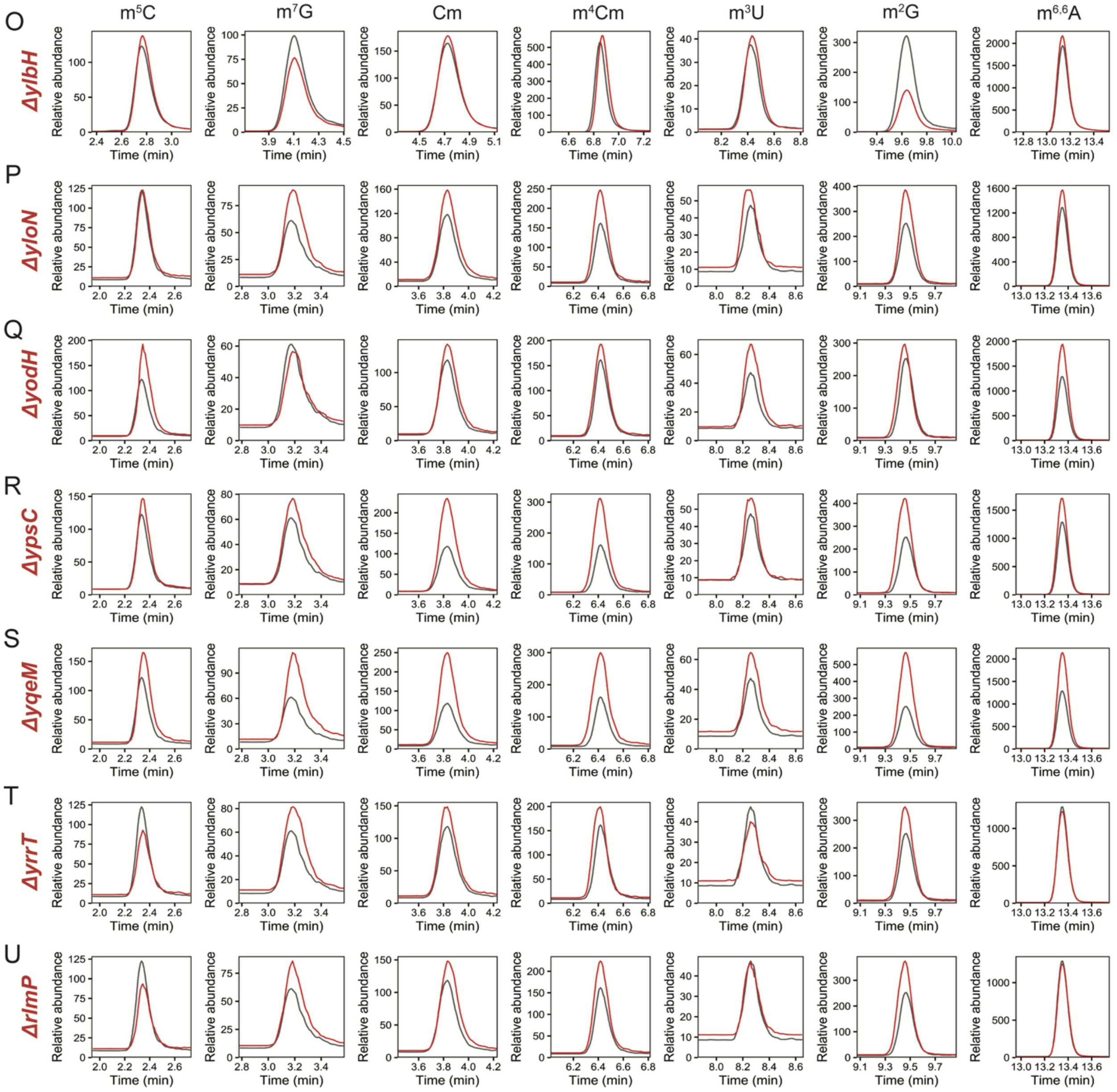


**Supplemental Figure S17 (*cont.*).** EICs of methylated nucleosides in *B. subtilis* 16S rRNA for profiled strains. Data were normalized to the maximum signal of the internal standard, and retention times were aligned to the wildtype sample. Wildtype rRNA nucleosides are shown in gray and mutant strains are shown in red. **O)** EICs from Δ*ylbH.* **P)** EICs from Δ*yloN.* **Q)** EICs from Δ*yodH.* **R)** EICs from Δ*ypsC.* **S)** EICs from Δ*yqeM.* **T)** EICs from Δ*yrrT.* **U)** EICs from Δ*rlmP.*

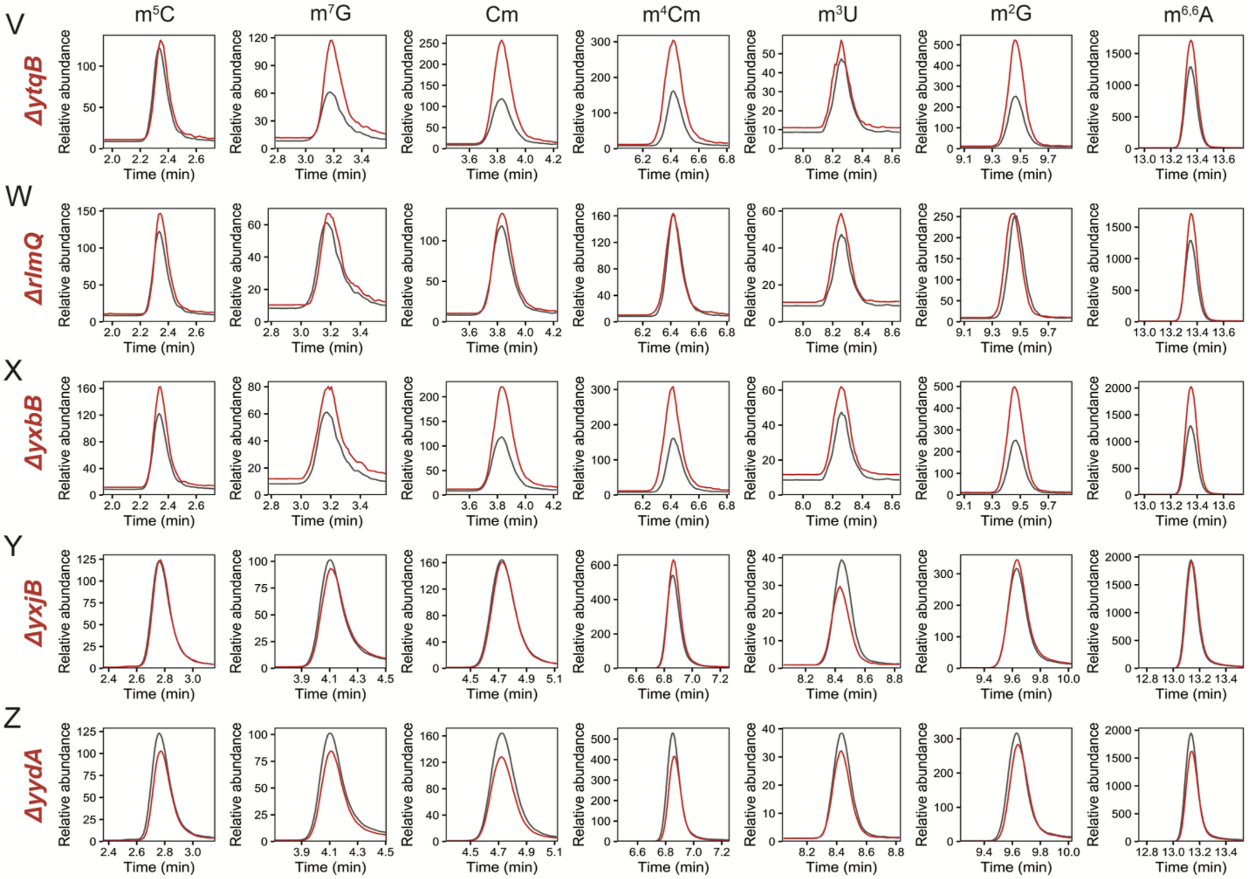


**Supplemental Figure S17 (*cont.*).** EICs of methylated nucleosides in *B. subtilis* 16S rRNA for profiled strains. Data were normalized to the maximum signal of the internal standard, and retention times were aligned to the wildtype sample. Wildtype rRNA nucleosides are shown in gray and mutant strains are shown in red. **V)** EICs from Δ*ytqB.* **W)** EICs from Δ*rlmQ.* **X)** EICs from Δ*yxbB.* **Y)** EICs from Δ*yxjB.* **Z)** EICs from Δ*yydA.*

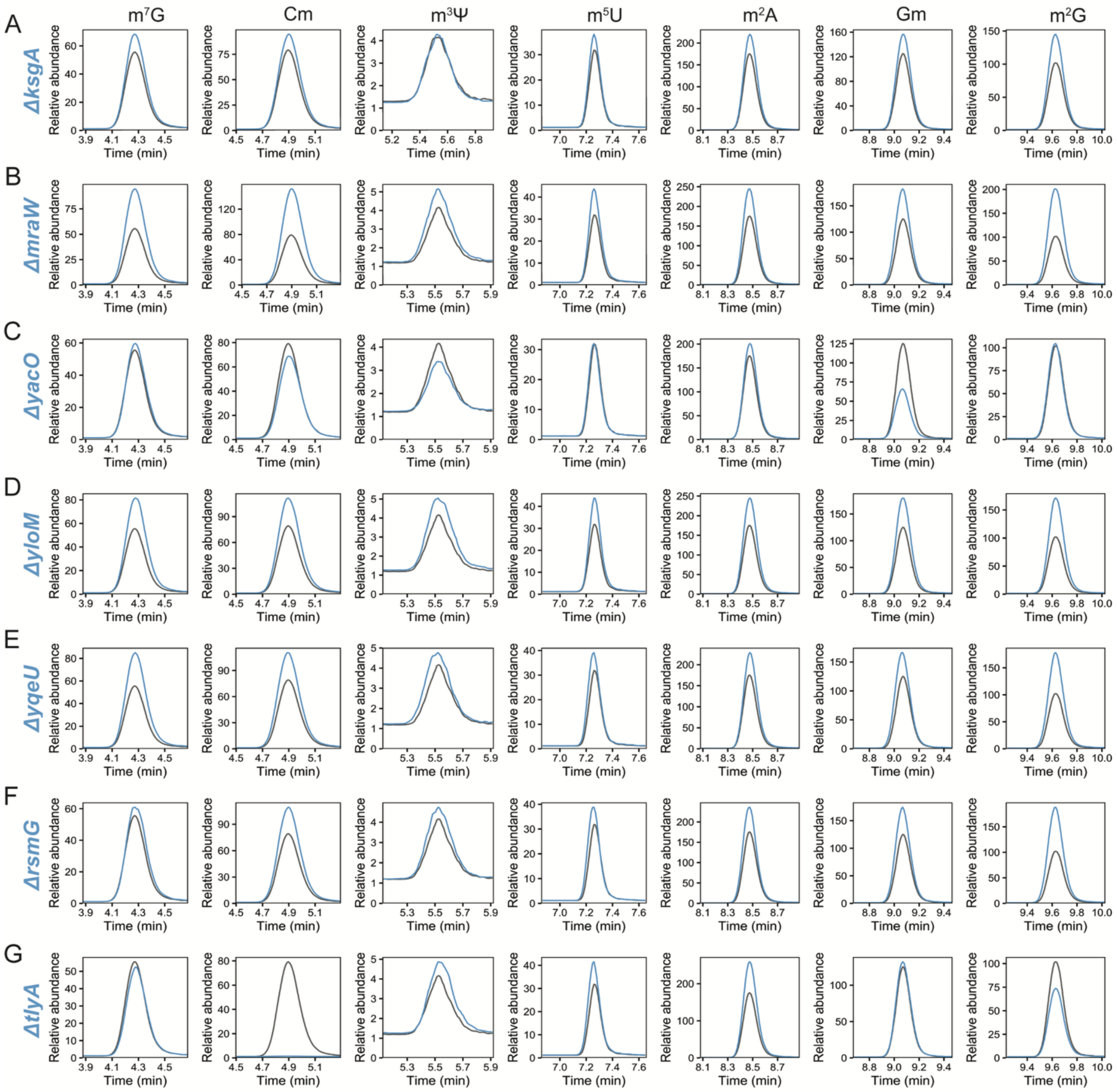


**Supplemental Figure S18.** EICs of methylated nucleosides in *B. subtilis* 23S rRNA for profiled strains. Data were normalized to the maximum signal of the internal standard, and retention times were aligned to the wildtype sample. Wildtype rRNA nucleosides are shown in gray and mutant rRNA nucleosides are shown in blue. Strains quantified from only calibration curve set A are missing an EIC because the signal for m$^3$Ψ was below the limit of detection (LOD). **A)** EICs from Δ*ksgA*. **B)** EICs from Δ*mraW*. **C)** EICs from Δ*yacO*. **D)** EICs from Δ*yloM*. **E)** EICs from Δ*yqeU*. **F)** EICs from Δ*rsmG*. **G)** EICs from Δ*tlyA*.

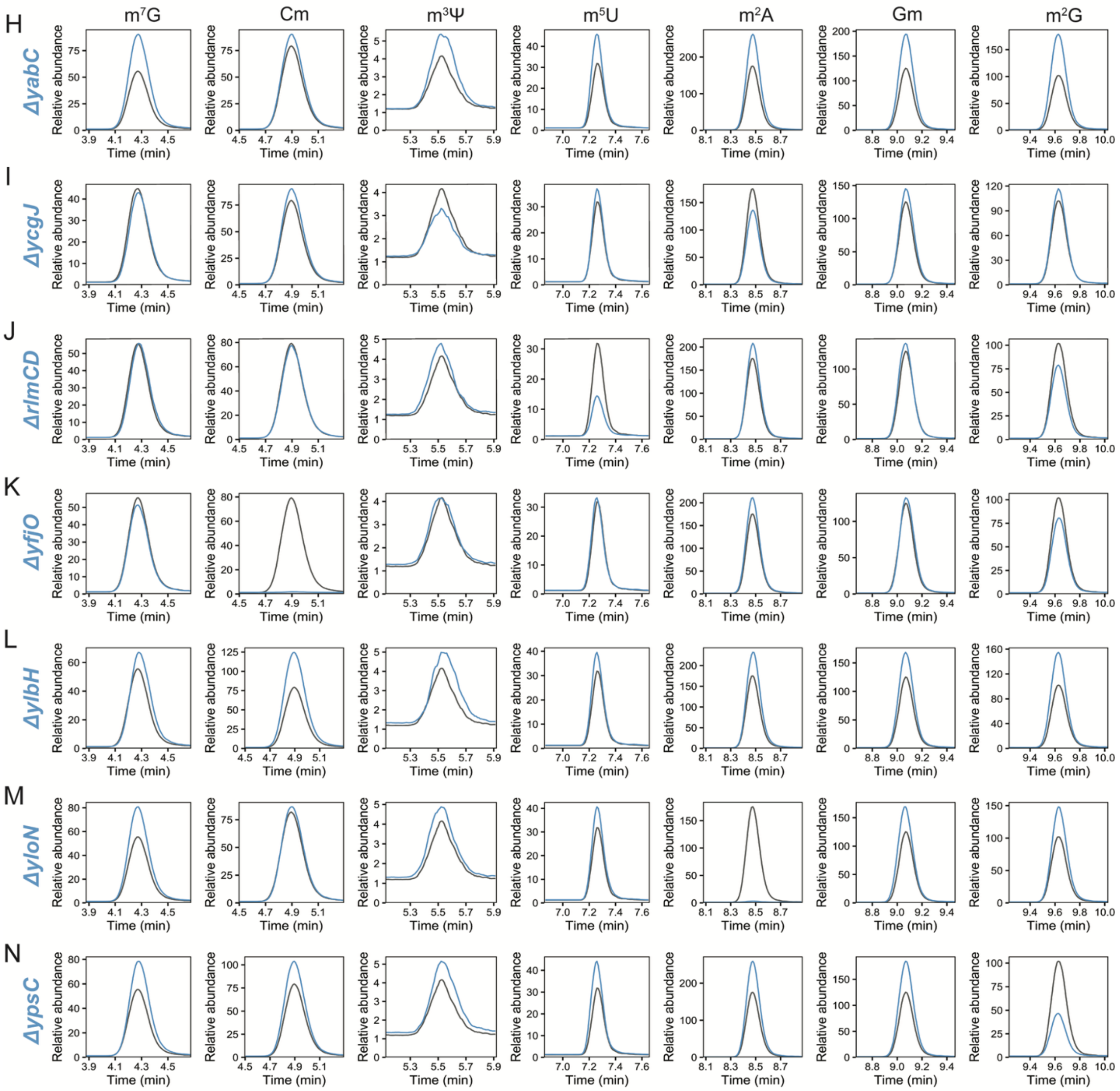


**Supplemental Figure S18 (*cont.*).** EICs of methylated nucleosides in *B. subtilis* 23S rRNA for profiled strains. Data were normalized to the maximum signal of the internal standard, and retention times were aligned to the wildtype sample. Wildtype rRNA nucleosides are shown in gray and mutant rRNA nucleosides are shown in blue. Strains quantified from only calibration curve set A are missing an EIC because the signal for $m^3\Psi$ was below the limit of detection (LOD). **H)** EICs from Δ*yabC*. **I)** EICs from Δ*ycgJ*. **J)** EICs from Δ*rlmCD*. **K)** EICs from Δ*yfjO*. **L)** EICs from Δ*ylbH*. **M)** EICs from Δ*yloN*. **N)** EICs from Δ*ypsC*.

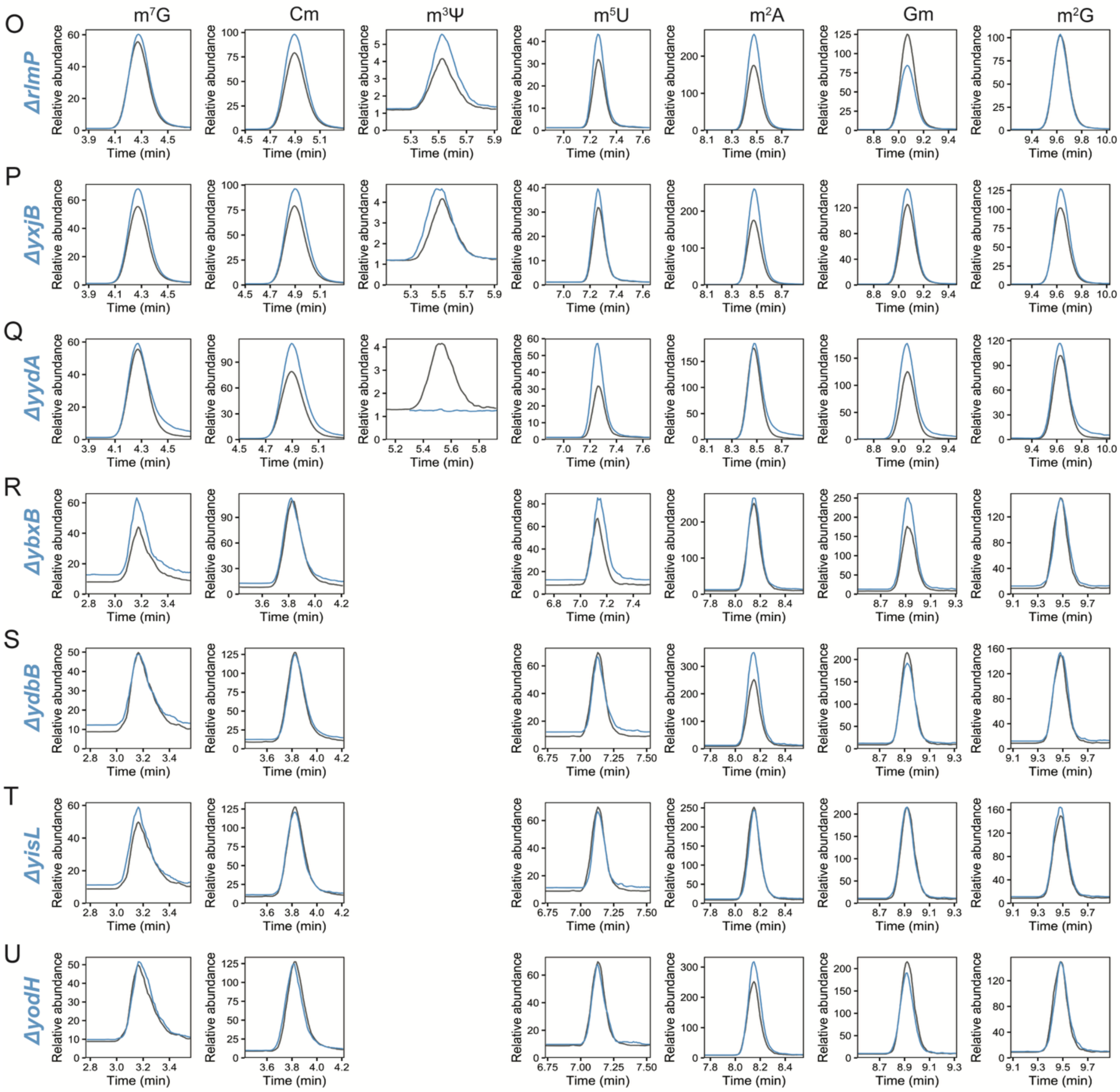


**Supplemental Figure S18 (*cont.*).** EICs of methylated nucleosides in *B. subtilis* 23S rRNA for profiled strains. Data were normalized to the maximum signal of the internal standard, and retention times were aligned to the wildtype sample. Wildtype rRNA nucleosides are shown in gray and mutant rRNA nucleosides are shown in blue. Strains quantified from only calibration curve set A are missing an EIC because the signal for $m^3\Psi$ was below the limit of detection (LOD). **O)** EICs from Δ*rlmP*. **P)** EICs from Δ*yxjB*. **Q)** EICs from Δ*yydA*. **R)** EICs from Δ*ybxB*. **S)** EICs from Δ*ydbB*. **T)** EICs from Δ*yisL*. **U)** EICs from Δ*yodH*.

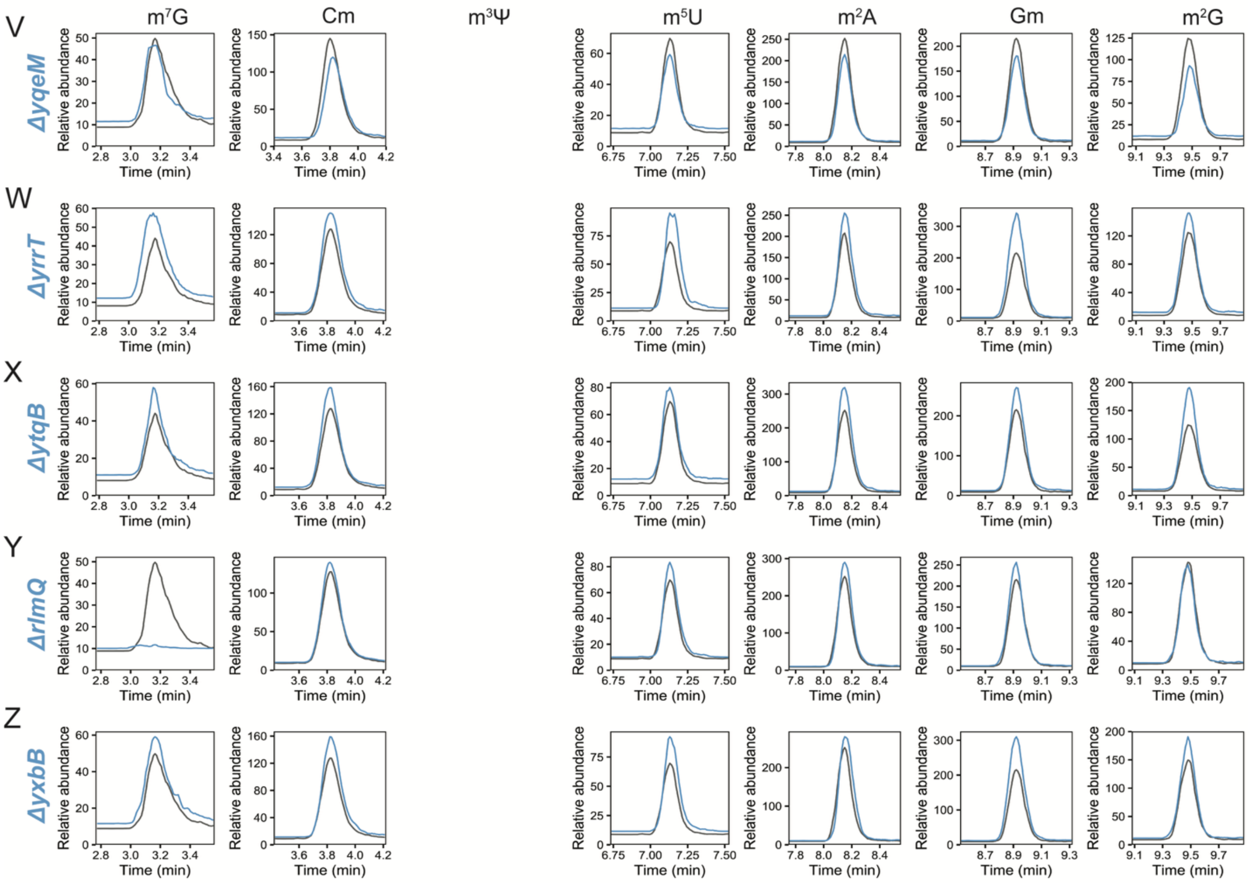


**Supplemental Figure S18 (*cont.*).** EICs of methylated nucleosides in *B. subtilis* 23S rRNA for profiled strains. Data were normalized to the maximum signal of the internal standard, and retention times were aligned to the wildtype sample. Wildtype rRNA nucleosides are shown in gray and mutant rRNA nucleosides are shown in blue. Strains quantified from only calibration curve set A are missing an EIC because the signal for $m^3\Psi$ was below the limit of detection (LOD). **V)** EICs from Δ*yqeM*. **W)** EICs from Δ*yrrT*. **X)** EICs from Δ*ytqB*. **Y)** EICs from Δ*rlmQ*. **Z)** EICs from Δ*yxbB*.

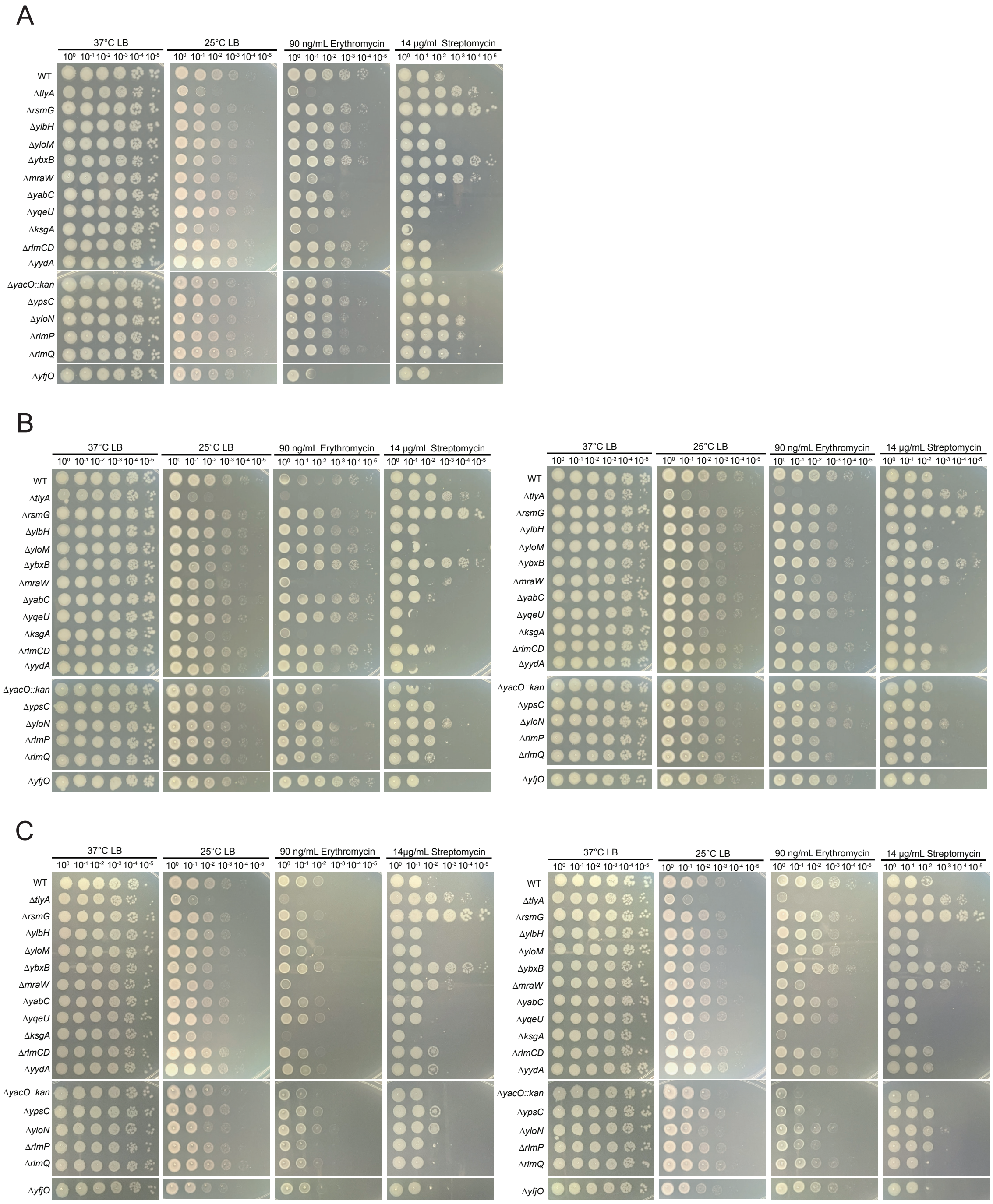


**Supporting Figure S19 Spot plate replicates.** Shown are three biological replicates in under the same stress conditions. **A)** One replicate is shown here the other is shown as Figure 4A in the main text. **B** and **C)** are biological replicates done in duplicate using the same conditions. In each panel all strains spotted for a given condition were spotted onto the same LB agar plate. The spot dilutions where then cropped from the same plate and arranged.

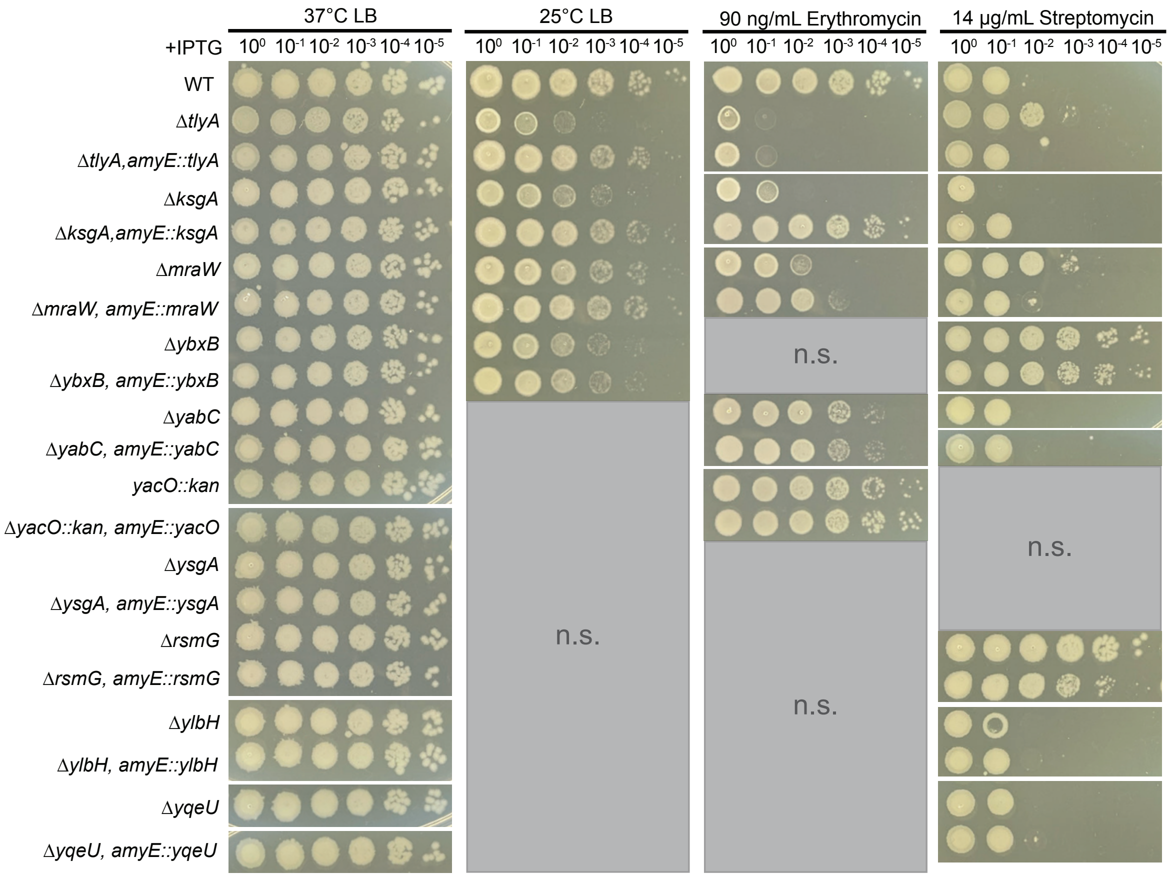


**Supporting Figure S20. Spot plate of genetic complementation (replicate #1).** Shown are the indicated strains with a complementation allele expressed ectopically from the *amyE* locus. Where deletion strains showed sensitive growth or resistance complementation was done to show that growth could be restored to the WT phenotype. Not Sensitive (N.S.) corresponds to gene deletions that did not differ from WT.

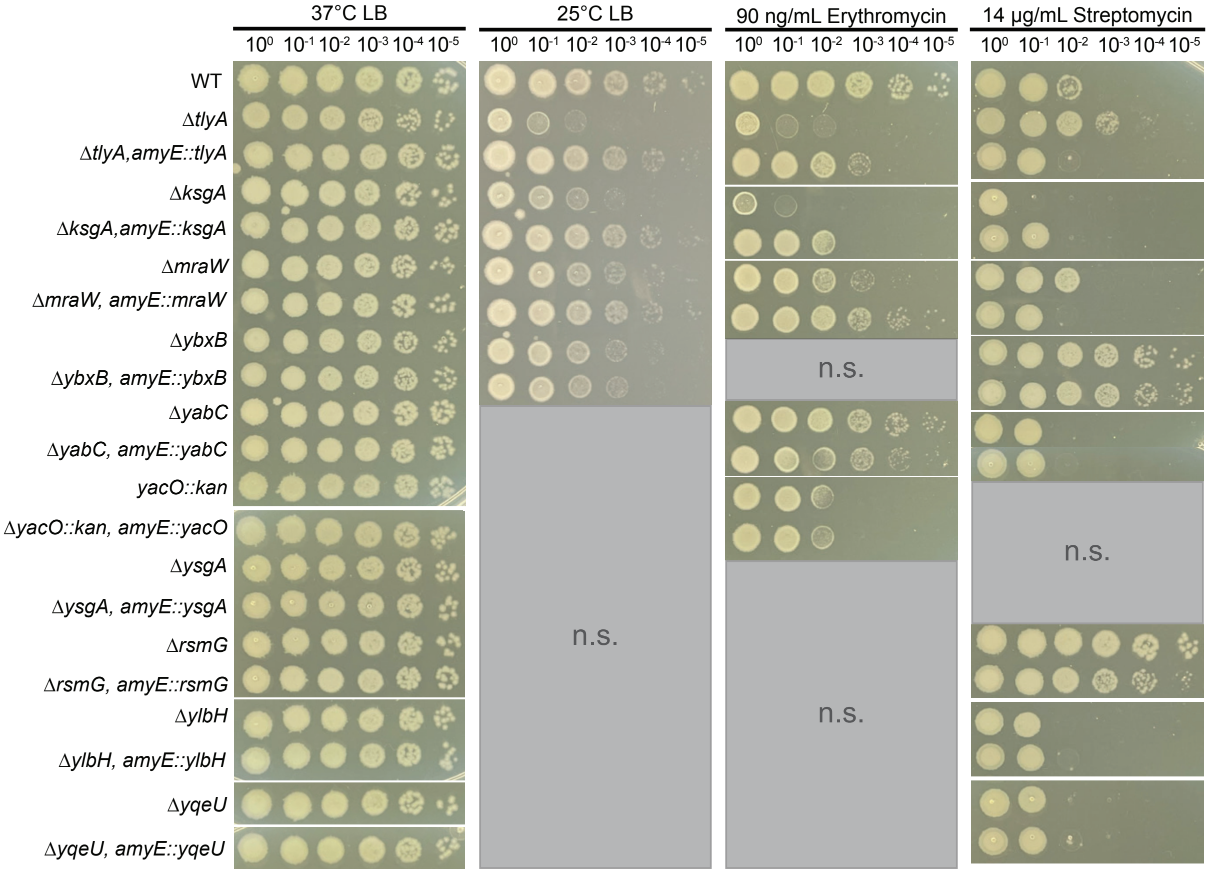


**Supporting Figure S20 continued. Spot plate of genetic complementation (replicate #2).** Shown are the indicated strains with a complementation allele expressed ectopically from the *amyE* locus. Where deletion strains showed sensitive growth or resistance complementation was done to show that growth could be restored to the WT phenotype. Not Sensitive (N.S.) corresponds to gene deletions that did not differ from WT.

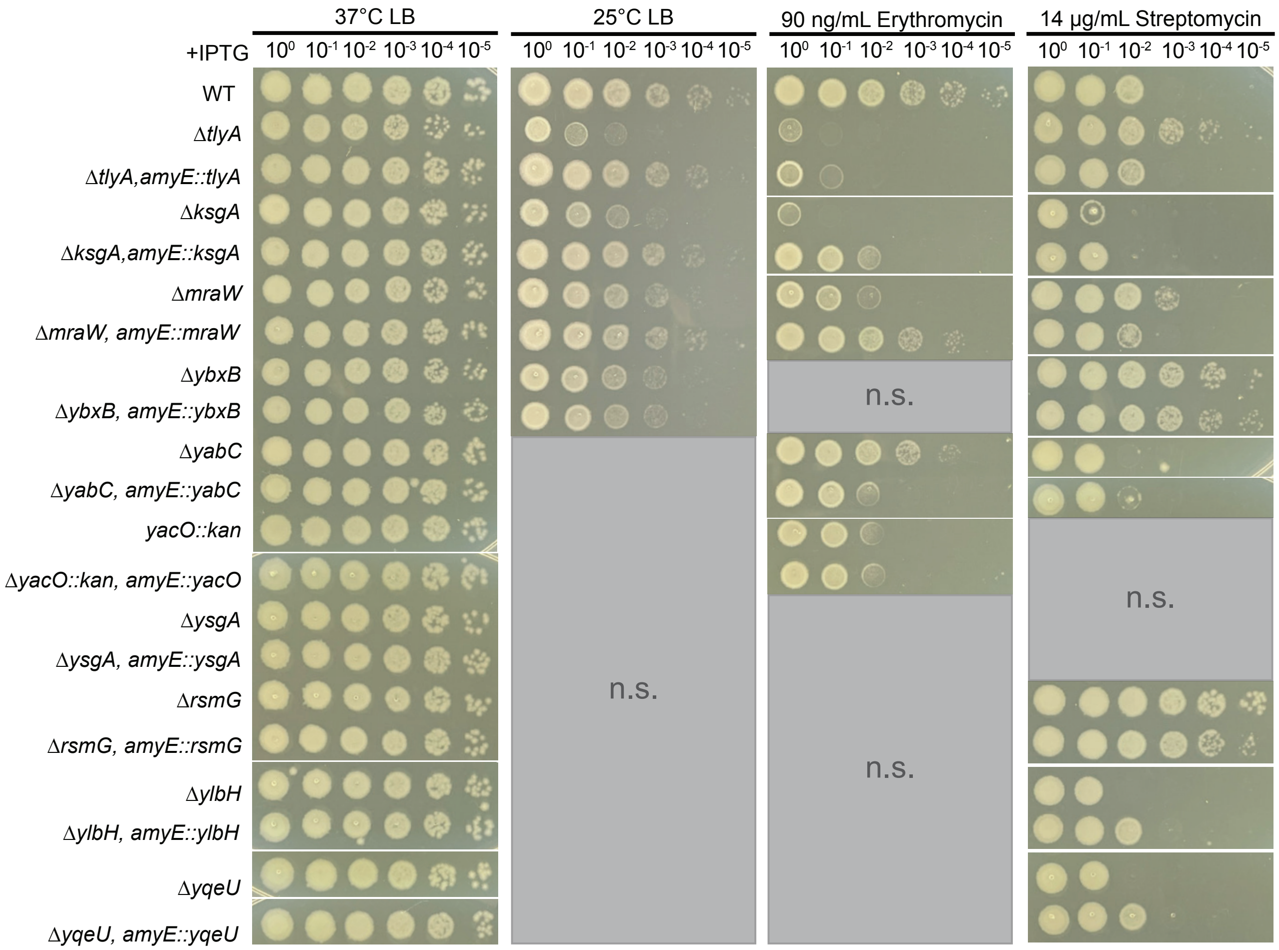


**Supporting Figure S20 continued. Spot plate of genetic complementation (replate #3)** Shown are the indicated strains with a complementation allele expressed ectopically from the *amyE* locus. Where deletion strains showed sensitive growth or resistance complementation was done to show that growth could be restored to the WT phenotype. Not Sensitive (N.S.) corresponds to gene deletions that did not differ from WT.

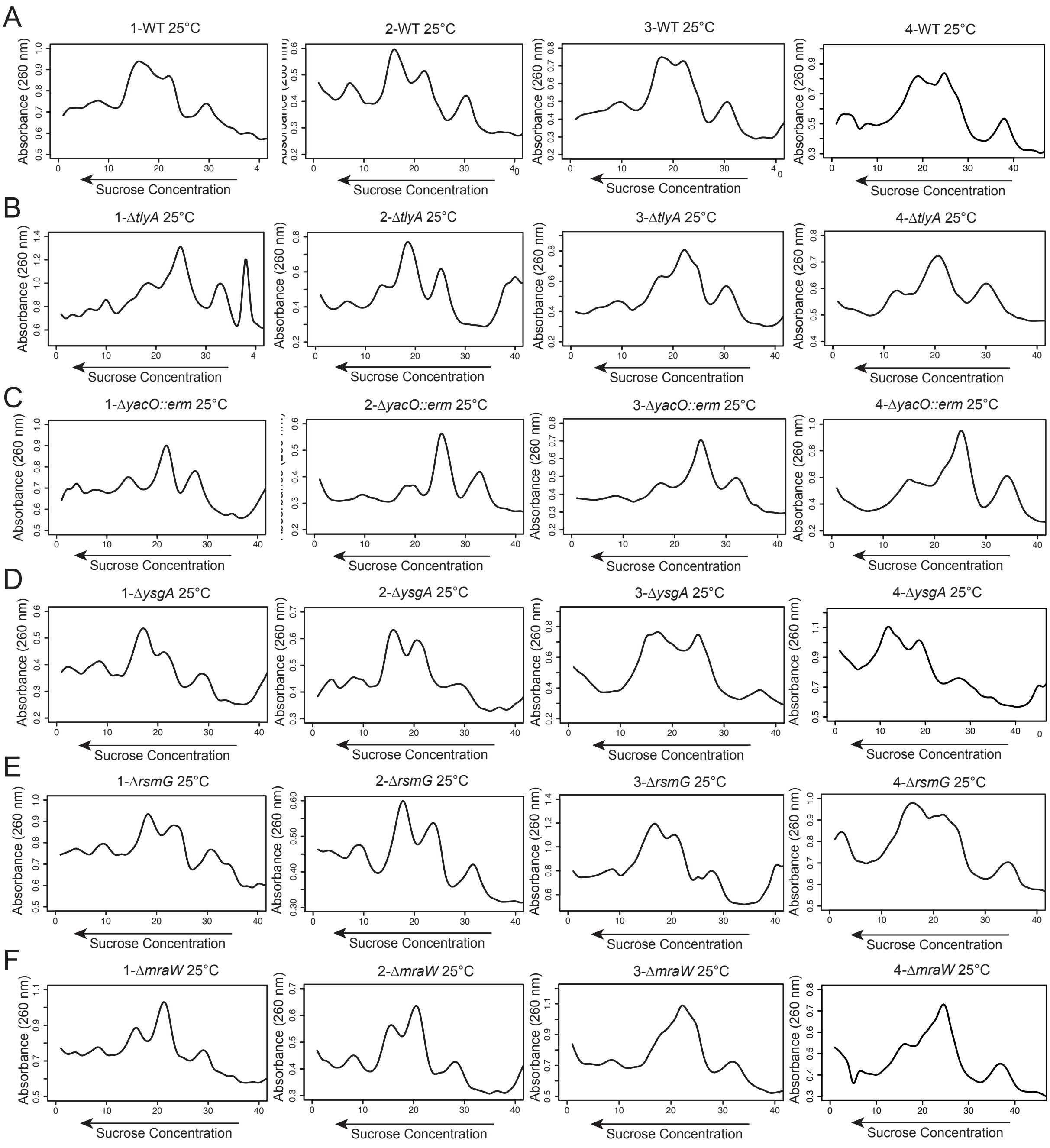


**Supplemental Figure S21. Replicates of sucrose gradients of ribosome assembly at 25ºC.** Shown are sucrose gradients to assess ribosome assembly from the indicated deletion mutants with cells grown at 25ºC under associating conditions. The samples are as follows: **A)** WT, **B)** Δ*tlyA*, **C)** Δ*yacO::erm*, **D)** Δ*ysgA*, **E)** Δ*rsmG*, **F)** Δ*mraW.*

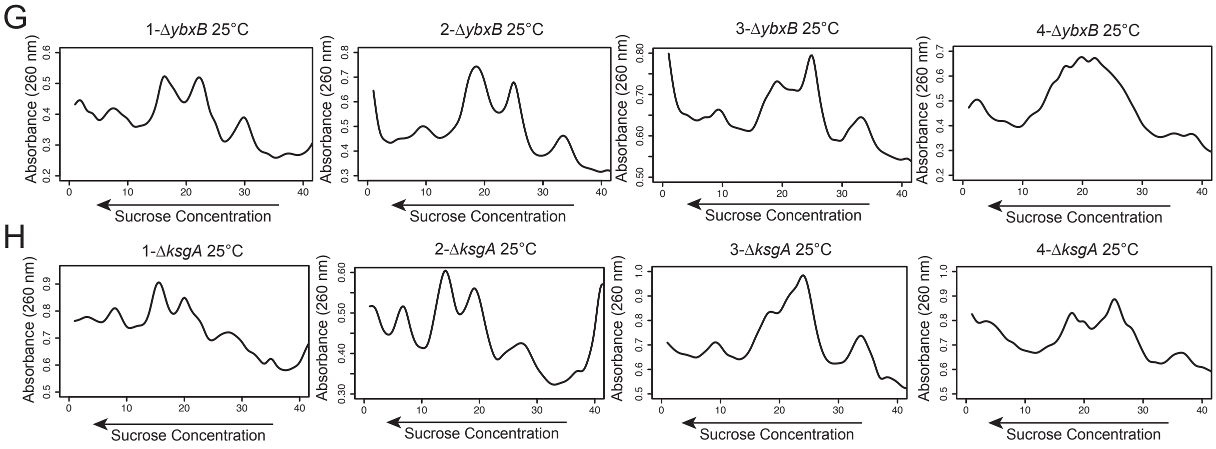


**Supplemental Figure S21 continued. Replicates of sucrose gradients of ribosome assembly at 25$^{o}$C.** Shown are sucrose gradients to assess ribosome assembly from the indicated deletion mutants with cells grown at 25$^{o}$C under associating conditions. The samples are as follows: **G)** Δ*ybxB*, **H)** Δ*ksgA.*

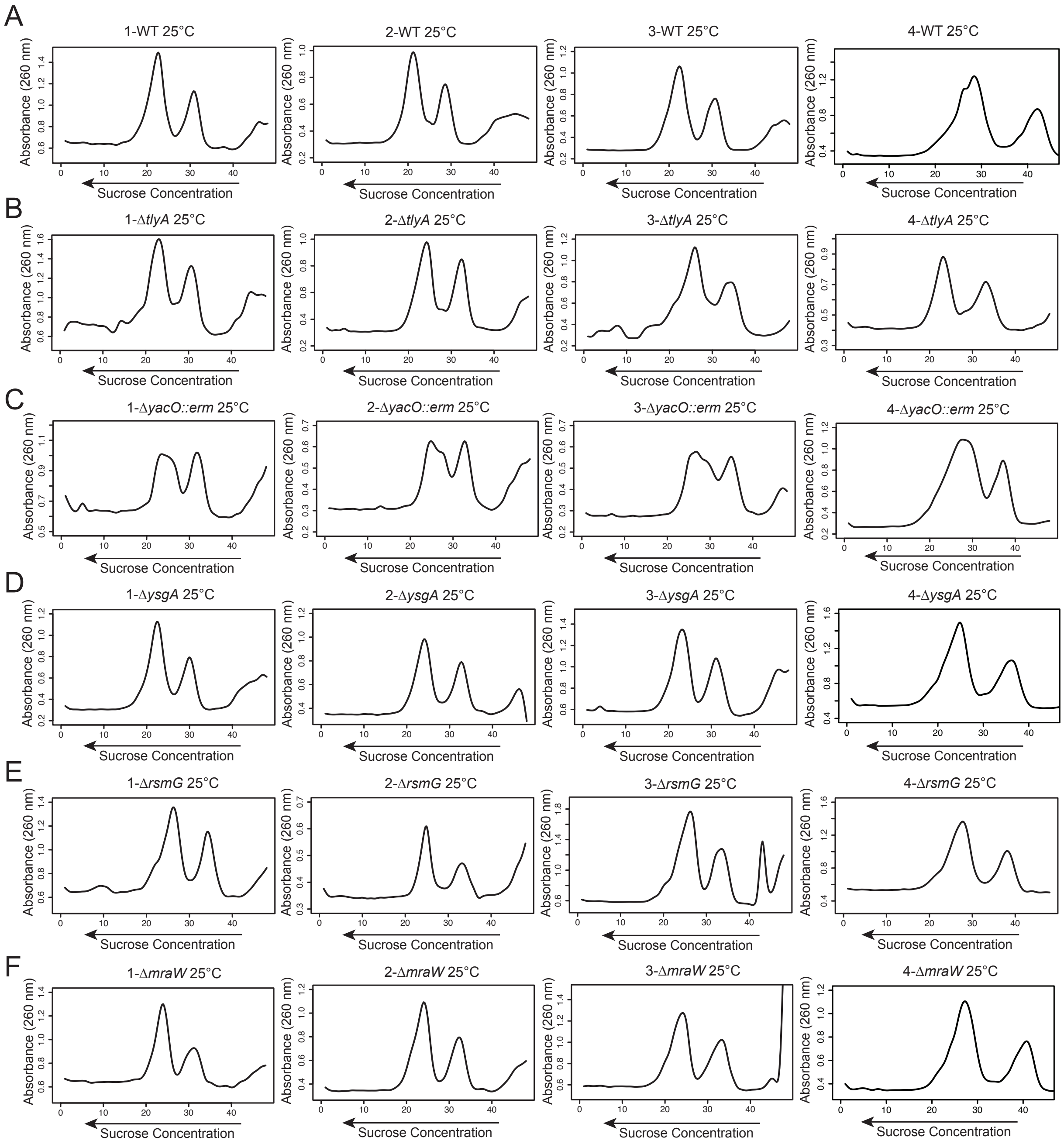


**Supplemental Figure S22. Replicates of sucrose gradients during dissociating conditions at 25ºC.** Shown are sucrose gradients to assess 50S and 30S subunit accumulation from the indicated deletion mutants with cells grown at 25ºC under dissociating conditions. The samples are as follows: **A)** WT, **B)** Δ*tlyA*, **C)** Δ*yacO::erm*, **D)** Δ*ysgA*, **E)** Δ*rsmG*, **F)** Δ*mraW*.

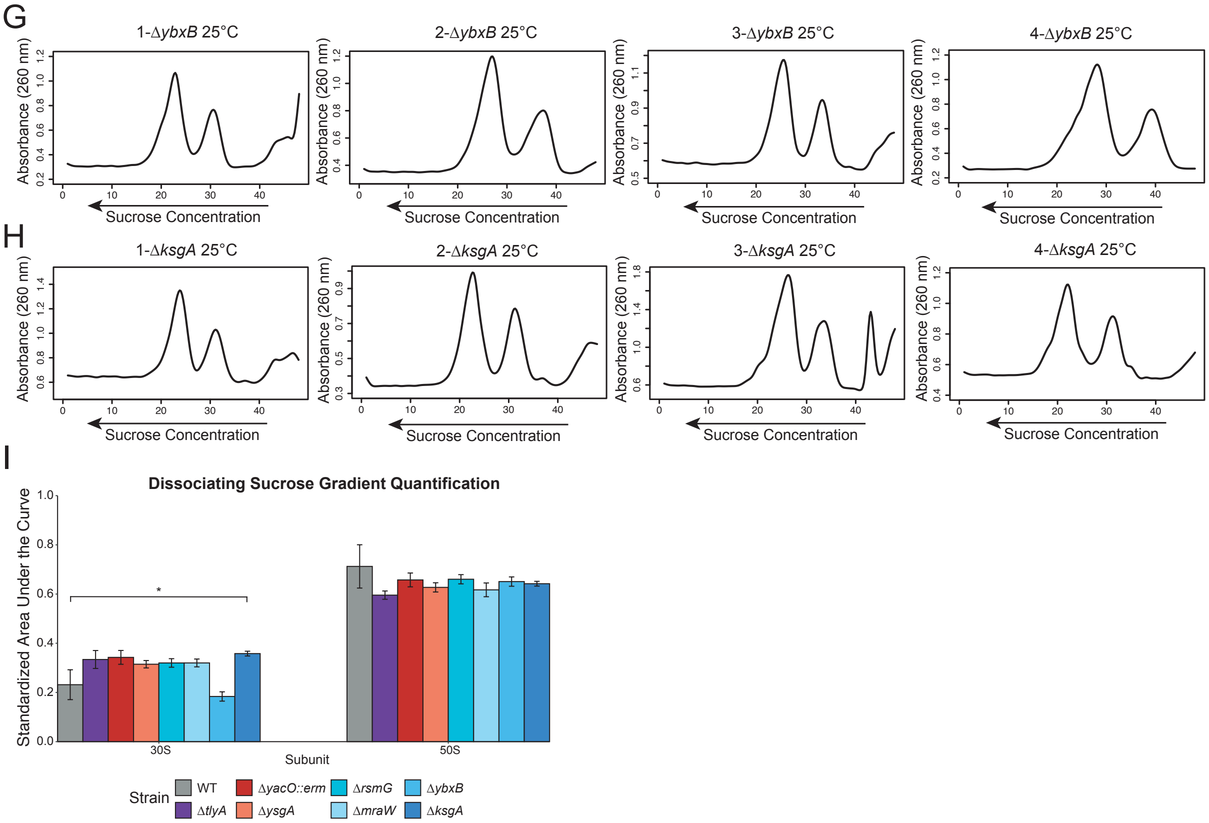


**Supplemental Figure S22 continued. Replicates of sucrose gradients during dissociating conditions at 25ºC.** Shown are sucrose gradients to assess 50S and 30S subunit accumulation from the indicated deletion mutants with cells grown at 25ºC under dissociating conditions. The samples are as follows: **G)** Δ*ybxB*, **H)** Δ*ksgA.* **I)** quantified area under the curve for 30S and 50S subunits. 50S were the same as wild type. We did detect differences in 30S accumulation with p<0.05*.

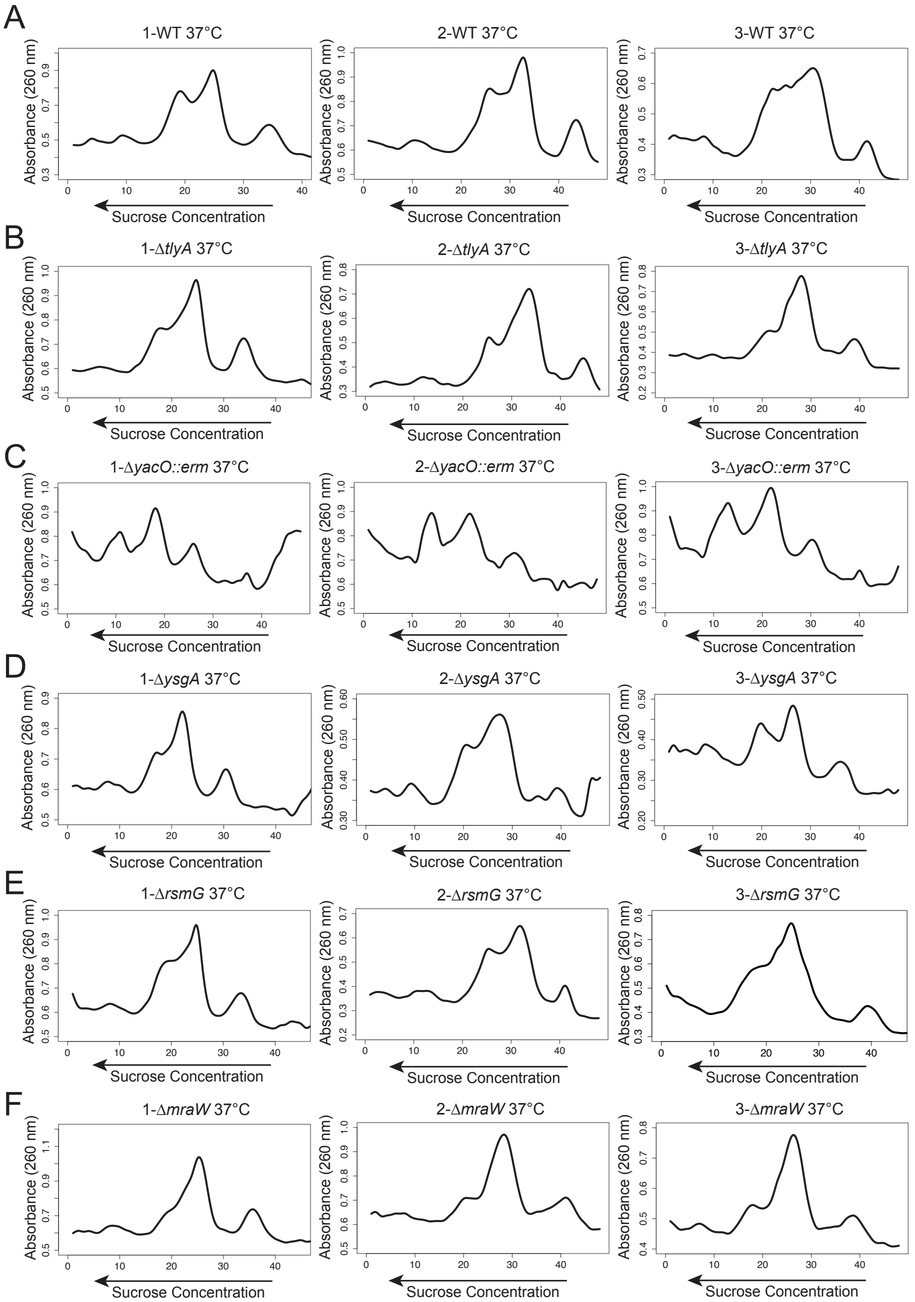


**Supplemental Figure S23. Replicates of sucrose gradients of ribosome assembly at 37°C.** Shown are sucrose gradients to assess ribosome assembly from the indicated deletion mutants with cells grown at 25°C under associating conditions. The samples are as follows: **A)** WT, **B)** Δ*tlyA*, **C)** Δ*yacO::erm*, **D)** Δ*ysgA*, **E)** Δ*rsmG*, **F)** Δ*mraW.*

G

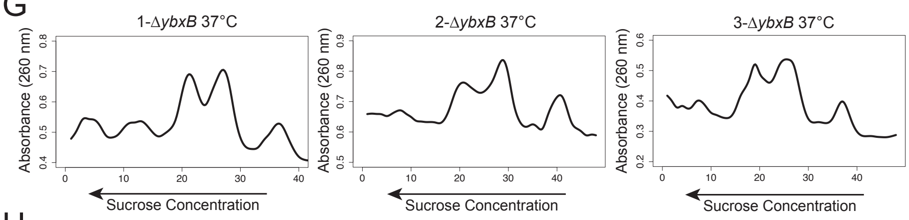


H

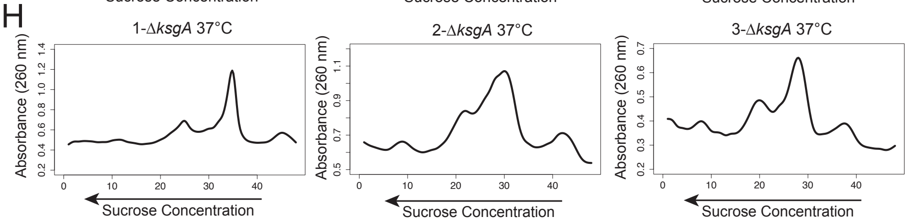


I

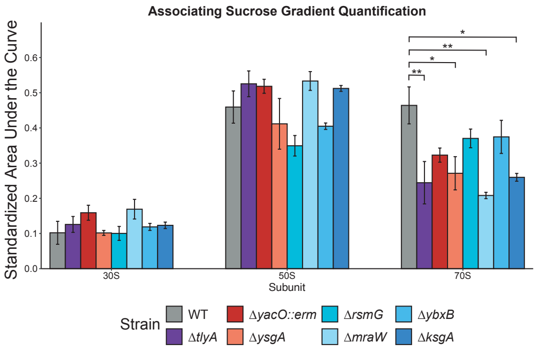


**Supplemental Figure S23 continued. Replicates of sucrose gradients of ribosome assembly at 37°C.** Shown are sucrose gradients to assess 50S and 30S subunit accumulation from the indicated deletion mutants with cells grown at 25°C under dissociating conditions. The samples are as follows: **G)** Δ*ybxB*, **H)** Δ*ksgA.* **I)** quantified area under the curve for 30S and 50S subunits. 50S were the same as wild type. We did detect differences in 30S accumulation with p<0.05* and p<0.01**.

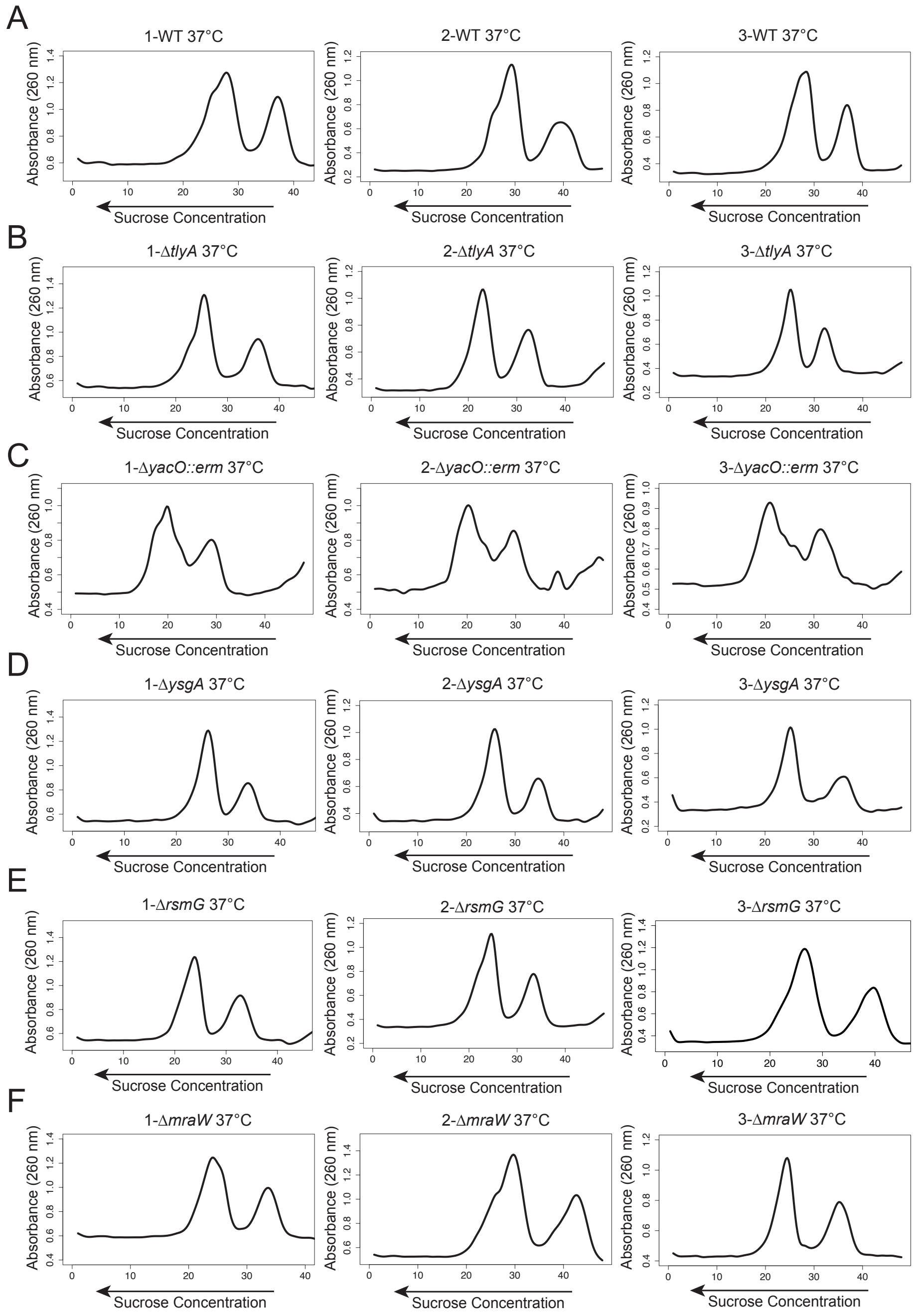


**Supplemental Figure S24. Replicates of sucrose gradients during dissociating conditions at 37ºC.** Shown are sucrose gradients to assess 50S and 30S subunit accumulation from the indicated deletion mutants with cells grown at 25ºC under dissociating conditions. The samples are as follows: **A)** WT, **B)** Δ*tlyA*, **C)** Δ*yacO::erm*, **D)** Δ*ysgA*, **E)** Δ*rsmG*, **F)** Δ*mraW*.

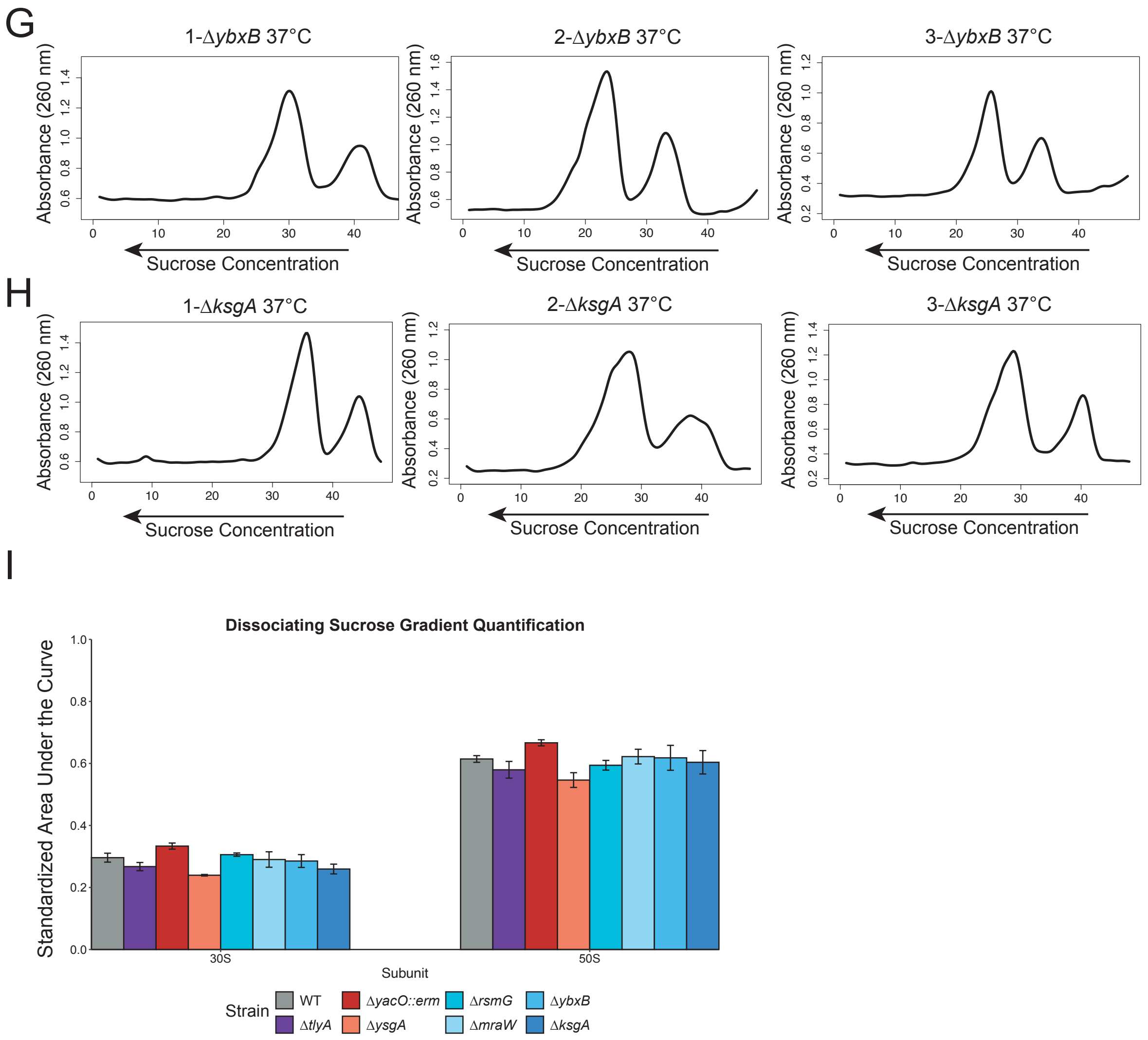


**Supplemental Figure S24 continued. Replicates of sucrose gradients during dissociating conditions at 37°C.** Shown are sucrose gradients to assess 50S and 30S subunit accumulation from the indicated deletion mutants with cells grown at 25°C under dissociating conditions. The samples are as follows: **G)** Δ*ybxB*, **H)** Δ*ksgA.* **I)** Quantified area under the curve for 30S and 50S subunits. 50S were the same as wild type.

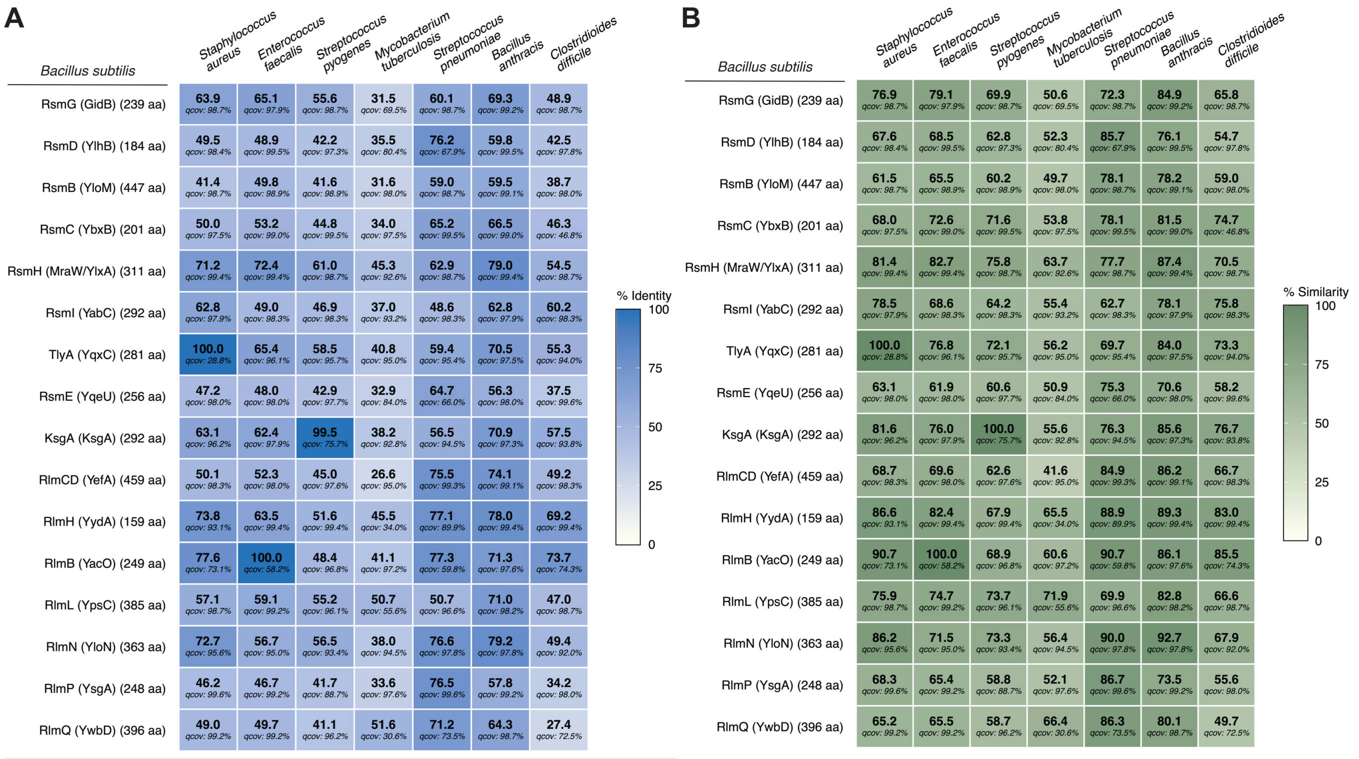


**Figure S25. rRNA MTases in *B. subtilis* are conserved in relevant bacterial pathogens.** Conservation heatmap of methyltransferase enzymes characterized in this study across common gram-positive pathogenic species. B. subtilis protein names and lengths listed on the left. Percent of exact **(A)** or similar **(B)** amino acid sequence matches along with percent query coverage (qcov) for BLAST searches are reported in each heatmap tile.
